# An unusual trafficking domain in MSRP6 defines a complex needed for Maurer’s clefts anchoring and maintenance in *P. falciparum* infected red blood cells

**DOI:** 10.1101/2021.12.03.471078

**Authors:** Alexandra Blancke Soares, Jan Stäcker, Svenja Schwald, Wieteke Hoijmakers, Nahla Galal Metwally, Jakob Cronshagen, Hanno Schoeler, Sven Flemming, Katharina Höhn, Ulrike Fröhlke, Paolo Mesén-Ramírez, Bärbel Bergmann, Melissa Khosh-Naucke, Iris Bruchhaus, Richárd Bártfai, Tobias Spielmann

## Abstract

Intracellular malaria blood stage parasites remodel their host cell, a process essential for parasite survival and a cause of pathology in malaria infections. Host cell remodeling depends on the export of different classes of exported parasite proteins into the infected red blood cell (RBC). Here we show that members of a recently discovered group of difficult to predict exported proteins harbor an N-terminal export domain, similar to other classes of exported proteins, indicating that this is a common theme among all classes of exported proteins. For one such protein, MSRP6 (MSP-7 related protein 6), we identified a second, untypical export-mediating domain that corresponded to its MSP7-like region. In addition to its function in export, this domain also mediated attachment to the Maurer’s clefts, prominent parasite-induced structures in the host cell where MSRP6 is located. Using BioID with the Maurer’s clefts attachment domain of MSRP6 to identify interactors and compartment neighbors in live parasites we discovered a novel complex of proteins at the Maurer’s clefts. We show that this complex is necessary for the anchoring and maintaining the structural integrity of the Maurer’s clefts. The Maurer’s clefts are believed to be involved in the transport of the major virulence factor PfEMP1 to the host cell surface where it mediates cytoadherence of infected RBCs to endothelial cells, a main reason for the importance of host cell modifications for parasite virulence in the human host. Taking advantage of MSRP6 complex mutants and IT4 parasites that we modified to express only one specific PfEMP1 we find that abolishing Maurer’s clefts anchoring was neither needed for PfEMP1 transport to the host cell surface nor for cytoadherence. Altogether, this work reveals parasite proteins involved in Maurer’s clefts anchoring and maintenance and unexpectedly finds that these functions seem to be dispensable for virulence factor transport and surface display.

## Introduction

The protozoan parasite *Plasmodium falciparum* causes the severest form of human malaria, a disease responsible for an estimated ∼600’000 deaths in 2022 (WHO, 2023). The pathology of malaria results from the propagation of *P. falciparum* parasites in the blood of the human host. In this phase the parasites develop within red blood cells (RBC) in a continuous cycle consisting of RBC invasion, intracellular growth, and production of daughter cells that are released under destruction of the host cell, leading to an exponential multiplication of parasites in the blood stream. The intracellular parasite exports a large number of different kinds of proteins (the ‘exportome’) into the host RBC (Spielmann and Gilberger, 2010). These exported proteins are secreted beyond the parasite’s plasma membrane (PPM) followed by translocation across the parasitophorous vacuolar membrane (PVM) that surrounds the parasite in the host cell (Beck et al., 2014; de Koning-Ward et al., 2009; Elsworth et al., 2014; Ho et al., 2018; Marti and Spielmann, 2013). The exported proteins alter the properties of the RBC to support the growth of the parasite in its intracellular niche (de Koning-Ward et al., 2016; Spillman et al., 2015). Exported proteins for instance mediate the cytoadherence of the infected RBC to the endothelium of blood vessels to avoid splenic clearance of the infected cell. The resulting sequestration of infected RBCs in organs of the host is a main reason for the virulence of P*. falciparum* parasites (Miller et al., 2002).

Knowing the parasite’s exportome, currently estimated to comprise ∼10% of the proteins encoded in the *P. falciparum* genome (Spielmann and Gilberger, 2015), is of critical importance to understand the molecular basis of the host cell modifications and their function. The prediction of the exportome is facilitated by a short amino acid motif, the PEXEL (*Plasmodium* export element) or HT (host targeting signal), a protease recognition sequence located downstream of a signal peptide (SP) that is found in many exported proteins (Boddey et al., 2010; Chang et al., 2008; Hiller et al., 2004; Marti et al., 2004; Russo et al., 2010; Sargeant et al., 2006). A second group comprises exported proteins that do not possess a PEXEL, the PEXEL negative exported proteins (PNEPs) and is thus missed by PEXEL-based predictions of the *P. falciparum* exportome (Spielmann&Gilberger2010; Spielmann et al., 2006). The initially discovered PNEPs contain no canonical SP but a transmembrane domain. The N-terminal region of these PNEPs and the mature N-terminus of PEXEL were found to be exchangeable in their property to promote export (Gruring et al., 2012). Based on these findings it was proposed that the mature N-terminus of exported proteins represents a unifying export domain (Gruring et al., 2012; Heiber et al., 2013). A different type of PNEP that contain an N-terminal SP were discovered more recently (Heiber et al., 2013; Kulzer et al., 2012). These proteins, here named SP-PNEPs, have so far not been extensively analysed and it is unclear if the N-terminal export domain indeed is a universal feature of all exported proteins of the parasite.

Many exported proteins are not essential for *in vitro* growth but likely are important for parasite survival and virulence in the host (Maier et al., 2008), a concept supported by work with orthologues in a rodent malaria model (De Niz et al., 2016). One such protein in *P. falciparum* is the main virulence factor PfEMP1 that is transported to the host cell surface where it is responsible for the cytoadherence of the infected RBC. Many other exported proteins indirectly contribute to cytoadherence, because they are needed for the transport and correct surface display of PfEMP1. Such proteins are SBP1 (Cooke et al., 2006; Maier et al., 2007), MAHRP1 (Spycher et al., 2008), REX1 (Dixon et al., 2011; McHugh et al., 2015), PTPs (Carmo et al., 2022; Maier et al., 2008; Rug et al., 2014), PHISTs (Oberli et al., 2016; Proellocks et al., 2014), GEXP07 (McHugh et al., 2020) and the co-chaperon PFA66(Diehl et al., 2021). However, how these proteins influence PfEMP1 transport is unclear. PfEMP1 is found in abundance at large disk-shaped vesicular structures in the host cell termed Maurer’s clefts (Hanssen et al., 2008b; Lanzer et al., 2006; Mundwiler-Pachlatko and Beck, 2013) that are installed by the parasite and are believed to function as hubs for proteins destined for the infected RBC surface (Wickham et al., 2001). PfEMP1 therefore likely traffics via these structures before being displayed on the host cell surface (Kriek et al., 2003; McMillan et al., 2013). Deletions of REX1, PTP1, MAHRP1, PTP7 or GEXP07 resulted in an altered architecture of the Maurer’s clefts, indicating that the correct organisation of these structures may be important for PfEMP1 transport (Carmo et al., 2022; Hanssen et al., 2008a; Maier et al., 2008; McHugh et al., 2020; Rug et al., 2014; Spycher et al., 2006). This is also supported by findings with RBCs from hemoglobinopathy patients that display a disturbed organisation of the Maurer’s clefts, which may cause reduced cytoadhesion and contribute to the protection from severe disease observed in these patients (Cyrklaff et al., 2011). However, other proteins needed for PfEMP1 trafficking, such as SBP1, do not cause gross changes of the Maurer’s clefts when deleted (Cooke et al., 2006; Maier et al., 2007; Maier et al., 2008).

In RBC containing early stage *P. falciparum* parasites, the Maurer’s clefts are mobile and become fixed in their position in the host cell during a narrow time window when parasites develop from the ring to the trophozoite stage (Gruring et al., 2011). The time of this Maurer’s clefts arrest coincides with the time PfEMP1 appears on the host cell surface and parasites become cytoadherent (Kriek et al., 2003; McMillan et al., 2013). Tether structures at the Maurer’s clefts containing the exported parasite protein MAHRP2 and host cell actin have both been implicated in the arrest of the Maurer’s clefts and this arrest was speculated to be important for PfEMP1 surface transport (Cyrklaff et al., 2011; Kilian et al., 2013; McMillan et al., 2013; Pachlatko et al., 2010; Rug et al., 2014). However, so far no parasite protein involved in the anchoring of the clefts has been identified and the importance of this phenomenon for PfEMP1 transport and cytoadherence is unclear.

Here we show that also in SP-PNEPs the mature N-terminus is an export domain, indicating that this is a unifying principle shared by all classes of exported proteins. Furthermore, we find an unusual, additional export domain in the SP-PNEP MSRP6. This export region also mediates Maurer’s clefts attachment and defines a Maurer’s clefts complex that appears in the middle of blood stage development when the majority of exported proteins have already been exported. Our results indicate that the MSRP6 complex mediates Maurer’s clefts anchoring and is needed to maintain the integrity of the Maurer’s clefts which disintegrate or aggregate in its absence. Unexpectedly, neither the failure to anchor the Maurer’s clefts negatively nor their disintegration affected virulence factor transport and cytoadhesion, indicating that Maurer’s clefts anchoring is not needed for PfEMP1 to reach the host cell surface.

## Results

### A shared export region in the N-terminal part of soluble PNEPs

PF08_0005 (PF3D7_0830300) and MSRP6 are two previously identified SP-PNEPs (Heiber et al., 2013). To identify the regions mediating export in this new type of PNEPs, we expressed GFP-tagged constructs with various deletions in *P. falciparum* parasites and analysed their export. For PF08_0005 we initially generated 3 deletions, one from amino acid (aa) 22-141 (PF08_0005Δ1, missing 120 aa immediately after the signal peptide), one from 142-266 (PF08_0005Δ2) and one missing the C-terminus (aa 267-388; PF08_0005Δ3) (Fig. 1A). While the deletion of part 2 and 3 had no noticeable effect on export, the protein missing part 1 could not be detected, suggesting that this deletion destabilised it. We therefore subdivided part 1 into 3 smaller parts [1a (25 aa), 1b (50 aa), and 1c (45 aa)] and tested the export of the proteins missing only these subparts. While part 1c was dispensable for export, PF08_0005 missing either part 1a or part 1b both showed a severe export defect (Fig. 1B). Deletion of part 1a led to an accumulation of the protein in the parasite periphery (typical for PV), whereas deletion of part 1b caused an accumulation around the nucleus (typical for ER) and in the PV. We conclude that the first 75 aa after the signal peptide of PF08_0005 are essential for the export of this protein. To test whether this region is also sufficient for export, we generated a minimal construct encompassing the signal peptide and parts 1a and 1b fused to GFP. This construct was efficiently exported, confirming this to be the region mediating export (Fig. 1C). When the SP with only part 1a was used, export with some additional accumulation of the reporter around the parasite was observed (Fig. 1C). These results indicated that the first 25 aa after the signal peptide are sufficient to mediate export but that either additional sequences from part 1b or a greater distance from the fusion tag (GFP) are required to obtain full export. Hence, in this SP-PNEP, the N-terminal region is necessary and sufficient for export.

**Figure 1.**
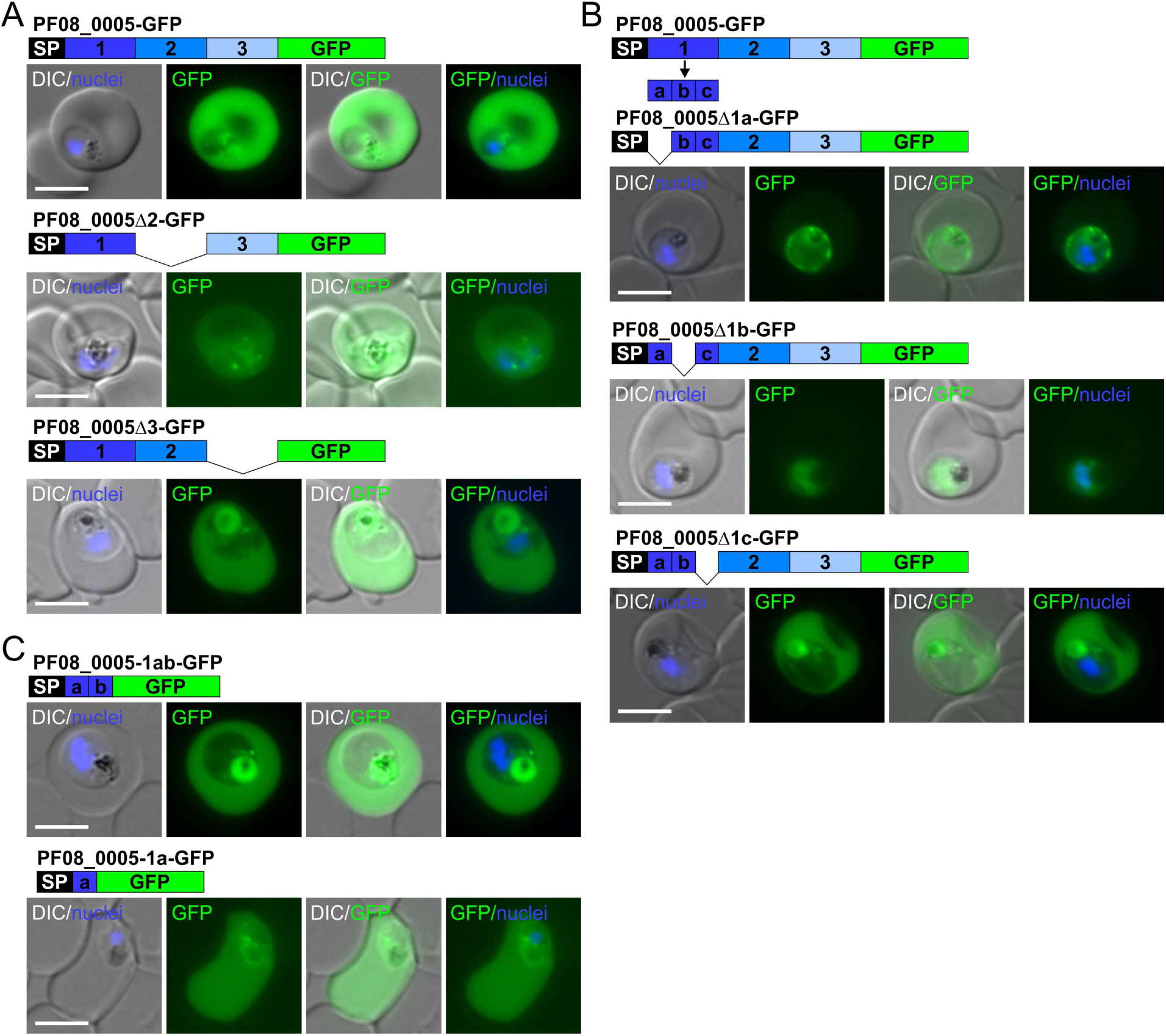
An N-terminal export domain in PF08_0005 (PF3D7_0830300). Representative live cell images of *P. falciparum* 3D7 iRBCs expressing episomal GFP fusion constructs under the *crt* promoter. Construct names and schematics are depicted above respective micrographs. Nuclei were stained with DAPI. Scale bar 5 µm. (A) The PF08_0005 sequence downstream of the signal peptide (SP) was divided into three parts. Shown are parasites expressing the full-length protein, and the protein lacking part 2 and 3, respectively. (B) PF08_0005 part 1 was further subdivided into three parts, a, b and c. Parasites expressing deletion constructs missing part a, b and c, respectively, are shown. (C) Parasites expressing minimal constructs containing only part a and b, or only part a.

To assess whether this is a common theme or not, we next, we carried out a similar deletion analysis for MSRP6, expressing GFP-tagged versions missing either the 30 amino acids (23-53) after the signal peptide (MSRP6Δ1), aa 54-323 (MSRP6Δ2), or 324-593 (MSRP6Δ3). Unexpectedly, none of these deletions prevented export (Fig. 2A). The only striking difference between these constructs was the observation that removal of part 3 abolished the Maurer’s clefts localisation and led to a soluble distribution in the host cell but did not affect export (Fig. 2A). We next replaced the signal peptide with that of the PVM protein ETRAMP10.1 (Spielmann et al., 2003) but this also had no influence on export of MSRP6 (Fig. 2A). Considering the possibility that multiple regions mediate export of MSRP6 and that in analogy to PF08_0005 (Fig. 1) and to Hsp70x (Kulzer et al., 2012) the N-terminus may be one of the export promoting regions, we generated a truncated construct of MSRP6 comprising the signal peptide, the following 102 aa and GFP. This construct was efficiently exported into the host cell (Fig. 2B). However, a shorter construct containing the SP and only the 31 following aa was not exported (Fig. 2B). Hence, the 102 amino acids of the mature MSRP6 are sufficient to mediate export but are not necessary for it in the full length MSRP6, possibly because of additional regions positively influencing export.

**Figure 2.**
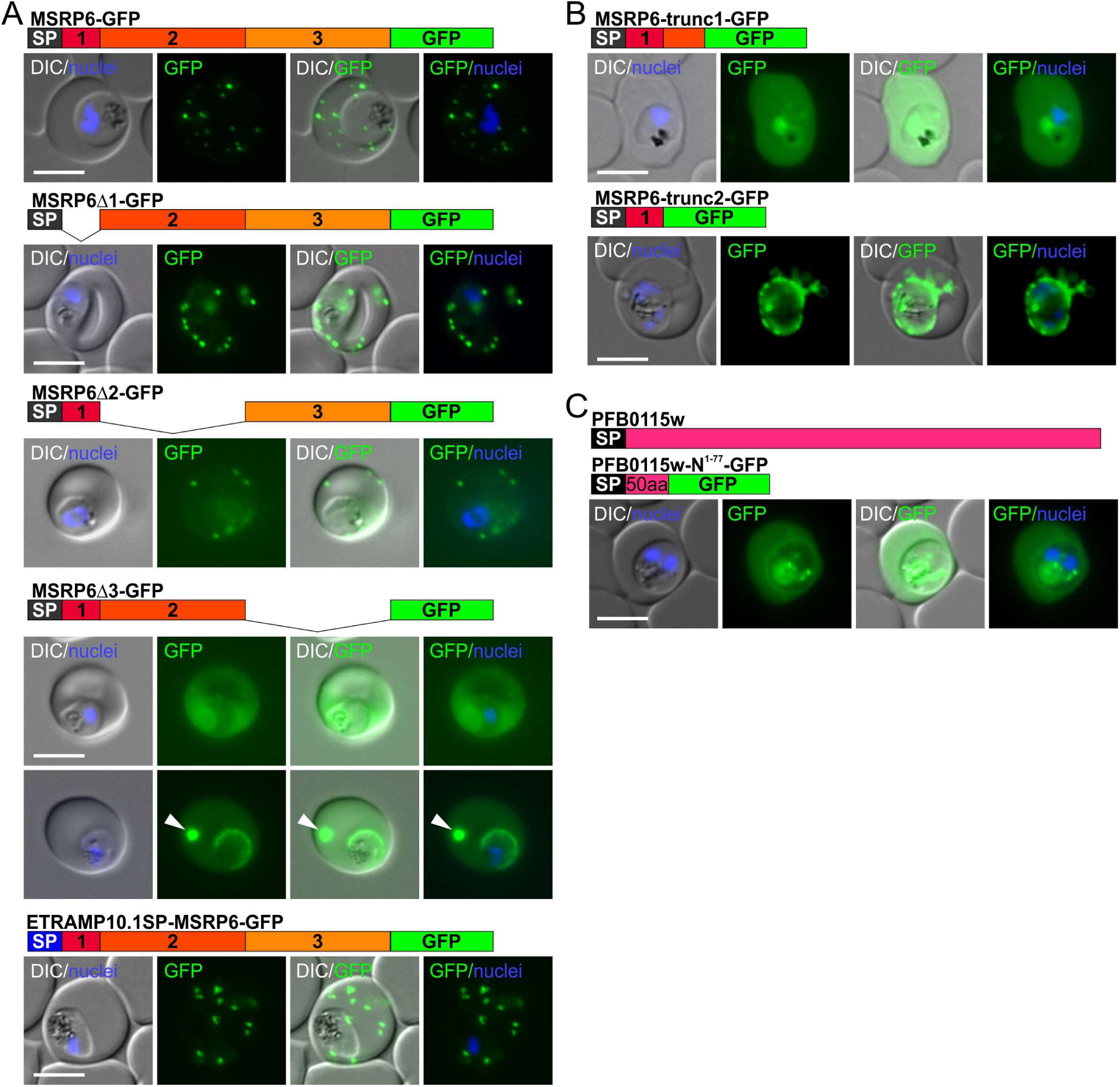
Export domains in MSRP6 and PFB0115w. Representative live cell images of *P. falciparum* 3D7 iRBCs expressing episomal GFP fusion constructs under the *crt* promoter. Construct names and schematics are depicted above the respective micrographs. Nuclei were stained with DAPI. Scale bar 5 µm. (A) MSRP6 downstream of the signal peptide (SP) was divided into three parts (1-3). Shown are parasites expressing the indicated constructs. The SP replacement was done with the SP from ETRAMP10.1. Arrowheads indicate a large focus of GFP fluorescence likely representing an overexpression induced aggregate. (B) Parasites expressing minimal constructs of MSRP6 containing part 1 and a fraction of part 2 or only part 1. (C) Parasites expressing minimal construct of PFB0115w containing the SP and downstream 50 aa.

As the N-terminus seemed to be a general export domain in SP-PNEPs, we tested in a recently discovered PNEP of this type (PFB0115c (PF3D7_0202400), (Birnbaum et al., 2017) now named (Chan et al., 2017)), whether the N-terminus was sufficient to mediate export. The corresponding construct, containing the SP and the subsequent 50 aa, was efficiently exported (Fig. 2C). We conclude that all PNEPs of this type analysed so far contain an export domain in the mature N-terminus, similar to other exported proteins (Gruring et al., 2012; Heiber et al., 2013), indicating that this is a general feature of all classes of exported proteins.

### MSRP6 possesses an unusual C-terminal export region that also mediates Maurer’s clefts localization

Our deletion constructs indicated that part3 of MSRP6, the region encompassing the MSP7-like domain, mediates Maurer’s clefts association (Fig. 2A). To narrow down the sequence responsible for this phenotype, we subdivided part 3 into four parts, 3a (aa 324-390), 3b (aa 391-447), 3c (aa 448-515) and 3d (aa 516-593) (Fig. 3A). We then expressed GFP-tagged MSRP6 missing overlapping regions, either 3ab, 3bc or 3cd, respectively. Deletion of 3ab had no effect on the Maurer’s clefts association of MSRP6, but both, MSRP6 missing 3bc and 3cd showed a soluble pool in the host cell (Fig 3B). An additional frequently observed focus probably represents an aggregate of the GFP-fusion protein. We conclude that the Maurer’s clefts binding domain must be contained in region 3cd. Indeed, when we appended 3cd to the soluble exported protein REX3 (Spielmann et al., 2006), the resulting protein (REX3cd-GFP) was now found at the Maurer’s clefts (Fig. 3C). To test the extent of the region required for Maurer’s clefts association, we generated REX3 versions harboring only 3c or 3d (REX3c-GFP and REX3d-GFP). However, neither of the regions alone mediated efficient Maurer’s clefts localisation (Fig. 3C). We conclude that both, region 3c and 3d, are necessary for Maurer’s clefts recruitment. Interestingly, part 3cd, corresponded to the region containing the MSP-7 like domain of MSRP6 (Fig. 3A).

**Figure 3.**
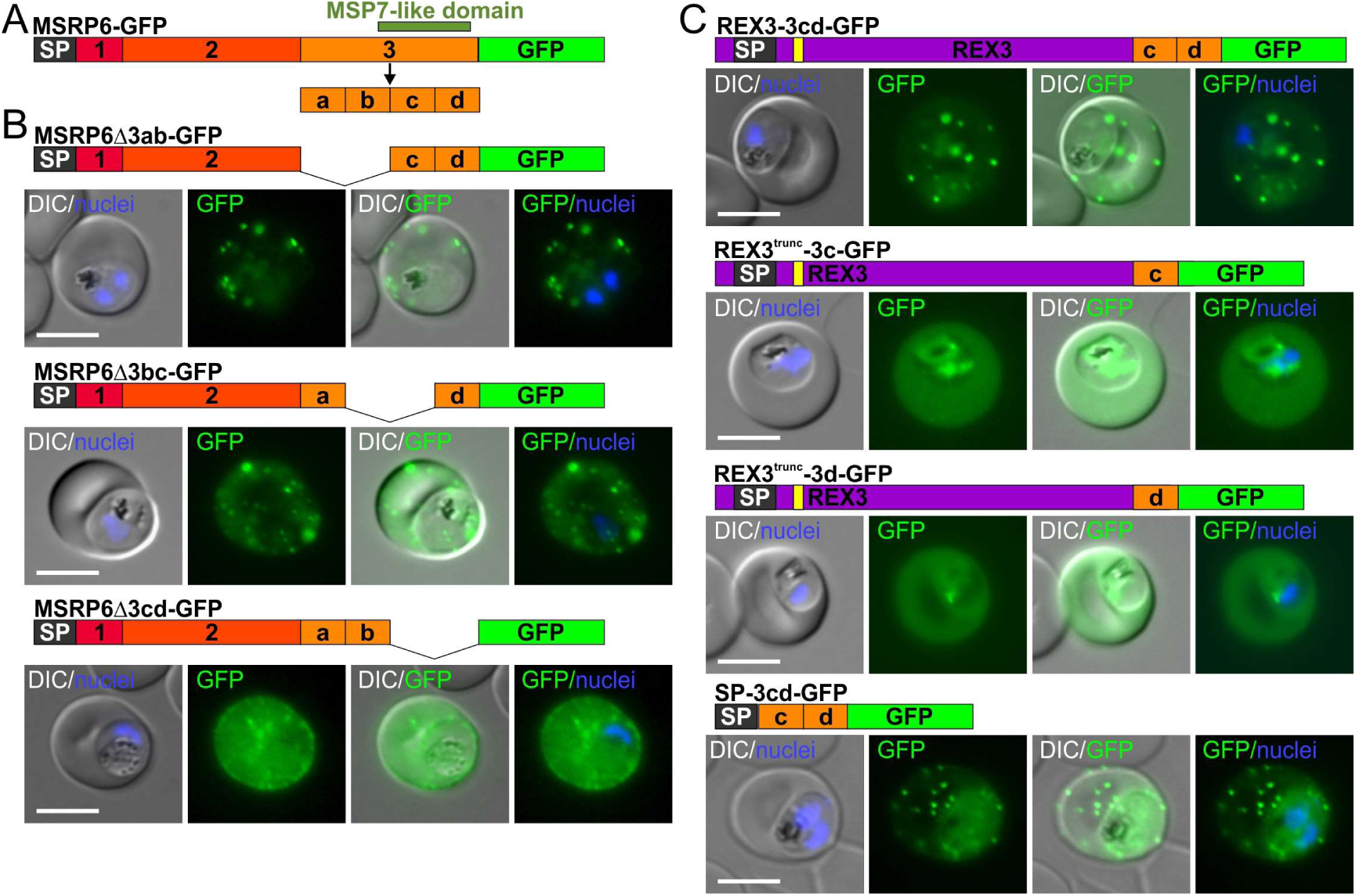
The MSP-7-related region in MSRP6 is sufficient for export and Maurer’s clefts attachment. Representative live cell images of *P. falciparum* 3d7 iRBCs expressing episomal GFP fusion constructs under the *crt* promoter. Construct names and schematics are depicted above respective micrographs. Yellow bar, PEXEL. Nuclei were stained with DAPI. Scale bar 5 µm. (A) Schematic showing subdivision of MSRP6 part 3 into 4 parts, a, b, c, and d. (B) Parasites expressing MSRP6 constructs with the indicated deletions. (C) Parasites expressing REX3 containing MSRP6 part c, d or c+d at its C-terminus or only the MSRP6 SP fused to parts c and d.

As the 3cd domain might mediate Maurer’s clefts attachment via protein-protein interactions, we reasoned that it could also contribute to the trafficking of MSRP6, for instance by a piggy back mechanism to another exported protein. We therefore tested whether 3cd alone can promote export by appending this region after the MSRP6 SP and expressing it as a GFP fusion (SP-3cd-GFP). This construct was exported and localised to the Maurer’s clefts (Fig. 3C). We conclude that MSRP6 contains two regions that are sufficient to mediate export, one at the mature N-terminus, similar to other exported proteins, and one that at the same time also mediates Maurer’s clefts association. The 3cd region of MSRP6 is therefore henceforth termed MAD (MSRP6 attachment domain).

### BioID identifies potential MSRP6 interaction partners

MSRP6 is found in electron dense, cloudy structures at the Maurer’s clefts (Heiber et al., 2013). As the MAD of this protein was sufficient and necessary for the Maurer’s clefts association, we used BioID (Kimmel et al., 2021; Roux et al., 2012) to identify interactors of this domain (Fig. 4). To this end we added BirA* to a construct similar to REX3 containing the MAD (REX3-3cd-GFP, Fig. 3C), except that REX3 was truncated to encompass the export domain only (REX3^trunc^-MAD-GFP-BirA*) to offer fewer MAD-unrelated interaction interfaces (Fig. 4A). This construct had the purpose to tag interaction partners of the MAD (in effect the MSP-7-like region in MSRP6) in growing parasites through the promiscuous biotinylation activity of BirA*. As a control for soluble proteins and hits specific for the REX3 part, we used the identical construct without the MAD (REX3^trunc^-GFP-BirA*). To distinguish the MAD-specific Maurer’s clefts proteome from other proteins present at this compartment, we used a third construct consisting of an N-terminal fragment of STEVOR to mediate trafficking to the Maurer’s clefts (Przyborski et al., 2005) fused to GFP and BirA* (Fig. 4A).

**Figure 4:**
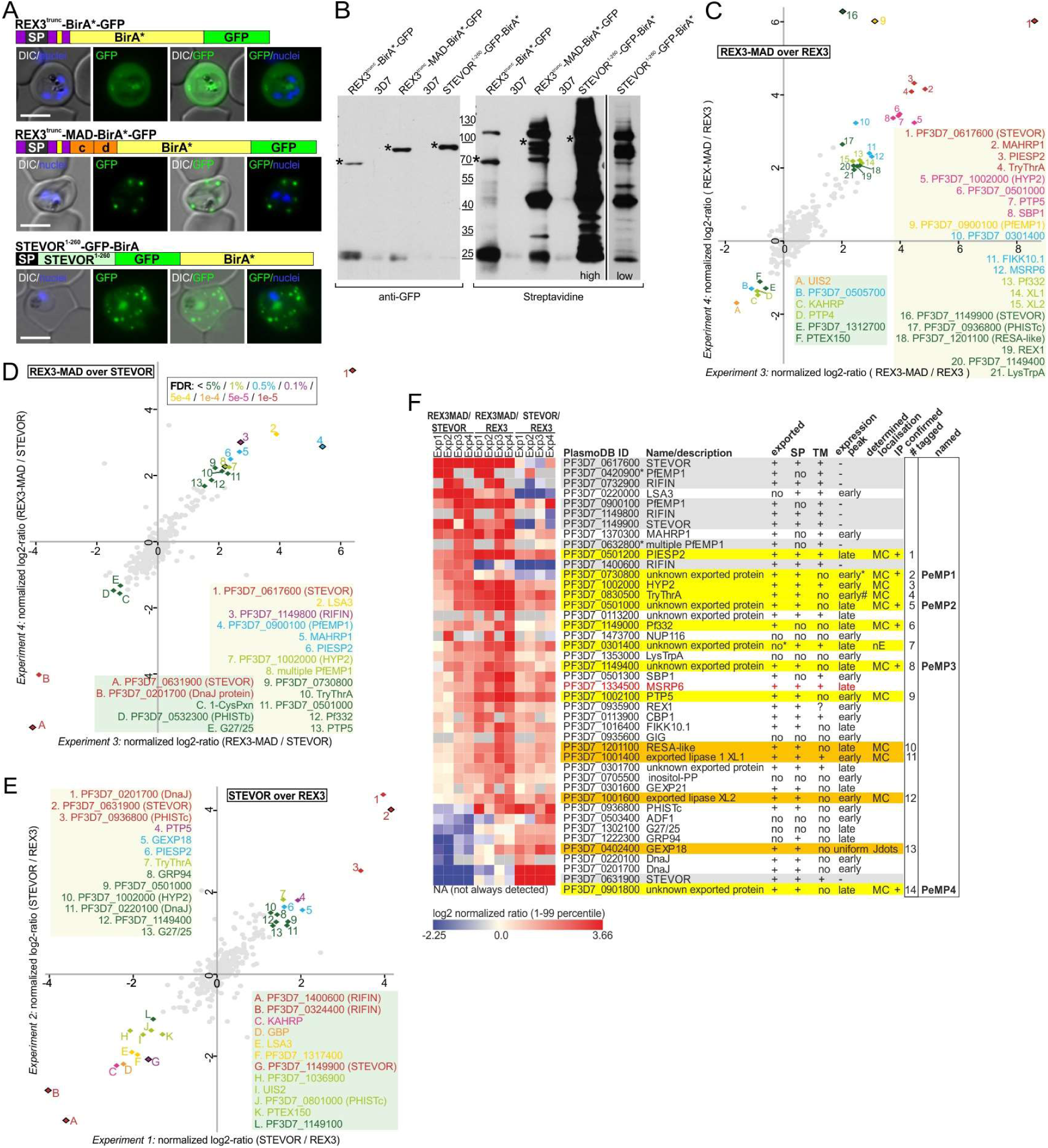
Bio-ID of Maurer’s cleft proteome and MSRP6 MAD interactome. (A) Live cell images of *P. falciparum* 3D7 infected RBCs expressing the episomal BirA*-GFP fusion constructs schematically shown above the panels. Yellow bar, PEXEL (top, N-terminal 70 aa of REX3 (REX3trunc) fused to BirA* and GFP (host cell soluble control), middle, REX3trunc fused to MSRP6 parts c and d, BirA* and GFP (MAD-specific), bottom, N-terminal 260 aa of STEVOR fused to GFP and BirA* (general Maurer’s clefts control)). Nuclei were stained with DAPI. Scale bar 5 µm. (B) Western blots with parasite lysates of the 3 cell lines in (A) and 3D7 control probed with α-GFP (left) and Streptavidin (right). Two different exposures of the strepavidin probed blot are shown for STEVOR^1-260^-GFP-BirA* (low and high) for comparability to the REX3 samples. Asterisks, bands representing the respective fusion proteins. (C-E) Scatterplots of quantitative Bio-ID experiments (see Fig. S1 for all plots of all experiments and Table S1 for full data) identifying proteins significantly enriched or depleted in the MAD interactome compared to the host cell cytosolic proteome (C) or Maurer’s cleft proteome (D) or in the Maurer’s cleft proteome compared to the host cell cytosolic proteome (E). Normalized ratios were calculated for proteins identified by at least two peptides and normalized log2-ratios of replicate experiments were plotted. Intensity-based outlier statistics (two-sided Benjamini-Hochberg test) was applied to calculate FDR values and proteins enriched or depleted with an FDR below 5% in both replicates were labelled with a color-code reflecting the level of significance in the least significant experiment (<5% dark green, <1% light green, <0.5% blue, <0.1% purple, <5e^-4^ yellow, <1e^-4^ orange, <5e^-5^ pink, <1e^-5^ red). Significant hits are numbered and gene-IDs or short unique names are given. Proteins encoded by multigene families (STEVORs, RIFINs, PfEMP1) that might differ in expression between the distinct cell lines and hence may show as false-positives, are marked with rhomboids with black frame. (F) A heatmap representation of all replicates each of REX3-MAD-over-STEVOR (MAD over unrelated Maurer’s cleft proteome), REX3-MAD-over-REX3 (MAD over host cell cytosolic proteome) and STEVOR-over-REX3 (Maurer’s cleft proteome over host cell cytosolic proteome) quantitative Bio-ID experiments (C-E and Fig. S3, see Table S1 for full data) for proteins enriched with an FDR <5% in at least 2 out of 4 replicate reactions. Proteins are ranked from high to low on the average normalized log2 ratio of the REX3-MAD-over-STEVOR comparison and color intensity portrays the normalized log2 ratio per experiment (red enriched, yellow neutral, blue depleted based on the 1-99 percentile of values for all identified proteins). Grey blocks are proteins not identified in an experiment or for which no ratio could be calculated due to a missing label. PlasmoDB gene identifiers, short names and protein product descriptions are listed. Yellow labelled proteins were selected for follow-up as putative MAD interactors, orange labelled proteins showed moderate to no enrichment as MAD interactors but were enrichment in the Maurer’s cleft proteome. Proteins encoded by clonally variant multigene families (likely false-positives) are labelled grey. The heatmap was generated using the web-based Morpheus tool from the Broad Institute (Harvard, 2017).

Expression of these constructs in the parasite resulted in the expected localisation: REX3^trunc^-MAD-BirA*-GFP and the STEVOR construct were found at the Maurer’s clefts whereas REX3^trunc^-BirA*-GFP was evenly distributed in the host cell cytoplasm (Fig. 4A). In each of the 3 cell lines expressing one of these constructs, addition of biotin led to biotinylation of multiple proteins, as evident by streptavidin Western blots (Fig. 4B). No biotinylation was observed in 3D7 parasites, suggesting that the trace amounts of biotin in the culture medium were in this case not sufficient to cause detectable protein biotinylation. We then purified the biotinylated protein from the 3 cell lines using streptavidin beads and carried out quantitative mass spectrometry using dimethyl labelling (Boersema et al., 2009). This was done in two independent experiments, each split into 2 technical replicas (experiment 1-4, Table S1). To remove REX3 specific hits and the background of soluble host cell proteins, we first compared REX3^trunc^-MAD-BirA*-GFP with REX3^trunc^-BirA*-GFP (Fig. 4C and Fig. S1). Using a false discovery rate (FDR) of ≤0.05 as cut off we found 30 hits enriched in the construct containing the MAD over the REX3^trunc^ only control of which 22 were present in at least 3 of the 4 experiments (Fig. 4C, Fig. S1 and Table S1). Of these 22, 20 were annotated or known to be exported (according to PlasmoDBv33), suggesting efficient enrichment of relevant proteins (Table S1). Next we compared the MAD-specific sample (REX3^trunc^-MAD-BirA*-GFP) with the general Maurer’s clefts proteome generated with the truncated STEVOR (STEVOR-BirA*-GFP). Twenty seven hits were found to be enriched (with a FDR ≤ 0.1; FDR cut-off was adapted in different experiments based on the different distribution and shift in the background cloud) in the MSRP6 MAD-specific Maurer’s clefts proteome compared to the STEVOR-derived Maurer’ clefts proteome in one or both replicas (Fig. 4D, Fig. S1 and Table S1). Of these, 26 are known or annotated to be exported (PlasmodBv33). The 5 top hits were replicated in both independent experiments with an FDR ≤0.01 (Fig. 4D, Fig. S1 and Table S1). Several of these hits were also overrepresented in STEVOR-BirA*-GFP over REX3^trunc^-MAD-BirA*-GFP (Fig. 4E, Fig. S1 and Table S1), as expected if the truncated STEVOR enriches for general Maurer’s clefts proteins. Altogether we found 20 hits that were overrepresented (FDR ≤ 0.1) in the MAD-specific Maurer’s clefts proteome over both, the general STEVOR-derived Maurer’s clefts proteome and the REX3 soluble host cell control (Fig. 4F, Table S1). A summary of the significant hits in at least 2 replicas for any of the constructs is shown in Fig. 4F. Enrichment of members of clonally-variant protein families (e.g. PfEMP1s, STEVORs, RIFINs) were not further considered, as these likely were due to differential expression of family members in the three cell lines.

### General and MSRP6-MAD specific BioID hits are Maurer’s clefts proteins

To validate the MS results we endogenously GFP-tagged (by modification of the genomic locus using SLI (Birnbaum et al., 2017), Fig. S2) 9 candidates that were significantly enriched in at least two experiments in both, MAD over REX3 and MAD over STEVOR (PIESP2, PF3D7_1002000/PF10_0024, PF3D7_0830500/PF08_0003, PF3D7_0501000/PFE0050w, Pf332, PF3D7_0301400/PFC0070c, PF3D7_1149400/PF11_0511, PTP5), and one candidate significantly enriched in MAD over STEVOR only (PF3D7_0730800/Mal7P1.170). These candidates are putative MSRP6-specific Maurer’s clefts proteins (Fig. 5A). We also tagged 3 candidates significantly enriched in MAD over REX3 only (PF3D7_1201100/PFL0055c, XL1, XL2), one that was enriched in STEVOR over REX3 only (GEXP18) and one that was enriched in MAD over REX3 and STEVOR over REX3, but that was not detected in all experiments (PF3D7_0901800/PFI0086w) and hence is not included in the heat map in Figure 4F. These candidates comprise putative non-MSRP6-specific Maurer’s clefts proteins (Fig. 5B). Thirteen of the 14 tested proteins from both groups were found to be exported by this approach of endogenously tagging the protein with GFP (Fig. 5A, B, Fig. S2). The non-exported protein was PF3D7_0301400/PFC0070c (present in foci with an additional soluble pool in the parasite). This protein contains a C-terminal transmembrane domain, a feature that can prevent export when the protein is C-terminally tagged with GFP (Heiber et al., 2013). As it also contains a PEXEL, it may therefore nevertheless still be an exported protein that here appeared non-exported due to the tag. Of the 13 exported proteins, 12 showed a Maurer’s clefts typical pattern, with some showing an additional soluble pool in the host cell (Fig. 5A, B). Some of the proteins were only exported or recruited to the Maurer’s clefts in later stage parasites, as younger stages showed no export or no Maurer’s clefts association (Fig. 5C). The Maurer’s clefts localisation was confirmed for 9 candidates by co-localisation with a Maurer’s clefts marker (Fig. S2). Of note, PIESP2-GFP cells usually showed more foci than typical for Maurer’s clefts but this was matched by the Maurer’s clefts marker, suggesting a Maurer’s clefts phenotype due to GFP-tagging this protein (Fig. S2A and see below). One protein (GEXP18), a non-MSRP6-specific hit, showed a punctate pattern that did only partially overlap with Maurer’s clefts and showed more foci than typical for Maurer’s clefts (Fig. 5B and Fig. S2B) in agreement with an episomally expressed version of this protein previously localised to the J dots (Zhang et al., 2017). Overall, 12 out of the 14 tested proteins are therefore likely Maurer’s clefts proteins, validating our approach.

**Figure 5.**
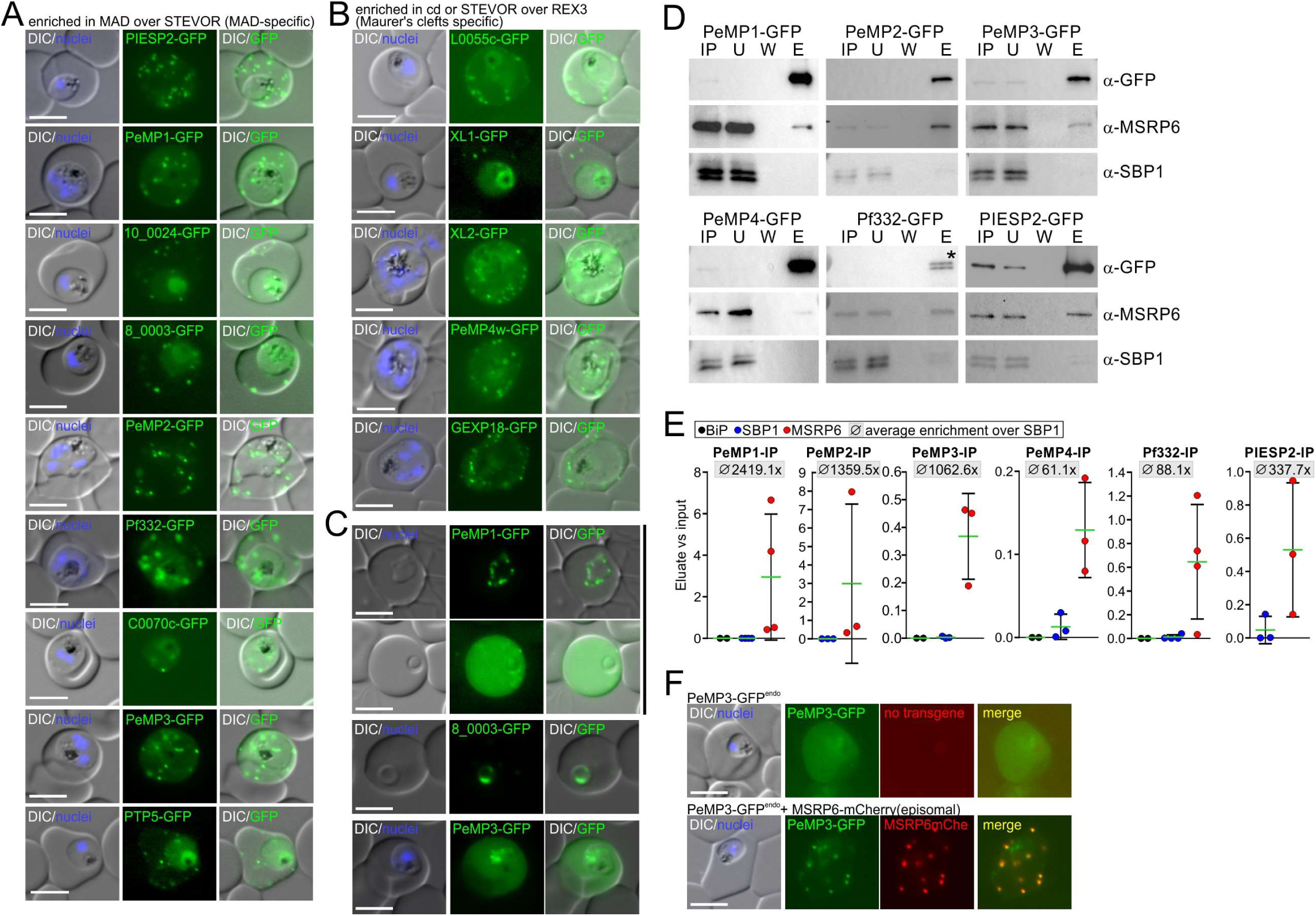
Validation of Bio-ID hits. (A,B) Representative live cell images of *P. falciparum* infected RBCs expressing endogenously GFP-tagged (using the standard 2xFKBP-GFP tag (Birnbaum et al., 2017)) proteins enriched in REX3^trunc^-MAD-over STEVOR^1-260^-STEVOR-GFP-BirA*in BioID experiments (Fig. 4), representing potential MAD specific interactiors (A) or enriched in MAD or STEVOR over REX3^trunc^-GFP-BirA* (Fig. 4), representing Maurer’s cleft specific proteins (B). (C) Ring stage parasites showing stage specific localisations of PeMP1, PF08_0003 and PeMP3. (D) Co-IP experiments showing interaction of MSRP6 with the pulled down endogenously GFP-tagged bait protein (indicated above the blot) and probed with the indicated antibodies. IP, total input lysate; U, unbound protein of the lysate after IP; W, fifth wash; E, eluate. Asterisk shows Pf332 degradation product, this protein is too large to detect full length on a standard PAGE. All replicas and full blots shown in Fig. S6. (E) Quantification of IP experiments in (D) and Fig. S6, see Table S2 for exact values. Error bars show SD, green line the mean. Average of enrichment of MSRP6 over SBP1 is indicated above each graph. (F) Representative live cell images of *P. falciparum* young trophozoite iRBCs expressing endogenously GFP-tagged PeMP3 with (bottom) and without episomally expressed MSRP6-mCherry. If no simple short name was known (Fig. 4F) the protein designations in the imaging panels were 10_0024-GFP for PF3D7_1002000/PF10_0024, 8_0003-GFP for PF3D7_0830500/PF08_0003, C0070c for PF3D7_0301400/PFC0070c and L0055c for PF3D7_1201100/PFL0055c. A, B, and F, Nuclei were stained with DAPI; merge, overlay of red and green channel; Scale bar 5 µm.

Remarkably, 9 of the 12 Maurer’s clefts localising proteins did not contain a transmembrane domain (Fig. 4F), indicating that many of the putative MAD interaction partners are peripherally attached to the Maurer’s clefts, similar to MSRP6 itself. We therefore named these proteins ‘Peripheral Maurer’s clefts protein’ (PeMP), except for those that already had a name or a different defining feature (Fig. 4F). To confirm interaction with MSRP6, we carried out Co-IPs with PIESP2, Pf332, PeMP1, PeMP2, PeMP3, and PeMP4. All of these proteins co-immunoprecipitated MSRP6 (Fig. 5D, Fig. S3). As a stringent control we used SBP1, a Maurer’s clefts protein present at these structures throughout the asexual cycle (Gruring et al., 2012) and suggested to be part of multiple complexes (Rug et al., 2014; Takano et al., 2019). As in some IPs small amounts of SBP1 also co-precipitated, we included BiP as a control in some experiments. Only negligible amounts of BiP co-precipitated (Fig. 5E, Fig. S5). This indicated that very small amounts of SBP1 were also co-purified on some occasions. However, MSRP6 was manifold more enriched than SBP1 in all replicas with an average of >60 fold up to >2000 fold enrichment (depending on the candidate) over SBP1 (Fig. 5D,E and Fig. S3, Table S2), suggesting the tested BioID candidates were specific interactors in a complex with MSRP6. In contrast, SBP1 may either be a very weak or indirect interactor or may have been detected due to low level immunoprecipitation of cleft fragments. Overall, these findings indicated the existence of an MSRP6-specific complex. In the case of PeMP3 the interaction with MSRP6 also appeared to promote Maurer’s clefts association. This was indicated by experiments showing that overexpression of MSRP6 under an earlier promoter led to efficient recruitment of PeMP3 to the Maurer’s clefts in younger (one nucleus) trophozoites in all cells that harbored the episomally expressed MSRP6, but was absent from the clefts in cells that lacked the overexpressed MSRP6-mCherry (Fig. 5F, Fig. S4). This suggested a direct dependency on MSRP6 for PeMP3 Maurer’s clefts recruitment.

We conclude that the MSRP6-MAD interactome identifies a complex or interacting network of Maurer’s clefts proteins of which many lacked a transmembrane domain. Interestingly, in contrast to the majority of known exported proteins (Marti et al., 2004), most of the MAD-specific hits were either expressed at later stages of the intraerythrocytic cycle (MSRP6 itself, PIESP2, Pf332, PeMP2, PeMP3, and PeMP4) (Fig. 4F) or if not, they were late exported (PF3D7_0830500, Fig. 5C and (Heiber et al., 2013)) or recruited to the Maurer’s clefts only in late stages (PeMP1, PeMP3, Fig. 5C), matching the expression profile of MSRP6. As many of the early expressed exported proteins, such as SBP1, remain at the Maurer’s clefts throughout the cycle and therefore are still present at the stage the MSRP6 complex proteins reach the clefts, this suggests that the MSRP6 complex is a late stage Maurer’s clefts complex that is independent of the proteins already residing at the Maurer’s clefts (Table S1).

### MSRP6 complex proteins are required for Maurer’s clefts anchoring and maintenance in late stages

To assess the function of MSRP6 complex proteins, we generated disruptions of 6 of the confirmed MSRP6 interactors PeMP1, PeMP2, PeMP3, PeMP4, PIESP2 and Pf332 (Fig. S5A) using SLI-TGD (Birnbaum et al., 2017) and assessed the state of the Maurer’s clefts using episomally expressed SBP1-mScarlet as marker. No difference in the number of clefts per cell was observed in the TGDs of PeMP1, PeMP2, PeMP3, and PeMP4 when compared to endogenously GFP-tagged PeMP2-GFP used as control (Fig. 6A) and the number of clefts was similar to that reported for wild type parasites in previous work (Gruring et al., 2011; Hanssen et al., 2008a). In the parasites with a disruption of Pf332 we observed cells with one to few large and intense foci of SBP1-mScarlet instead of the typical multiple foci as well as cells with the large focus and additional, small foci (Fig. 6B, D). For the PIESP2 disruption we observed cells with more SBP1 foci than typical, similar to the line with the GFP-tagged PIESP2 (Fig. 6B, C; Fig. 5A, FigS2A). While the PeMP2-TGD parasites had similar numbers of Maurer’s clefts compared to controls, we noticed that in some cells (43%, n=42) Maurer’s clefts were present in pairs and in later stages were frequently in aggregates that were present in addition to unaggregated clefts (Fig. 6B). Furthermore, in trophozoites of the Pf332-TGD, PIESP2-TGD and PeMP2-TGD parasites, the Maurer’s clefts still appeared to be mobile (Fig. 6D). At this stage the Maurer’s clefts in wild type parasites are arrested (Gruring et al., 2011), indicating an impairment of the anchoring of the Maurer’s cleft in these three mutants. Disruptions of PeMP1, PeMP3 and PeMP4 did not impair anchoring of the Maurer’s clefts (Fig. S5B). As the PIESP2 phenotype in part deviated from that in previous work (no Maurer’s clefts phenotype using SBP1 as a marker (Maier et al., 2008) or a smaller size of the Maurer’s clefts (Zhang et al., 2018)), we complemented the PIESP2-TGD. We used an untagged PIESP2 for this since tagging the endogenously expressed PIESP2 already caused an effect similar to that of the disruption (Fig. 5A and Fig S2A). In the complemented parasites, the number of Maurer’s clefts was reverted to that typical for wt parasites (evident from PeMP2-GFP^endo^ parasites used as control here) (Fig. 6C) and Maurer’s clefts anchoring was restored (Fig. 6E), confirming that these effects were specific for PIESP2 disruption and not the result of unrelated alterations in this cell line.

**Figure 6.**
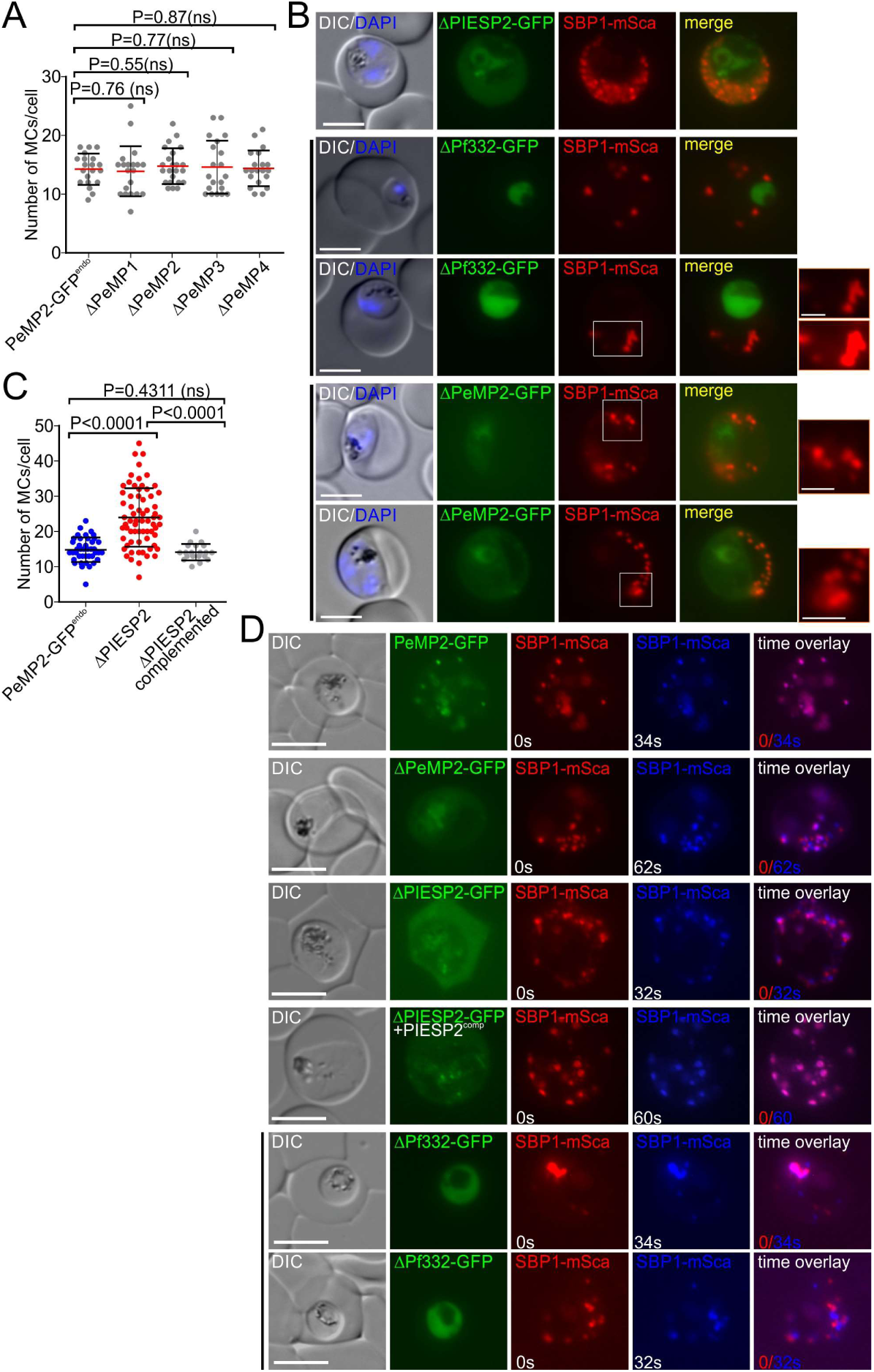
Maurer’s clefts numbers and anchoring in MSRP6 complex member mutants. (A) Quantification of the number of Maurer’s clefts (MCs) per iRBC in the indicated cell lines (Δ indicates disrupted gene of the indicated protein). Each dot represents the Maurer’s clefts number in a single cell. n = 20 cells for each cell line except ΔPeMP2 (23 cells). (B) Live cell images of ΔPIESP2-GFP (top panel), ΔPf332-GFP (panel 2 and 3) or ΔPeMP2-GFP (panel 4 and 5) parasites, respectively, co-expressing episomal SBP1-mScarlet as a Maurer’s clefts marker. Boxes, areas enlarged on the right. Nuclei stained with DAPI. (C) Quantification of the number of Maurer’s clefts per infected RBC in the indicated cell lines as in (A). n = 41 cells for the PeMP2-GFP parasites, 67 cells for the ΔPIESP2 parasites and 20 cells for the ΔPIESP2-complemented parasites, respectively. (D) Short-term time-lapse images of PeMP2-GFP (top panel), ΔPeMP2-GFP (panel 2), ΔPIESP2-GFP (panel 3) or ΔPf332-GFP (panel 4 and 5) parasites, respectively, co-expressing episomal SBP1-mScarlet. Images of the same cell were taken at the time points indicated. Images from the second time point were converted to blue and overlaid with the first time point to visualise MC movement (purple, overlay of the SBP1-signals indicating arrest of MCs, signal with original colour indicative of MCs movement). (B, D), scale bars, 5 μm and 2 µm for enlarged areas. (A, C), mean and error bars (SD) are shown; two-tailed unpaired t-test, P-values indicated. DIC, differential interference contrast; ns, not significant.

As anchoring of the Maurer’s clefts had not previously been analysed and the published phenotypes only partially matched those of our PIESP2- and Pf332-TGD lines, we carried out a detailed analysis of these two mutants. The PIESP2-TGD phenotype was stage-specific. In young trophozoites the Maurer’s clefts number per cell was comparable to controls, whereas in later stages the number increased with parasite age (Fig. 7A). This indicated that the Maurer’s clefts fragmented in later stage parasites which was confirmed by electron microscopy (EM) analysis. Already tagging of PIESP (PIESP2-GFP^endo^) resulted in a decreased Maurer’s cleft length in late trophozoite parasites compared to controls, while the number of clefts per cell section was increased (Fig. 7B-D). In the PIESP2-TGD line, the cleft length was even shorter but by EM the number of clefts appeared lower than in controls (Fig. 7B-D). This was likely because the clefts became too small for detection by that method, an interpretation supported by the live cell images which showed many small foci and a blurred background with the cleft marker SBP1 (Fig. 6B, C).

**Figure 7.**
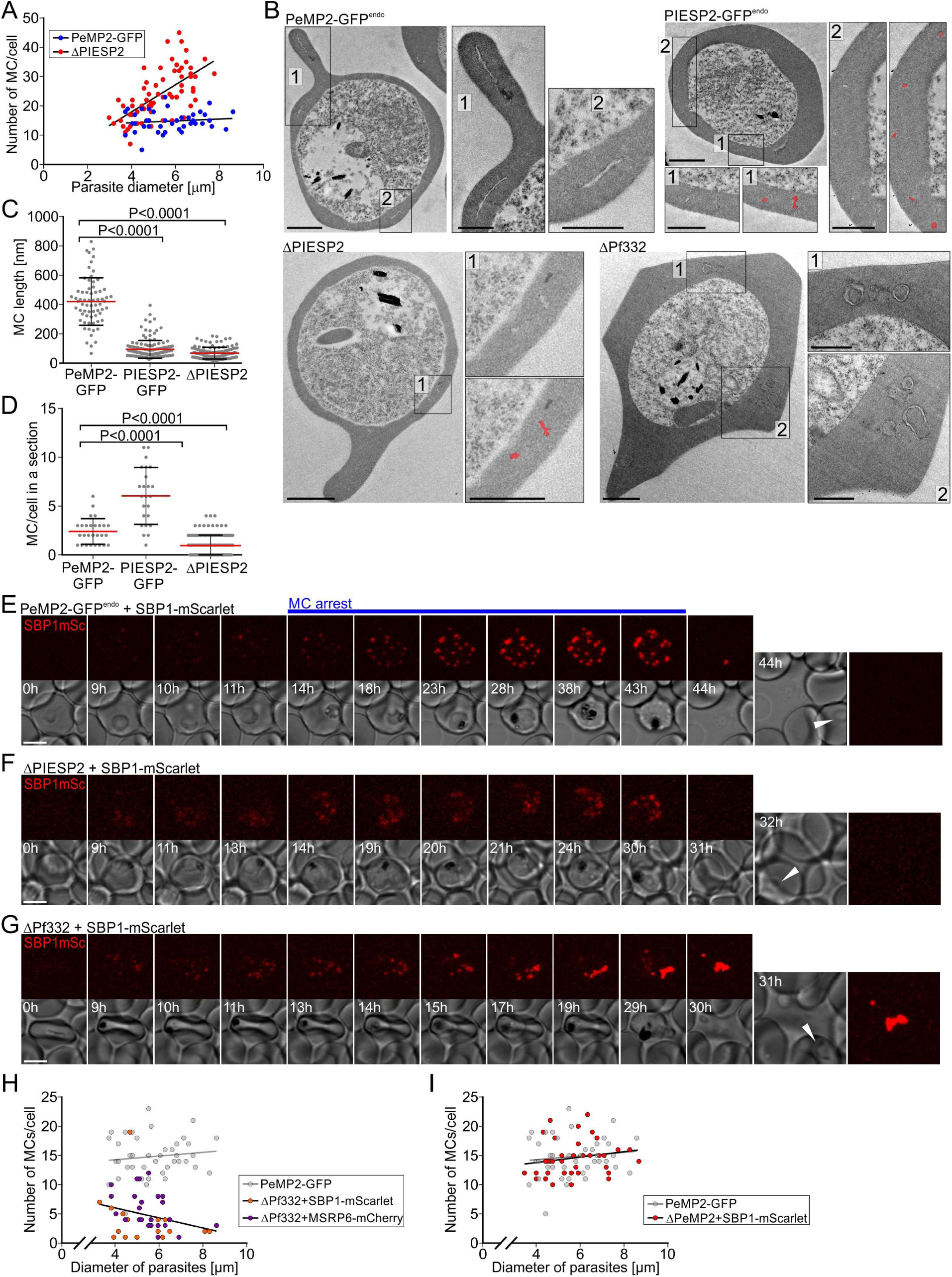
Detailed phenotpyes of PIESP2- and Pf332-TGDs. (A) Quantification of the number of Maurer’s clefts (MCs) per iRBC in the indicated integration cell lines in relation to the parasite age determined by parasite diameter (n = 41 cells for PeMP2-GFP parasites and 67 cells for ΔPIESP2 parasites, data from Fig. 6C). (B) Transmission electron microscopy images of 27 - 36 hours post invasion (hpi) endogenously tagged PeMP2 (PeMP2-GFP) and PIESP2 (PIESP2-GFP) parasites, and ΔPIESP2 or ΔPf332 parasites. Black squares with numbers show enlarged areas. For PIESP2-GFP and ΔPIESP2, MCs are highlighted in red. Scale bars, 0.5 μm. (C, D) Quantification of MC length (C, n = 67 (PeMP2-GFP), 148 (PIESP2-GFP) and 117 (ΔPIESP2) sections), and number of detectable MCs per cell in a section (D, n = 27 (PeMP2-GFP), n = 24 (PIESP2-GFP) and n = 123 (ΔPIESP2) individual sections). Red lines, mean; error bars, SD; two-tailed unpaired t-test, P-values indicated. (E-G) Selected images (full set in Fig. S6) of long-term 3D time-lapse experiments in PeMP2-GFP (E), ΔPIESP2 (F) and ΔPf332 (G) parasites co-expressing episomal SBP1-mScarlet. DIC shows single slice per time point, red signal maximum intensity projection of reconstructed z-stack. Time is indicated as h after start of imaging; re-invasion after completion of the cycle is indicated by white arrows. Onset of MC arrest in PeMP2-GFP indicated by blue line. Scale bars, 5 µm. One representative of 32 (PeMP2-GFP^endo^), 18 (ΔPIESP2), 12 (ΔPf332) cells. (H,I) Quantification of the number of MCs per iRBC of indicated cell lines in relation to the parasite age determined by parasite diameter (n = 17 and 26 cells for ΔPf332+SBP1-mScarlet and ΔPf332+MSRP6-mCherry parasites, respectively (H) and n = cells including those in Fig. 6A; PeMP2-GFP same data as in (A) shown as a reference in both graphs). (A, H and I), linear regression lines indicated.

The apparently stage-specific nature of the phenotype in the PIESP2 mutant prompted us to analyse the fate of the Maurer’s clefts during blood stage development in individual parasites using long term 3D time lapse imaging (Gruring et al., 2011). To visualise the Maurer’s clefts we took advantage of episomally expressed SBP1-mScarlet in the mutant cell line (Fig. 7E-F). The failure of clefts arrest in the PIESP2-TGD parasites was clearly evident by a blurred Maurer’s clefts pattern (due to their movement between individual layers of the z-stack taken for each time point) as soon as the clefts became detectable and this phenotype persisted until the end of the cycle (selected time points in Fig. 7F, full data Fig. S6A). In contrast, the control cell line (PeMP2-GFP^endo^) showed moving clefts when they became first detectable in the cycle (mid ring stage), followed by arrest of their position with transition of parasites to the trophozoite stage (Fig. 7E, Fig. S6B), fitting previous observations of the dynamics of Maurer’s clefts (Gruring et al., 2011). Overall, this indicated that PIESP2 is needed for Maurer’s clefts anchoring and for maintenance of cleft integrity, as its absence led to an increasing fragmentation of the clefts in later stages. The stage-specific nature of the phenotype might explain the discrepancy in previous findings and the different phenotypes in mixed stage cultures.

Time lapse analysis of the parasites with the disrupted Pf332 showed that also in these cells the Maurer’s clefts were mobile from the first time point they became detectable and failed to arrest, although they were more clearly visible as individual foci than in the PIESP2-TGD (Fig. 7G, full time lapse data in Fig. S6C). With increasing age of the parasite, most Maurer’s clefts aggregated into one to few large foci, indicating that also in this mutant the morphology phenotype becomes more pronounced with parasite age and the mixture of cells showing aggregated and non-aggregated Maurer’s clefts is (at least in part) due to different developmental stages (Fig. 7G, H). The EM analysis showed that the Maurer’s clefts were indeed found in aggregates but in addition also showed an altered, more bent and circular morphology (Fig. 7B). We conclude that disruption of Pf332 had a partially similar phenotype to disrupting PIESP2 as in both mutants the clefts failed to arrest but that in the Pf332-TGD parasites the clefts aggregated, whereas in the PIESP2-TGD they fragmented with increasing developmental age of the parasite. As loss of PeMP2 also causes a Maurer’s clefts arrest defect (Fig. 6D), we assessed if this mutant also displays a stage-dependent fragmentation of the Maurer’s clefts that we may have missed in our analysis of asynchronous parasites (Fig. 6A), but this was not the case (Fig. 7I).

### PIESP2-TGD corresponds to a Pf332 and PIESP2 double knock out implicating Pf332 in Maurer’s clefts anchoring

It was previously indicated by antibody data that Pf332 transport depends on PIESP2 (Zhang et al., 2018). We disrupted PIESP2 in the line expressing endogenously GFP-tagged Pf332 using a system permitting the use of SLI for two independent genomic modifications (Cronshagen et al., 2024) (Fig. 8A-B and Fig. S7A). This resulted in an accumulation of Pf332-GFP in the parasite periphery with a faint pool in the host cell but a lack of signal at the Maurer’s clefts (Fig. 8B), demonstrating that Pf332 depends on PIESP2 for transport to the Maurer’s clefts. Hence, the PIESP2-TGD functionally corresponds to a Pf332/PIESP2 double knock out. As in both, the PIESP2- and the Pf332-TGD, Maurer’s clefts anchoring is impaired, this is likely mediated by Pf332 and aggregation of the clefts occurs if PIESP2 is still present.

**Figure 8.**
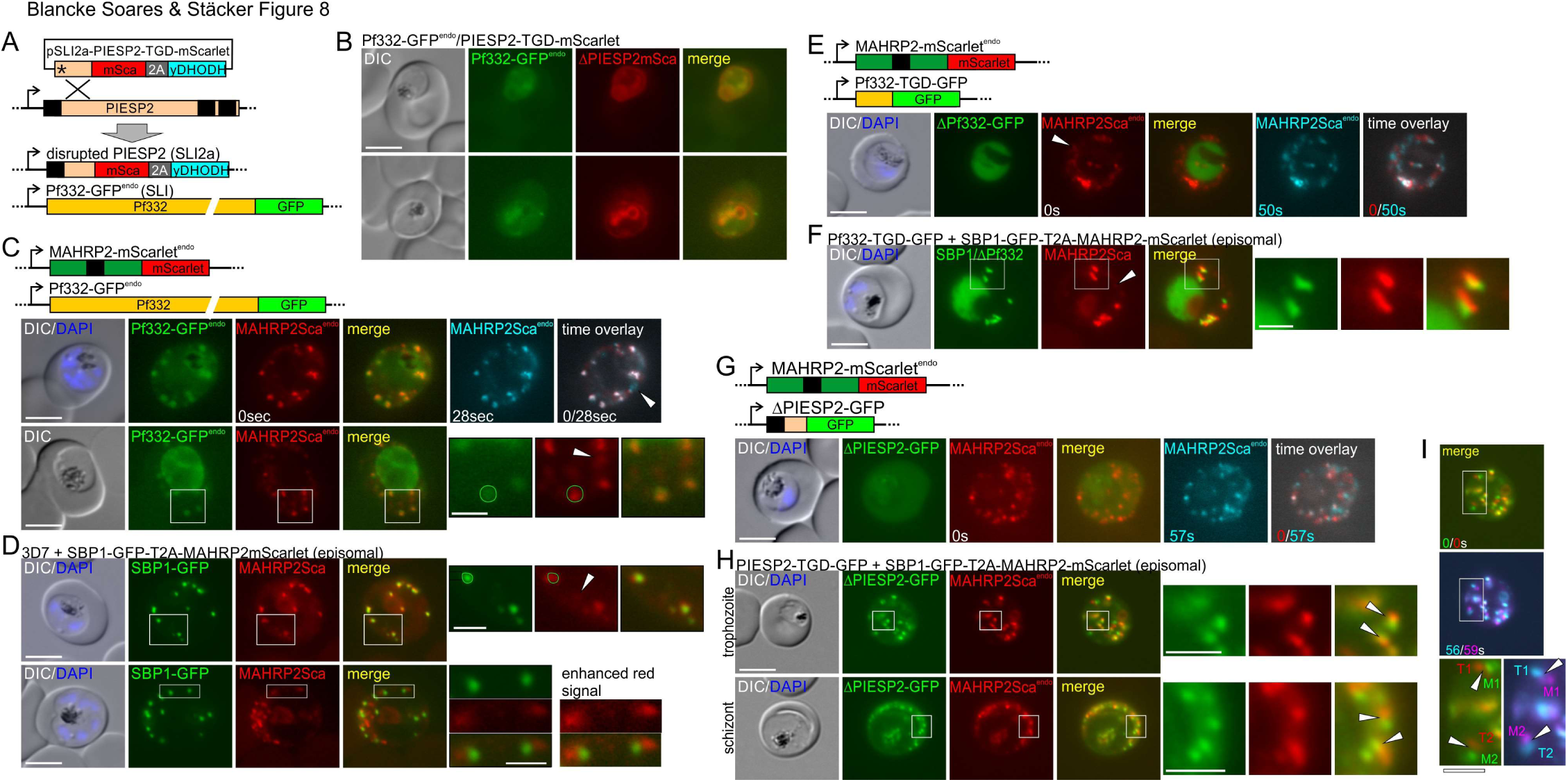
Anchoring defects in PIESP2- and Pf332-TGDs have no profound effect on tether-Maurer’s clefts connection. (A) Schematic of the disruption of PIESP2 by homologous recombination using SLI2a in parasites with SLI-based endogenously GFP-tagged Pf332; mSca (mScarlet); yDHODH, yeast dihydroorotate dehydrogenase; asterisk stop codon; black arrows, native promotors; black boxes, N-terminal signal peptide and transmembrane domains. (B, C) Representative live cell images of Pf332-GFP^endo^ parasites with a disrupted, mScarlet-tagged PIESP2 (ΔPIESP2-mSca) (B) or endogenously mScarlet-tagged MAHRP2 (MAHRP2Sca^endo^) corresponding to tethers. Top panel in (C), short-term time-lapse as done in Fig. 6D. (D) Live cell images of 3D7 parasites episomally expressing SBP1-GFP (MC) and MAHRP2-mScarlet (tethers) using a single expression cassette (SBP1-GFP-T2A-MAHRP2-mScarlet). (E-H) Representative live cell images of parasites with an endogenously mScarlet-tagged MAHRP2 (MAHRP2Sca^endo^) with a disrupted Pf332 (ΔPf332) (E) or PIESP2 (ΔPIESP2) (G) or disrupted Pf332 with episomal SBP1-GFP-T2A-MAHRP2-mScarlet (F). Short term time overlays in (E) and (F) done as in Fig. 6D. (I), Time overlay of merge of red and green signal from the top cell shown in (H) for which the signal in the second time point was converted from green to magenta (SBP1, Maurer’s clefts) and from red to turquoise (MAHRP2, tethers). Individual clefts and tether foci are numbered to show relative position between the two time points. (C-F) Nuclei were stained with DAPI, white arrowheads show fainter mobile tether signals. (H, I), arrowheads show connection of tethers with MC. Schematics of the modified loci are shown for (E) and (G). White boxes show enlarged regions. Scale bars, 5 μm. DIC, differential interference contrast.

MAHRP2 was identified as a marker for the Maurer’s clefts tethers (Pachlatko et al., 2010). As the tethers might be a reason for the arrest of the Maurer’s clefts movement in wt parasites and hence their anchoring (McMillan et al., 2013), we generated endogenously mScarlet-tagged MAHRP2 in the Pf332-GFP^endo^ and Pf332-TGD parasites (Fig. S7B). We first analysed the MAHRP2 signal in the Pf332-GFP^endo^ parasites. Interestingly, the Maurer’s clefts foci of Pf332-GFP^endo^ encompassed the entire MAHRP2 signal (Fig. 8C), differing from other Maurer’s clefts proteins such as e.g. SBP1, which displayed the typical Maurer’s clefts-tether pattern with partially overlapping or neighboring SBP1 and MAHRP2 foci (Fig. 8D). This suggested that Pf332 either localises to both, the clefts and the tethers or that the location of this large protein extends further from the Maurer’s clefts radius than other Maurer’s clefts proteins. The latter is more likely, as the Pf332-GFP signal did not appear to specifically match the MAHRP2 signal but encompassed also regions not positive for MAHRP2 (Fig. 8C). In the parasite with a disrupted Pf332 (Fig. 8E), MAHRP2 was still associated with the aggregated Maurer’s clefts, as evident from expressing SBP1-GFP and MAHRP2-mScarlet together in this cell line (Fig. 8F, note that in this cell line SBP1 and the disrupted Pf332 carry a GFP, but only SBP1-GFP is at the Maurer’s clefts, whereas the truncated Pf332 is in the parasite cytosol and therefore does not interfere with the purpose to visualise the clefts in relation to the MAHRP2-mScarlet marked tethers). In both, parasites with intact or disrupted Pf332, there were faint and smaller MAHRP2 foci not associated with the Maurer’s clefts that were mobile (Fig. 8C-F arrowheads), likely corresponding to the previously described free population of tethers ((Pachlatko et al., 2010); Fig. S7C). There were no profound alterations in the amount of soluble MAHRP2 in the parasite with disrupted Pf332 and those with an intact Pf332 (Fig. S7C). Analysing the relation of SBP1 and MAHRP2 in the PIESP2-TGD confirmed that the Maurer’s clefts and tethers were mobile in this mutant (Fig. 8G). The large number and the movement of these structures in later stage parasites when the PIESP2 phenotype become more profound precluded a finer analysis but similarly to the Pf332 mutant, the tethers still appeared to be attached to the fragmented Maurer’s clefts (Fig. 8H) and remained attached over time (Fig. 8I).

We conclude that ablation of Pf332 does not remove the association of MAHRP2 from the Maurer’s clefts, indicating that failure of clefts anchoring in the Pf332 mutant is not due to failure of tether attachment. However, given the spatial extent of the Pf332-GFP signal, this protein reaches as far from the Maurer’s clefts body as the tethers and this protein might therefore itself connect to other structures. As Pf332 was previously shown to bind host actin *in vitro* (Waller et al., 2010) and host actin was implicated in anchoring of the clefts (Kilian et al., 2013), we next tested if disrupting actin using cytochalasin D (CytD) had any effect similar to our TGD parasite lines. Incubation with CytD starting with synchronous ring stages before Maurer’s clefts arrest occurs did not prevent Maurer’s clefts anchoring and the Maurer’s clefts appeared normal in later stage parasites (Fig. S8A). Short term incubation (10 min) as used previously (Kilian et al., 2013) resulted in small cleft movement in some of the clefts per infected cell in some cells (6 of 17 cells compared to 0 of 18 cells in the control, Fig. S8B). We conclude that CytD treatment had no or only a small effect. Taken together these findings indicate that neither binding to host cell actin nor an altered connection of the tethers to the Maurer’s clefts explain the phenotype in our TGD parasites, suggesting that other or additional factors contribute to anchoring such as e.g. Pf332 binding other host cell cytoskeletal components (Waller et al., 2010).

### Disruption of MSRP6 shows an intermediate Maurer’s clefts phenotype

Next, we analysed if MSRP6 itself influenced the Maurer’s clefts in a similar manner to PIESP2 and Pf332. For this we generated a new line with a disrupted MSRP6 using SLI2 (MSRP6-TGD, Fig. 9A; Fig. S9A), as the previously used knock out misses a larger genomic region encompassing the MSRP6 locus (Kadekoppala et al., 2010). Maurer’s clefts anchoring in the resulting MSRP6-TGD parasites was still intact (Fig. 9B). The number of Maurer’s clefts per cell in the MSRP6 mutant was not significantly different to controls although there was a small trend towards more Maurer’s clefts (Fig. 9C). We noticed that this was predominantly due to late stage parasites that had more Maurer’s clefts than average. Given the fact that the PIESP2 phenotype became more pronounced with parasite age, we tested if this could also be the case in the MSRP6-TGD parasites. Indeed, the number of Maurer’s clefts per cell was increased in dependence of parasite age (Fig. 9D) and when only late stage parasites (diameter ≥6 µm) were analysed (using the late stages shown in Fig. 9C together with additional cells), a significantly increased number of Maurer’s clefts was detected in the MSRP6-TGD parasites (Fig. 9E). This indicated that MSRP6 contributed to the same functions as PIESP2, although at a modest scale.

**Figure 9.**
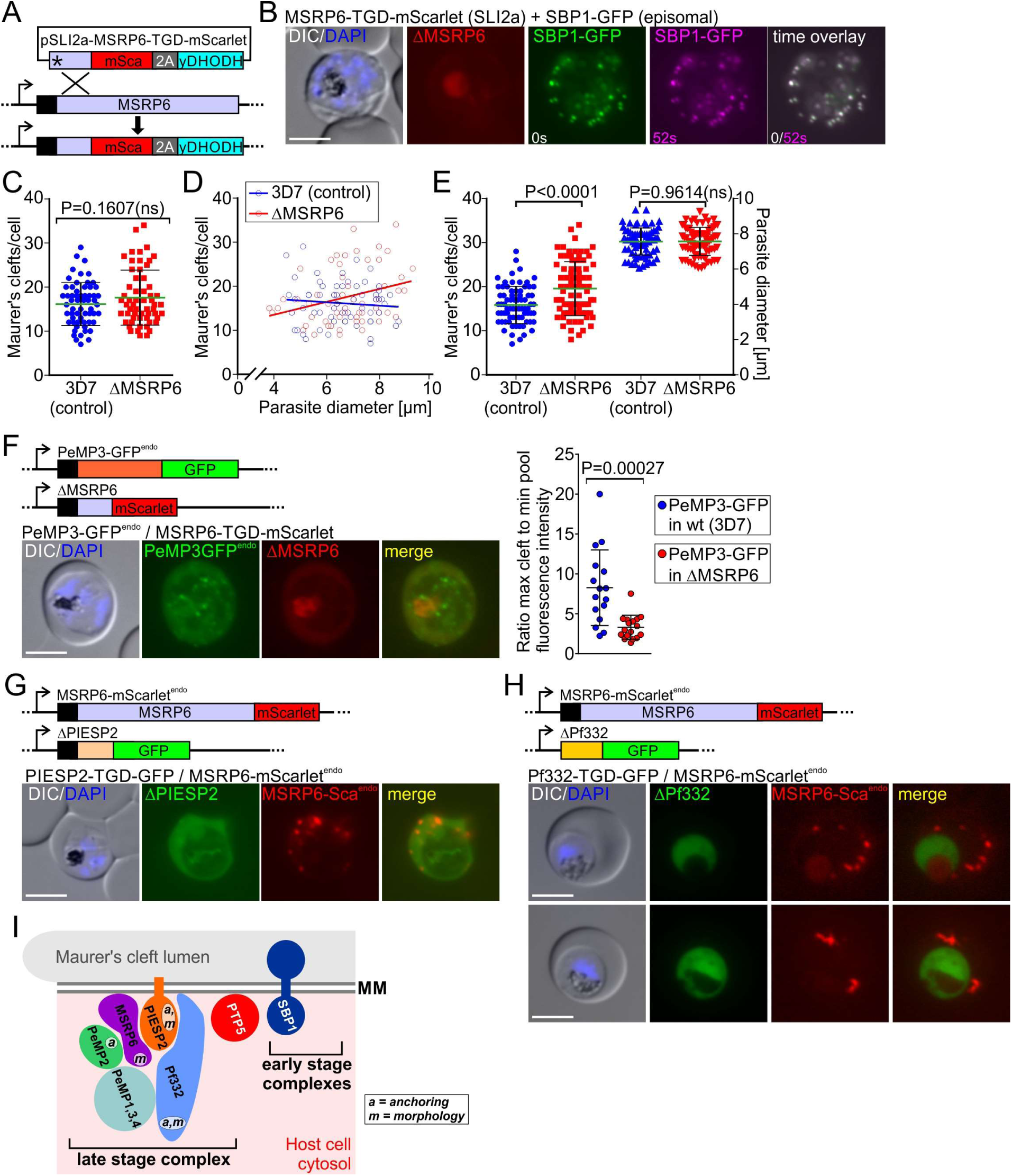
Analysis of the MSRP6-TGD. (A) Schematic MSRP6 disruption using SLI2a; asterisk, stop codon; yDHODH, yeast dihydroorotate dehydrogenase; mSca, mScarlet; black arrows, native promotor; black box, N-terminal signal peptide. (B) Short-term time-lapse live cell images of ΔMSRP6-mScarlet parasites, expressing episomal SBP1-GFP; time overlay as done in Fig. 6D. (C-E) Quantification of the number of Maurer’s clefts per iRBC in the indicated cell lines. (D) Shows the relation to parasite age determined by parasite diameter. Linear regression lines are shown. (C, D), n = 60 and 58 cells for 3D7 and ΔMSRP6 parasites, respectively. (E) Shows only late-stage parasites (diameter ≥6 µm) including the parasites from (C) fulfilling this criterion (n = 74 and 82 cells for 3D7 and ΔMSRP6 parasites, respectively). (F) Live cell images of PeMP3GFP^endo^ parasites with a disrupted MSRP6 (ΔMSRP6-Sca). The graph shows a quantification of the Maurer’s clefts-associated PeMP3-GFP signal relative to the distributed signal (soluble population of the protein) in the indicated cell lines (n = 16 and 17 cells for PeMP3-GFP in 3D7 and PeMP3-GFP in ΔMSRP6 parasites, respectively). (G, H) Live cell images of ΔPIESP2-GFP (G) or ΔPf332-GFP (H) parasites with an endogenously mScarlet-tagged MSRP6 (MSRP6-Sca^endo^). (I) One possible model of MSRP6 complex (’late stage complex’) at the Maurer’s clefts. (B, F-H), nuclei were stained with DAPI; scale bars, 5 μm; DIC, differential interference contrast. (F-H) Show schematics of the modified loci. (C, E, and F), mean and error bars (SD) are shown; two-tailed unpaired t-test, P-values indicated.

Taking advantage of the MSRP6-TGD parasites we sought to confirm the suspected dependence of PeMP3 on MSRP6 for Maurer’s clefts recruitment (Fig. 5F). We endogenously tagged PeMP3 with GFP in the MSRP6-TGD (Fig. S9A) and compared its distribution to that of the cell line expressing PeMP3-GFP^endo^ with an intact MSRP6. Loss of MSRP6 reduced PeMP3 Maurer’s clefts association as judged by a reduced Maurer’s clefts intensity but increased cytoplasmic pool in the host cell in MSRP6-TGD trophozoites (2 nuclei and older when PeMP3 associates with the Maurer’s clefts) compared to PeMP3-GFP^endo^ parasites with an intact MSRP6 locus (Fig. 9F). Together with the finding that overexpressed MSRP6 led to increased Maurer’s clefts association of PeMP3 (Fig. 5F), it can be assumed that a direct or indirect interaction of this protein with MSRP6, at least in part, is responsible for its recruitment to the Maurer’s clefts and supports the BioID and IP data.

Loss of Pf332, PIESP2, PeMP1, 3 and 4 did not abolish MSRP6 Maurer’s clefts association (Fig. 9G, H and Fig. S9B, C), demonstrating that they are either not the direct or not the only MSRP6 contact at the Maurer’s clefts. Overall, these findings suggest multiple interactions between the MSRP6 complex proteins, hinting at a complex network of proteins including Pf332, PIESP2, PeMP1-4 and MSRP6 (Fig. 9I). This is consistent with the finding that central functions of Pf332, PIESP2 and PeMP2 depend only to a small extent (Maurer’s clefts maintenance) or not (Maurer’s clefts anchoring) on MSRP6.

### Effect of Maurer’s clefts anchoring on virulence protein transport and cytoadherence

The assembly of the MSRP6 complex at the Maurer’s clefts takes place at the transition to the trophozoite stage when PfEMP1 reaches the host cell surface (Kriek et al., 2003) and when the anchoring of the Maurer’s clefts occurs (Gruring et al., 2011). Clefts anchoring might therefore be a prerequisite for PfEMP1 trafficking and our mutants provide a means to test this. Previous work showed conflicting results regarding the role of PIESP2 and Pf332 in PfEMP1 transport (Glenister et al., 2009; Hodder et al., 2009; Maier et al., 2008; Moll et al., 2007). To assess effects on PfEMP1 transport and cytoadhesion we used a recently established SLI-based system to HA-tag and activate a specific PfEMP1 (in this case IT4var01), resulting in IT4 parasites all expressing IT4var01 from the endogenous locus (Cronshagen et al., 2024). In these IT4var01-HA^act^ parasites we then used SLI2 (Cronshagen et al., 2024) to disrupt either PIESP2, Pf332 or MSRP6 (Fig. 10A-C, Fig S10A). As a positive control IT4var01-HA^act^ line we also disrupted SBP1 (Fig. 10D, Fig S10A), a protein known to be needed for cytoadherence (Cooke et al., 2006; Maier et al., 2007). We used IT4 parasites rather than 3D7 for these experiments because 3D7 parasites (at least those used in our lab) lost the ability for efficient cytoadherence (Cronshagen et al., 2024).

**Figure 10.**
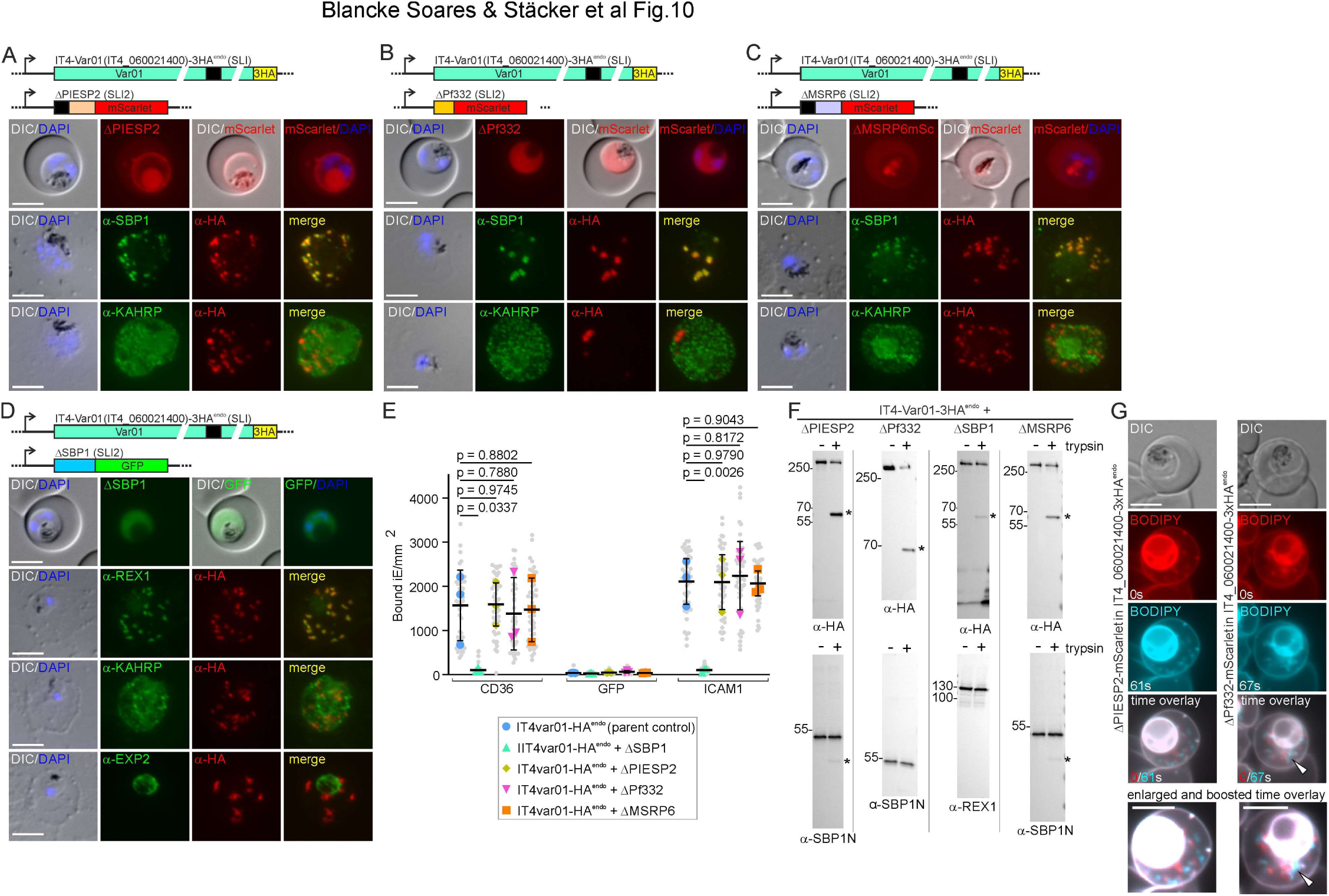
Cytoadherence and PfEMP1 transport in MSRP6 complex member mutants. (A-D) IT4 parasites with a SLI-activated Var01 (IT4var01-HA^act^ parasites) with disruptions of the genes encoding the indicated proteins (schematics above the panels). Images show live cells demonstrating loss of localisation of the disrupted protein (top), or IFAs with acetone fixed cells showing the PfEMP1 (a-HA) together with SBP1 (Maurer’s clefts) or KAHRP. For the SBP1 disruption REX1 was used instead of SBP1 and EXP2 to show the PVM. Nuclei were stained with DAPI; scale bars, 5 μm; DIC, differential interference contrast. (E) Binding of iRBCs of the indicated cell lines to CHO cells expressing CD36, ICAM1 or GFP shown by super plots (Lord et al., 2020) (three independent experiments with technical replicas. Error bars (SD) indicates mean of averages of biological replicates with SD; P-values (unpaired t-test) are shown; small grey dots: bound iE infected erythrocytes) per field of view extrapolated to mm^2^. Coloured shapes: average of bound iE/mm2/replicate). F) Immunoblots with extracts of saponin-lysed parasites of the indicated the cell lines treated with or without trypsin (n = 2, replica shown in Fig. S11B). α-HA detects the activated Var01 and the protected fragment is indicative of surface exposure. α-SBP1N served as control that the RBC was not breached which would also result in a protected fragment. Asterisks indicate protected fragments. (G) Fluorescence microscopy images of Bodipy stained parasites of the indicated cell lines, showing the same cell after a short interval. Second time point image converted to turquoise and overlaid (time overlay) with the first time point to illustrate Maurer’s clefts movement. A boosted and enlarged image of the overlay is shown at the bottom. Arrow shows a ΔPf332-typical aggregate. Scale bar, 5 μm. DIC, differential interference contrast.

Initial binding experiments indicated no major differences between the PIESP2, Pf332 and MSRP6-TGD parasites compared to the control (Fig. S10B). To more accurately assess this, we used an improved binding assay that includes semi-automated scoring of binding which increases the throughput and therefore the accuracy of the binding assay (Cronshagen et al., 2024). Using this assay, the IT4var01-HA^act^ parasites showed the previously reported binding to ICAM-1- and CD36- but not GFP-expressing CHO cells (Fig. 10F). After disruption of PIESP2 (corresponding to a double knock out of PIESP2 and Pf332), the IT4var01-HA^act^ parasites still exported PfEMP1 to the host cell surface (determined by surface trypsin assays) and bound to these receptors at levels comparable to the parent IT4var01-HA^act^ parasites (Fig. 10E, F). Equally, disrupting Pf332 or MSRP6 in the IT4var01-HA^act^ parasites did not significantly affect the binding of RBCs infected with these parasites compared to the IT4var01-HA^act^ parent (Fig. 10E). In contrast, the control line with the disrupted SBP1 lost receptor binding (Fig. 10E). Trypsin assays indicated that these parasites had only very small amounts of surface exposed Var01, contrasting with the IT4var01-HA^act^ parent and the parasites with disruptions of either PIESP2, MSRP6 or Pf332, fitting with the binding phenotype of these parasite lines (Fig. 10F and Fig. S10B). As the tagged PfEMP1 in the SBP1 mutant still showed the typical Maurer’s clefts pattern by IFA, this indicated that in the SBP1 mutant only surface translocation but not export of PfEMP1 to the host cell was negatively affected, confirming previous data (Cooke et al., 2006). We conclude that disruption of PIESP2, Pf332 and MSRP6 does not noticeably impair cytoadherence and does not appear to negatively affect PfEMP1 transport and function.

Finally, we wished to ensure that the Maurer’s clefts phenotypes observed in our mutants in 3D7 were also present in the mutants of the IT4var01-HA^act^ parasites. While the IFAs indicated that the Maurer’s clefts phenotype of the PIESP2 and Pf332 disruptions were similar in IT4 parasites to those observed in 3D7 parasites (Fig. 10A, B), this analysis did not permit to assess Maurer’s clefts movement. To confirm that the Maurer’s clefts arrest phenotype also occurred in the IT4var01-HA^act^ parasites, we stained the parasites with BODIPY-ceramide to visualise membranous structures in the iRBC. These experiments showed that the IT4var01-HA^act^ parasites with disrupted PIESP2 or disrupted Pf332 both had mobile Maurer’s clefts in trophozoites (Fig. 10 G).

Altogether these data indicate that MSRP6-complex dependent Maurer’s clefts anchoring is not critical for PfEMP1 surface transport and cytoadherence. Moreover, the disintegration of the Maurer’s clefts seen in the PIESP2 mutant and the aggregation in the Pf332 mutant also do not influence cytoadherence.

## Discussion

The Maurer’s clefts are one of the most prominent host cell modifications that were already detected in Giemsa-stained blood smears of malaria patients over 100 years ago (Maurer, 1902). A central function of these structures in cytoadherence became apparent through independent lines of evidence: (i) several Maurer’s clefts proteins are needed for PfEMP1 trafficking (Carmo et al., 2022; Cooke et al., 2006; Maier et al., 2007; Maier et al., 2008; McHugh et al., 2015; McHugh et al., 2020; Rug et al., 2014; Spycher et al., 2008), (ii) mutants or conditions negatively affecting Maurer’ clefts architecture also caused a loss of cytoadherence (Dixon et al., 2011; McHugh et al., 2015; McHugh et al., 2020; Rug et al., 2014), and (iii) PfEMP1 itself or other proteins important for cytoadherence such as KAHRP are found at least transiently at the Maurer’s clefts before reaching the host cell surface (Wickham et al., 2001). A striking event in the biogenesis of the Maurer’s clefts is their transition from an apparently free moving state in the host cell to becoming fixed in position in a short time window during parasite development (Gruring et al., 2011). This time window coincides with the stage PfEMP1 reaches the host cell surface and parasites become cytoadherent, giving rise to the idea that these two phenomena are connected (Kriek et al., 2003; McMillan et al., 2013). However, the parasite proteins involved in Maurer’s clefts anchoring have remained elusive, precluding establishment of a direct causal link between this anchoring and cytoadherence. Starting from an unusual trafficking domain we here identified a complex involved in Maurer’s clefts anchoring. Three proteins of this complex prevented cleft arrest when disrupted, PIESP2, PeMP2 and Pf332 but all showed partially different additional phenotypes. Furthermore, two of these, PIESP2 and PeMP2 as well as a further protein, MSRP6 itself, were needed to maintain the integrity of the Maurer’s clefts and their loss caused different levels of disintegration of the Maurer’s clefts. The similarity of the phenotypes between the proteins indicates that the MSRP6-defined complex is a functional unit but also suggests a certain level of redundancy between the complex members.

There is evidence for various interactions of different Maurer’s clefts proteins based on immunoprecipitation approaches, for instance in recent work studying PTP1, SBP1, PIESP2 or GEXP07 (Davies et al., 2023; McHugh et al., 2020; Rug et al., 2014; Takano et al., 2019; Zhang et al., 2018). In our approach we used BioID to identify interactors of the MSP7-like domain of MSRP6 which permits identification of interactors and compartment neighbours in the living cell (Kimmel et al., 2021; Roux et al., 2012). While this approach does not necessarily identify true interactors, it enriches for proteins that are consistently in a close spatial arrangement with the bait in the functioning cell. We confirmed several of the hits using co-immunoprecipitation with SBP1 as a stringent negative control. Overall, these results provide evidence for an MSRP6-specific complex including PeMP1-4, PIESP2 and Pf332 that is not or only peripherally or transiently in contact with SBP1 which showed no or only small enrichment in our immunoprecipitations. Functionally this is consistent with MSRP6 complex members not having an essential role in PfEMP1 transport which contrast with SBP1 or several other proteins found to interact with SBP1.

Our data indicates multiple interactions between the MSRP6 complex members and removal of individual components did not disband the entire complex. A first indicator for intra complex relationships became apparent for MSRP6 and PeMP3: PeMP3 depended on MSRP6 for Maurer’s clefts attachment and hence binds the complex at least in part via MSRP6 (Fig. 9I). It is of particular note that many of the MSRP6 complex proteins do not contain transmembrane domains, indicating that their location in the host cell is defined through protein-protein interactions, as otherwise these proteins are expected to be soluble in the RBC cytosol. PIESP2 is an exception and might be one protein anchoring the complex to the Maurer’s clefts (Fig. 9I). However, MSRP6 was still found at the clefts when PIESP2 and Pf332 were absent, indicating that other connectors of the MSRP6 complex to the Maurer’s clefts membrane exist (potentially also via protein-lipid interactions). Previous work using electron microscopy showed that MSRP6 is located in cloudy electron dense regions at the Maurer’s clefts (Heiber et al., 2013). It is possible that the proteins of the MSRP6 complex form an interconnected meshwork surrounding the Maurer’s clefts.

A main function in anchoring the Maurer’s clefts in the host cell can be ascribed to Pf332. The role of PIESP2 in anchoring is at least in part indirect, as its disruption also led to a failure of Pf332 to reach the Maurer’s clefts. Interestingly, previous work indicated that Pf332 binds the host cell cytoskeleton and *in vitro* work indicated direct binding to actin (Waller et al., 2010). Actin has previously been implicated in Maurer’s clefts anchoring (Cyrklaff et al., 2011; Kilian et al., 2013). This agrees with our work implicating Pf332 in this function. However, we did not find profound changes in the anchoring of the Maurer’s clefts when parasites were grown in the presence of CytD. The tethers were still associated with the Maurer’s clefts in our mutants. While it cannot be excluded that the host cell binding end of the tethers was negatively affected and that this contributed to the reduced anchoring of the Maurer’s clefts, the proteins studied here are located at the Maurer’s clefts which likely would not affect the Maurer’s clefts distal end of the tethers. Interestingly, Pf332-GFP reached further from the Maurer’s clefts circumference than other proteins, as evident from a fluorescence signal fully encompassing the tethers as well as the clefts in our live cell imaging. Pf332 might therefore directly reach structures of the host cell (such as the RBC cytoskeleton) and connect them with the Maurer’s clefts independently of the tethers. Taken together, our work indicates that Maurer’s clefts anchoring is not mediated by binding to host cell actin alone and that the phenotype is also not explained by a failure of the tethers to attach to the Maurer’s clefts. This indicates that the contribution of actin and the tethers in anchoring of the Maurer’s clefts needs to be more precisely defined.

Taking advantage of a cytoadherent parasite line expressing a specific HA-tagged PfEMP1 (Cronshagen et al., 2024) we tested the importance of Maurer’s clefts anchoring for cytoadherence. Parasites that lacked PIESP2 (and hence also Pf332) or Pf332 alone still successfully transported PfEMP1 to the host cell surface and were still cytoadherent. While it could be argued that the proteins of the MSRP6 complex might function only after PfEMP1 reaches the host cell surface, the fact that the Maurer’s clefts never arrested in our mutants speak against this idea, overall excluding a requirement of this phenomenon for PfEMP1 surface transport and function at the host cell membrane. It is however possible that a connection of the Maurer’s clefts to the host cell surface via actin (possibly transient or dynamic) still exists in these mutants and that this might still permit PfEMP1 transport even though the clefts fail to arrest.

Why the Maurer’s clefts aggregate in the Pf332 mutant is unclear, but as they do not in the PIESP2 disruption (which also lacks Pf332 at the Maurer’s clefts), it can be attributed to PIESP2 being at the Maurer’s clefts without Pf332. As these two proteins are part of the same complex, it is possible that interaction interfaces of PIESP2 are not saturated when Pf332 is absent and that this leads to an aggregation, for instance if interactions with components of the MSRP6 complex on other Maurer’s clefts occur. If Pf332 is present and anchoring of the clefts takes place, the Maurer’s clefts do not get into contact at the time the MSRP6 complex members reach the Maurer’s clefts, preventing an aggregation. A propensity for self-interaction in the absence of anchoring would provide one explanation for the need of the Maurer’s clefts to be fixed in place. Alternatively, as Pf332 seems to extend far from the Maurer’s clefts circumference, this protein might be needed as a sterical buffer to prevent undesired interactions between proteins of different Maurer’s clefts which would connect them and lead to the phenotype of aggregated clefts.

The fragmentation of the Maurer’s clefts in the PIESP2 mutant had no consequences for cytoadherence and PfEMP1 transport. Loss of PTP1 was also reported to cause a similarly altered architecture of the Maurer’s clefts but in contrast resulted in a PfEMP1 transport defect (Rug et al., 2014). However, PTP1 is expressed early in the cycle, and its loss might affect the Maurer’s clefts before the MSRP6 complex forms at the Maurer’s clefts, although the stage-specificity of the PTP1 phenotype is not known. It is possible that the Maurer’s clefts morphology phenotype in our mutants manifests later in the cycle than that of PTP1 and therefore might not affect PfEMP1 surface display. The aggregation of the Maurer’s clefts in our TGDs also did not affect cytoadherence. In contrast, the aggregation or other morphological alterations seen in REX1, PTP7 and GEXP7 mutants do affect PfEMP1 trafficking (Carmo et al., 2022; Hanssen et al., 2008a; McHugh et al., 2020) and may again do so because these proteins are expressed early in the cycle. Alternatively, the morphological changes with the Maurer’s clefts seen in these mutants could be secondary effects of the phenotype causing the PfEMP1 transport defect rather than directly causing it.

Our findings raise the question why the Maurer‘s clefts are anchored if not for cytoadherence and why the integrity of the Maurer’s clefts needs to be maintained in the second half of the cycle in red blood cells. As cleft anchoring is not essential for parasite growth *in vitro*, it is possible that it serves functions *in vivo*. Such functions are difficult to test and at present remain speculative. Anchoring of the clefts for instance could permit efficient egress (clefts attached to the hull of the host cell might interfere less with the release of merozoites than if free in the host cell). Maintaining clefts integrity and anchoring might also keep these antigen-rich structures from being released in multiple small and free fragments into the serum when the host cell ruptures which might lead to excessive immune reactions harming the host and thereby diminishing parasite transmission.

Interestingly, the domain in MSRP6 that is necessary and sufficient for its Maurer’s clefts attachment corresponds to the MSP-7 like domain that defines the MSP7-related proteins (Kadekoppala et al., 2010; Mello et al., 2002). As MSRPs are found in different locations of the cell (Heiber et al., 2013; Kadekoppala et al., 2010), it is possible that the MSP7-like region is a general interaction domain that mediates protein-protein interactions in proteins of different complexes and locations.

This work also has interesting implications for protein export. Firstly, it indicates that also SP-PNEPs contain an N-terminal export domain. Hence, it seems to be a unifying rule that the mature N-terminus of exported proteins (unprocessed if the protein lacks a signal peptide such as in ‘classical’ PNEPs or after cleavage of the signal peptide or in PEXEL proteins the PEXEL (Boddey et al., 2013; Boddey et al., 2009; Gruring et al., 2012; Tarr et al., 2013)), is sufficient to mediate export. Although MSRP6 also conforms to this unifying rule, its C-terminal MSP7-like homology region (corresponding to the MAD) is also sufficient for export. Interestingly, in *P. yoelii* both N-terminal and central parts of PNEPs that contain hydrophobic domains were shown to contain export signals composed of semi-conserved amino acids and secondary structure requirements (Siau et al., 2014). It is possible that this region in MSRP6 mediates interaction to other exported proteins already during transport within the parasite, and hence might be exported by piggy back with a protein containing an N-terminal export domain. Piggy back export would also explain why Pf332 is not exported when PIESP2 is missing. However, how such arrangements can be maintained through PTEX which requires unfolding of the transported proteins (Gehde et al., 2009; Gruring et al., 2012), is unclear.

This work identified a complex at the Maurer’s clefts that assembles in the second half of the asexual development of the parasite in the host cell and comprises proteins needed for Maurer’s clefts anchoring and for the maintenance of their structural integrity. Unexpectedly, we find that Maurer’s clefts anchoring and maintenance was not needed for PfEMP1 transport and cytoadhesion, indicating that the Maurer’s clefts need to be anchored and kept intact for hitherto unknown purposes.

## Supporting information

Supplementary Figures with legends

Data S1 (plasmids)

## Acknowledgments

This work was funded by the German Research Foundation (DFG) grant SP1209/1-3; JS thanks the Jürgen Manchot Stiftung for funding; ABS was associated with the GRK1459 of the DFG and was supported by the Leibniz Center Infection (LCI) Graduate School “Model Systems of Infectious Diseases”; WAMH was financially supported by the Netherlands Organisation for Scientific Research (NWO Veni fellowship 722.012.004); MK thanks the priority program SPP1580 (DFG grant SP1209/2-1). We are grateful to Brian Cooke for the rabbit α-KAHRP antibodies, to Jacobus Pharmaceuticals for WR99210 and the European Malaria Reagent Repository (http://www.malariaresearch.eu) for α-EXP2 antibodies.

## Author contributions

ABS, devising experiments, carrying out experiments, analysing data, writing paper, preparing figures

JS, devising experiments, analysing data, carrying out experiments, writing paper, preparing figures

SS, carrying out experiments, analysing data

WAMH, carrying out experiments, analysing data, writing paper, preparing figures

NGM, carrying out experiments, analysing data

JC, carrying out experiments, analysing data

HS, carrying out experiments

SF, carrying out experiments

KH, carrying out experiments, analysing data

UF, carrying out experiments

PMR, devising experiments, carrying out experiments

BB, carrying out experiments

MKN, carrying out experiments

IB, devising experiments, analysing data, supervising research

RB, devising experiments, analysing data, supervising research

TS, conceiving study, supervising research, devising experiments, analysing data, carrying out experiments, writing paper, preparing figures

All authors commented on the manuscript and approved submission.

## Data availability

The mass spectrometry data is available from PRIDE under the accession number PXD008208. All other data are in the paper or available from the corresponding author upon request.

---

**Table S1: Quantitative Bio-ID of MSRP6 MAD interactome, Maurer’s cleft and host cell cytosolic proteome** (A-D) Quantitative Bio-ID results of all proteins significantly enriched with FDR<5% in at least 2 out of the 4 replicates of REX3^trunc^-MAD- (REX3-MAD) over STEVOR^1-260^- (STEVOR) -GFP-BirA* (A), REX3^trunc^-MAD-over-REX3^trunc^-(REX3) (B) or STEVOR-over-REX3 (C) or of all proteins identified in the Bio-ID experiment no matter whether they are significant on not (D). Normalized log2 ratios per comparison, two-sided significance B-values, significant enrichment with FDR<5%, number of identified peptides and unique peptides, % sequence coverage, protein MW and sequence length, log10 intensities and iBAQ values are listed. Yellow highlighted proteins selected for follow-up as putative MAD interactors, while pink and orange highlight proteins with moderate to no enrichment as MAD interactor but enrichment in the Maurer’s cleft proteome. Grey highlights indicate proteins belonging to variantly expressed multigene families (STEVOR, RIFIN, PfEMP1) that are likely false-positives. See also Fig. 4 and Fig. S1.

**Table S2:** Summary of Co-IP quantification.

|  | <u>enrichment MSRP6 over SBP1 in eluate</u> |  |  |  |  |  |
| --- | --- | --- | --- | --- | --- | --- |
| replica | PIESP2-GFP IP | PF332-GFP IP | PeMP1-GFP IP | PeMP2-GFP IP | PeMP3-GFP IP | PeMP4-GFP |
| n = 1 | 686.28 | 30.43 | 2886.35 | 1724.84 | 67.03 | 23.64 |
| n = 2 | 6.62 | 19.5 | 623.70 | 116.77 | 110.74 | 24.74 |
| n = 3 | 320.3 | 214.24 | 3312.99 | 2236.96 | 3010.110 | 134.92 |
| n = 4 |  |  | 2853.21 |  |  |  |
| average | **337.73** | **88.06** | **2419.06** | **1359.52** | **1062.63** | **61.10** |

## Materials and Methods

### Plasmid constructs

For episomal expression of GFP- and BirA*-tagged proteins DNA sequences were PCR amplified from *P. falciparum* 3D7 cDNA and cloned into pARL1-GFP (Crabb et al. 2004) using either conventional cloning or one-step isothermal DNA assembly (Gibson et al., 2009). mCherry co-expression plasmids were generated by ligation of PCR amplicons into pARL2-mCherry. For simultaneous episomal expression of SBP1 and MAHRP2, DNA was PCR amplified from *P. falciparum* 3D7 gDNA or plasmid DNA and inserted into pARL2 under control of the *mal7*-promotor (Gruring et al., 2011) using one-step isothermal DNA assembly.

For endogenous FKBP-GFP tagging and targeted gene disruptions, DNA was PCR amplified from *P. falciparum* 3D7 gDNA and inserted into pSLI-FKBP-GFP and pSLI-TGD, respectively, (Birnbaum 2017) using one-step isothermal DNA assembly. For endogenous PfEMP1-tagging, DNA was PCR amplified from *P. falciparum* IT4 gDNA and inserted into pSLI-3xHA (Mesén-Ramírez et al. 2019) using conventional cloning. For double-integrations (tagging or TGDs), DNA was PCR amplified from *P. falciparum* 3D7 gDNA or plasmid DNA and inserted into pSLI2a-mScarlet or pSLI2a-GFP using either conventional cloning or one-step isothermal DNA assembly. The SLI2a-vector was generated by replacing the selection markers of pSLI (manuscript in preparation). For episomal selection, the blasticidin S deaminase (BSD) gene (Mamoun et al., 1999) was utilised. Integration of the SLI2a-plasmid into the parasite genome was selected using the yeast dihydroorotate dehydrogenase (yDHODH) gene (Ganesan et al., 2011). All plasmids were verified by sequencing. Sequences of plasmids and inserts are provided in supplemental file S1.

### Parasite culture and transfection

*P. falciparum* (3D7 or IT4 (FCR3S1.2)) blood stage parasites were cultured in 0+ erythrocytes (UKE, Hamburg, Germany) at 37°C in a controlled atmosphere (5% CO_2_, 1% O_2_, 94% N_2_) in RPMI 1640 supplemented with 0.5% AlbuMAX under standard conditions (Trager and Jensen, 1976). Parasites were transfected with 50-100 µg DNA using either the Amaxa system (Lonza) or the Biorad system as previously described (Moon et al., 2013; Wu et al., 1995). Transgenic parasites were selected with 4 nM WR99210 (Jacobus Pharmaceuticals) or 2 μg/ml blasticidin S (Invitrogen). To select for integrants by SLI, parasites were treated with G418 to a final concentration of 400 µg/ml as previously described or with 0.9 μM DSM1 (BEI resources) (Birnbaum et al., 2017). Correct integration was confirmed by PCR on isolated gDNA from the respective transgenic strains (primers in Table S3).

### Immunofluorescence assays (IFA)

Parasites washed in PBS were transferred to 10-well glass slides, air-dried and fixed in acetone for 30 minutes. The cells were re-hydrated with PBS for 15 minutes and non-specific binding sites blocked with 3% BSA in PBS for 30-60 minutes. Antibodies were applied in 3% BSA in PBS and staining performed according to standard procedures (Spielmann et al., 2003). Primary antibodies used were: mouse monoclonal (mixture of clones 7.1 and 13.1) anti-GFP (Roche), 1:500; rabbit polyclonal anti-GFP (Thermo5) anti-GFP, 1:400; mouse polyclonal anti-SBP1-C (Mesen-Ramirez et al., 2016), 1:1000; mouse polyclonal anti-MSRP6 (Heiber et al., 2013), 1:250; rat anti-HA (Roche), 1:500; mouse anti-EXP2 (European Malaria Reagent Repository, http://www.malariaresearch.eu), 1:2000; rabbit anti-KAHRP (kind gift of Brian Cooke), 1:500; rabbit anti-REX1 (Mesen-Ramirez et al., 2016), 1:20000. Secondary antibodies used were: goat anti-mouse-Alexa488, goat anti-mouse-Alexa594, donkey anti-rabbit-Alexa488, donkey anti-rabbit-Alexa594, goat anti-rat-Alexa594 (Invitrogen), all used at 1:2000. The secondary antibody solution contained 1 µg/ml DAPI. Slides were mounted using DAKO fluorescent mounting medium (Dako).

### Imaging

Microscopy of parasites was performed as described previously (Gruring and Spielmann, 2012). Before imaging, parasites were incubated for 10 min with 1 μg/ml DAPI to stain parasite nuclei. Wide field epifluorescence imaging was done with a Zeiss AxioImager M1 with a Hamamatsu Orca C4742-95 camera using either a 100×/1.4–numerical aperture lens or a 63×/1.4–numerical aperture lens. Images were processed and overlays done in Corel Photo-PaintX6.

3D time lapse imaging was done with a FluoView 1000 confocal microscope (Olympus) as described (Gruring et al., 2011). Three-dimensional reconstructions were generated with Imaris 7.7.2, and maximum-intensity projections were processed in Corel Photo Paint.

### Trypsin cleavage assay to detect surface exposed PfEMP1

Trypsin assays were done as described (Waterkeyn et al., 2000). Briefly, trophozoite-stage parasites were purified from total cultures using a Percoll gradient using a Percoll gradient as described (Heiber and Spielmann, 2014). Thereafter the trophozoite-infected RBCs were washed twice with 1xPBS, divided into two reaction tubes and parasites were incubated either with 125 µg/ml TPCK-treated trypsin (Merck) or with 1xPBS at 37°C for 45 minutes. After incubation, soybean trypsin inhibitor (Sigma Aldrich) was added to a final concentration of 1 mg/ml to the parasites suspension and kept on ice for 10 minutes to stop the reaction. After one washing step with 1xPBS, the parasites were re-suspended in 1xPBS containing 1 mg/ml soybean trypsin inhibitor and 25x protease inhibitor cocktail (Roche) before the parasites were pelleted and lysed in lysis buffer (4% SDS, 0.5% Triton X-100 and 0.5x PBS in dH2O). Lysates were frozen at −20°C until further analysis by SDS-polyacrylamide gel electrophoresis (SDS-PAGE) and western blotting.

### Immunoblots and biotin-detection blots

Western blots were done in principle as described previously (Spielmann et al., 2003). For the detection of biotinylated proteins in BirA* expressing parasites, the corresponding *P. falciparum* cultures were incubated with 20 mM Biotin (Sigma) for 24-48 hours before harvesting. Parasite lysates were generated by purifying trophozoites and schizonts from 500 µl packed iRBCs with a parasitemia of 5-10% using a Percoll gradient (Heiber and Spielmann, 2014). The isolated infected RBCs were washed in PBS twice and lysed in 4% SDS, 0.5% Triton X-100 in 0.5× PBS with protease inhibitors (Roche) and frozen at −20 °C. After thawing, extracts were centrifuged at 16,000g, and Laemmli sample buffer was added to the supernatants. The protein extracts were separated by SDS-PAGE and were subsequently transferred to Amersham Protran membranes (GE Healthcare) in a tankblot device (BioRad) using transfer buffer (0.192 M Glycine, 0.1% SDS, 25 mM Tris) with 20% methanol. Membranes were incubated with antibodies in 5% skim milk (Roth) in 1× PBS. Dilutions for primary antibodies were: mouse monoclonal (mixture of clones 7.1 and 13.1) anti-GFP (Roche), 1:1000; rabbit polyclonal anti-GFP (Thermo), 1:1000; rat anti-HA (Roche) 1:1000; mouse anti-MSRP6 (Heiber et al., 2013), 1:500; rat anti-RFP (Chromotek), 1:2000; rabbit anti-SBP1 C-terminus (Mesen-Ramirez et al., 2016), 1:5000; rabbit anti-SBP1 N-terminus (Mesen-Ramirez et al., 2016), 1:4000, rabbit anti-BIP (Struck et al. 2005), 1:2000; rabbit anti-REX1 (Mesen-Ramirez et al., 2016), 1:20000. Secondary antibodies were horseradish peroxidase–conjugated goat anti-rat (Dianova), 1:3000, goat anti-mouse (Dianova), 1:3000 and donkey anti-rabbit (Dianova), 1:2500. For the detection of biotinylated proteins horseradish peroxidase-conjugated streptavidin (Invitrogen) was used at a 1:5000 dilution. Detection was performed with ECL (Bio-Rad or Thermo), and images were acquired with a ChemiDoc XRS imaging system (Bio-Rad).

### Proximity dependent biotin labelling of parasite proteins for mass spectrometry

For biotin labelling of proteins in live *P. falciparum* parasite infected RBCs, 100 ml of parasite culture of each cell line was grown to 5%-10% parasitemia in T175 cell culture flasks (Sarstedt). Before harvesting the cells were incubated with 20 mM biotin for 48 hours, with one medium change after 24 hours. Subsequently, the parasite culture was pelleted by centrifugation (2000g for 5 min) and the pellet resuspended in 20 ml RPMI complete medium. Infected red blood cells containing trophozoites and schizonts were purified by magnetic purification using MACS separation columns (Miltenyi Biotech). The purified infected RBCs were pelleted, transferred to a 1.5 ml reaction tube (Eppendorf) and washed twice with PBS. Cells were lysed in 1 ml lysis buffer (50 ml Tris-HCl pH 7.5, 500 mM NaCl, 1% TritonX-100, 1 mM DTT, 2x protease inhibitor cocktail (Roche), 1 mM PMSF). Lysates were frozen at −80 °C and thawed twice, to aid protein extraction. After centrifugation at 16000g for 10 minutes, supernatants were transferred to 2 ml reaction tubes (Eppendorf) and diluted 1:1 with 1 ml of 50 mM Tris-HCl pH 7.5. The lysate was added to 50 µl Streptavidin-sepharose beads and incubated for 16 h at 4°C with gentle head over tail mixing. The sepharose beads were pelleted by centrifugation at 1600g for 1 min, washed twice with lysis buffer, once with dH_2_O, twice with 50 mM Tris-HCl pH 7.5 and three times with 100 mM Triethylammonium bicarbonate buffer (TEAB, Sigma-Aldrich) pH 8.5. Finally, the Sepharose beads were resuspended in 50 µl 100 mM TEAB and shipped on ice for further processing.

### Quantitative MS of the biotinylated proteins

On-bead digestion of the protein on the streptavidin beads was performed as in (Hubner et al., 2015): 2.5 µl of 100 mM Tris(2-carboethyl)phosphine hydrochloride (TCEP, Sigma) was added to a final concentration of 5 mM to the beads and disulfide bonds were reduced for 1 hour while shaking in a thermoshaker at 37°C to prevent settling of the beads. Cysteine alkylation was induced by addition of methyl methanethiolsulfonate (MMTS, Thermo Scientific) in 100% isopropanol to obtain a final concentration of 10 mM followed by a 10 min incubation in a thermoshaker at 37°C. Proteins were subsequently digested off the beads by addition of 0.4 µg Trypsin/LysC (Promega, #V5072) for 1 h while shaking in a thermoshaker at 37°C. Beads were pelleted by 2 min centrifugation at 400g at room temperature and 53.5 µl of supernatant was transferred to a clean eppendorf tube. 50 µl of 100 mM TEAB was added to the beads followed by a 5 min incubation while shaking at 37°C, after which the beads were again pelleted by 2 min centrifugation at 400g at RT. 50 µl of supernatant was removed from the beads and pooled with the first supernatant to improve the efficiency of protein isolation. Protein digestion was continued overnight (∼17-20 hours) in a 37°C waterbath. Di-methyl labelling was performed as in (Boersema et al., 2009) as follows: 4µl of a 4% label (“light”: CH_2_O, 37% formaldehyde solution, Sigma-Aldrich; “medium”: CD_2_O, 20% formaldehyde, d2 solution, Sigma-Aldrich; “heavy”: ^13^CD_2_O, 20% formaldehyde-^13^C, d2 solution, Sigma-Aldrich, diluted to 4% in mass-spec grade MQ prior to use) was added to each 103.5 µl sample, followed by addition of 4 µl 0.6 M NaBH_3_CN (Merck) to the “light” and “medium” or 4 µl of 0.6 M NaBD_3_CN (Sigma) to the “heavy” reactions. Samples were incubated for 1 h while shaking at room temperature, after which the reaction was stopped by addition of 16 µl 1% ammonia. For each experiment “light”-, “medium”- and “heavy”-labelled samples were pooled into a single tube and the sample pool was acidified by addition of 15 µl 100% trifluoroacetic acid (TFA, Biosolve). Each sample-pool was cleaned and concentrated divided over two 3-disk-C18 stage-tips (Rappsilber et al., 2007) and stored at 4°C until loading for mass spectrometry analysis. Di-methyllabelling of replicate samples was performed under label-swap conditions as follows. REX3^trunc^-MAD-BirA*-GFP (REX3-MAD) reactions were labelled with a “heavy” label in experiment 1 and 3 and a “medium” label in experiment 2 and 4; REX3^trunc^-BirA*-GFP (REX3) reactions were labelled with a “medium” label in experiment 1 and 3 and a “light” label in experiment 2 and 4; STEVOR^1-260^-BirA*-GFP (STEVOR) reactions were labelled with a “light” label in experiment 1 and 3 and a “heavy” label in experiment 2 and 4.

### Mass spectrometry (MS) and MS data analysis

For each sample pool, one loaded 3-disk-C18 stage-tip was rehydrated with 25 μl buffer A (0.1% formic acid) and peptides were eluted using 30 μl buffer B (80% acetonitrile, 0.1% formic acid) in PCR tubes. Acetonitrile evaporation was achieved by 15 min vacuum spin at room temperature after which each sample was reconstituted to 12μl with buffer A. 5 µl of reconstituted sample (∼21% of total sample) was separated over a 30 cm C18-reverse phase column (1.8 μm Reprosil-Pur C18-AQ, dr. Maisch) and eluted using an Easy-nLC 1000 (Thermo Scientific) over a 94 min gradient (5.6% acetonitrile/0.1% formic acid - 25.6% acetonitrile/0.1% formic acid). Eluted peptides were directly injected into a QExactive mass spectrometer (Thermo Scientific). Data was acquired in TOP10 data-dependent acquisition mode with dynamic exclusion enabled for 20 s. Resolution for MS was set at 70.000 at m/z = 400 and for MS/MS at 17.5000.

Raw mass spectra were analyzed as in (Kensche et al., 2016) with modifications as follows: Xcalibur raw files were processes using MaxQuant (version 1.5.3.30 (Cox and Mann, 2008)) set to default parameters unless indicated. Multiplicity was set at 3, with an added mass of 28.03Da (“light”-), 32.06Da (“medium”-) or 36.08Da (“heavy”-label) to all lysine residues and peptide N-termini. Trypsin/P was set as the specific digestion mode with maximum 2 missed cleavages and a mass of 45.99Da (MMTS) was set as fixed modification of cysteine residues. Match-between-runs and re-quantify options were enabled with default parameters and iBAQ values were calculated. Mass spectra were compared to peptide masses from the *Plasmodium falciparum* 3D7 annotated proteome (PlasmoDB release 33) with the entire human proteome included in the contaminants list using the integrated Andromeda search engine. Default search settings (mass tolerance at 4.5 ppm for precursor ions and 20 ppm for fragment ions) were enabled, and peptides and proteins were accepted with a 0.01 FDR cut-off. Protein quantification required minimally two “unique + razor” peptide-ratios.

The MaxQuant output ProteinGroup file was further analyzed using the Perseus software package (version 1.4.0.20 (Tyanova et al., 2016)). M/L-, H/L- and H/M- (normalized) ratios were log2 transformed while intensity values were log10 transformed. A –x transformation was applied to experiment 1 and 3 M/L- (normalized) ratios and experiment 2 and 4 H/M- (normalized) ratios to obtain REX3-MAD-over-STEVOR, REX3-MAD-over-REX3 and STEVOR-over-REX3 ratios. After filtering on ‘only identified by site’, ‘reverse’ and ‘potential contaminant’ hits 490 of the original 674 protein identifications remained. Significant-outliers were determined using the intensity-based Significance B option (two-sided Benjamini-Hochberg test) with a FDR cut-off set to 0.05. Further processing of the file to Table S1 was performed in Excel.

Scatterplots of experiment 1 *vs* experiment 2 and experiment 3 *vs* experiment 4 were generated in R (R i386 3.3.3 (R_Core_Team, 2017)), where only normalized log2-ratios of protein identifications based on at least two peptides, of which minimally one unique, were included and significantly enriched (numbered) or depleted (lettered). Outliers were labelled according to the experiment with the least stringent FDR subdivided into the following classes: <5%, <1%, <0.5%, <0.1%, <5e^-4^, <1e^-4^, <5e^-5^, <1e^-5^. The heatmap was generated using the web-based Morpheus tool from the Broad Institute (Harvard, 2017), plotting the normalized log2-ratios of all four replicates each of REX3-MAD-over-STEVOR (MAD interactome over Maurer’s cleft proteome), REX3-MAD-over-REX3 (MAD interactome over host cell cytosolic proteome) and STEVOR-over-REX3 (Maurer’s cleft proteome over host cell cytosolic proteome) comparisons. Only proteins enriched with an FDR <5% in at least 2 out of 4 replicate reactions for any comparison were included. Proteins were ranked from high to low on the average normalized log2 ratio of the REX3-MAD-over-STEVOR condition and color intensity (red enriched, yellow neutral, blue depleted) was based on the 1-99 percentile of values for all identified proteins. NaN values - where either a protein not identified in an experiment or no ratio can be given due to a missing label - are depicted as grey blocks in the heatmap. PlasmoDB gene identifiers and product descriptions were taken from release 33. Short names were assembled from PlasmoDB gene names or symbols and adjusted based on the product description if appropriate.

The mass spectrometry proteomics data was deposited to the ProteomeXchange Consortium (http://www.proteomexchange.org/) using the PRIDE partner repository (Vizcaino et al., 2013) with the dataset identifier PXD008208.

### Transmission electron microscopy

Trophozoite-stage parasites were isolated by a percoll-gradient. After two washing steps with PBS, the cells were chemically fixed with 2.5% glutaraldehyde (Electron Microscopy Sciences) in 50 mM cacodylate buffer (pH 7.4) for 30-60 minutes. Thereafter, the cells were washed twice with 50 mM cacodylate buffer, post-fixed with 2% OsO4 (Electron Microscopy Sciences) in water and incubated for 40 minutes on ice in the dark. To remove the OsO4 solution, the cells were rinsed 3 times with water. The cells were contrasted with 0.5% uranylacetate (Electron Microscopy Sciences) and incubated for 30 minutes at room temperature, followed by dehydration using an ethanol series. The cell pellets were embedded in epoxy resin (EPON) (Roth) and incubated at 60°C for 24-72 h for polymerization. Samples from the solid epon blocks were sectioned into 55-65 nm thick slices using a Leica EM UC7 microtome. The slices were examined with a Tecnai Spirit transmission electron microscope (Thermo Fisher) at 80 kV.

### Binding assays

The initial binding assays (Fig. S10B) were carried out as follows. One day before the assay, Chinese Hamster Ovary CHO-745-ICAM-1 (Metwally et al., 2017) were seeded into T25 flasks to reach a confluency of 80-90% on the day of assay. On the day of assay, enrichment of trophozoites was done using 1% gelatine (Waterkeyn et al., 2001) and 5×10^6^ of these trophozoites were suspended in 5ml serum free RPMI medium. The trophozoite infected RBC suspension was then added to the CHO-745-ICAM-1 cells for one hour with 3 times shaking every 15 min. The non-bound red blood cells were then aspirated, and the flask was washed with 5 ml RPMI for at least 5 times, each time shaking 3 times. The cells were fixed with 1% glutaraldehyde and stained with Giemsa solution. From each flask 25 images were recorded at a 200x magnification using a Thermo Fisher EVOS. All the binding infected red blood cells were counted in the entire field (0.2 mm^2^) then the average was calculated from every 5 consecutive images.

The binding assays shown in Fig. 10E were a modification as described (Cronshagen et al., 2024): CHO cells expressing CD36, ICAM-1, or GFP (Metwally et al., 2017) were seeded into a 24-well plate containing coverslips 2 days (1 x 10^5^ cells/ml) or 3 days (2 x 10^5^ cells/ml) before the assay to obtain 0.5 ml/ volume per well. After knob selectin the cells were washed in binding medium (16,4 g/l RPMI-HEPES with 20 g/l glucose, pH 7.2), and the parasitemia (by Giemsa smear) and number of erythrocytes/ml determined using a Neubauer counting chamber) to adjust the cells to 2×10^6^ infected erythrocytes/ml. The CHO-cell containing wells were pre-incubated with binding medium for 30 min, the parasite suspension (triplicates of wells per condition) added and incubated for 60 min at 37°C with soft shaking every 15 min. Thereafter the coverslips were washed (6 times) by careful dunking into binding medium, each time removing excess medium using absorption with paper. After face-down incubation of coverslips in binding medium for 30 min, the cells on the coverslips were fixed (1% glutaraldehyde in 1x PBS, 30 min) and stained with Giemsa. The coverslips were washed in water and glued face-down onto glass slides using with CV-Mount (Leica). A Thermo Fisher EVOS xl (75 % light intensity at 40x magnification) was used to collect 15 images per parasite line and condition (5 images from each of the 3 technical replicates) for each independent experiment. Automated counting of bound cells was done using an established pipeline as described (Cronshagen et al., 2024).

