## Supplementary Figures with legends for "An unusual trafficking domain in MSRP6 defines a complex needed for Maurer’s clefts anchoring and maintenance in *P. falciparum* infected red blood cells"

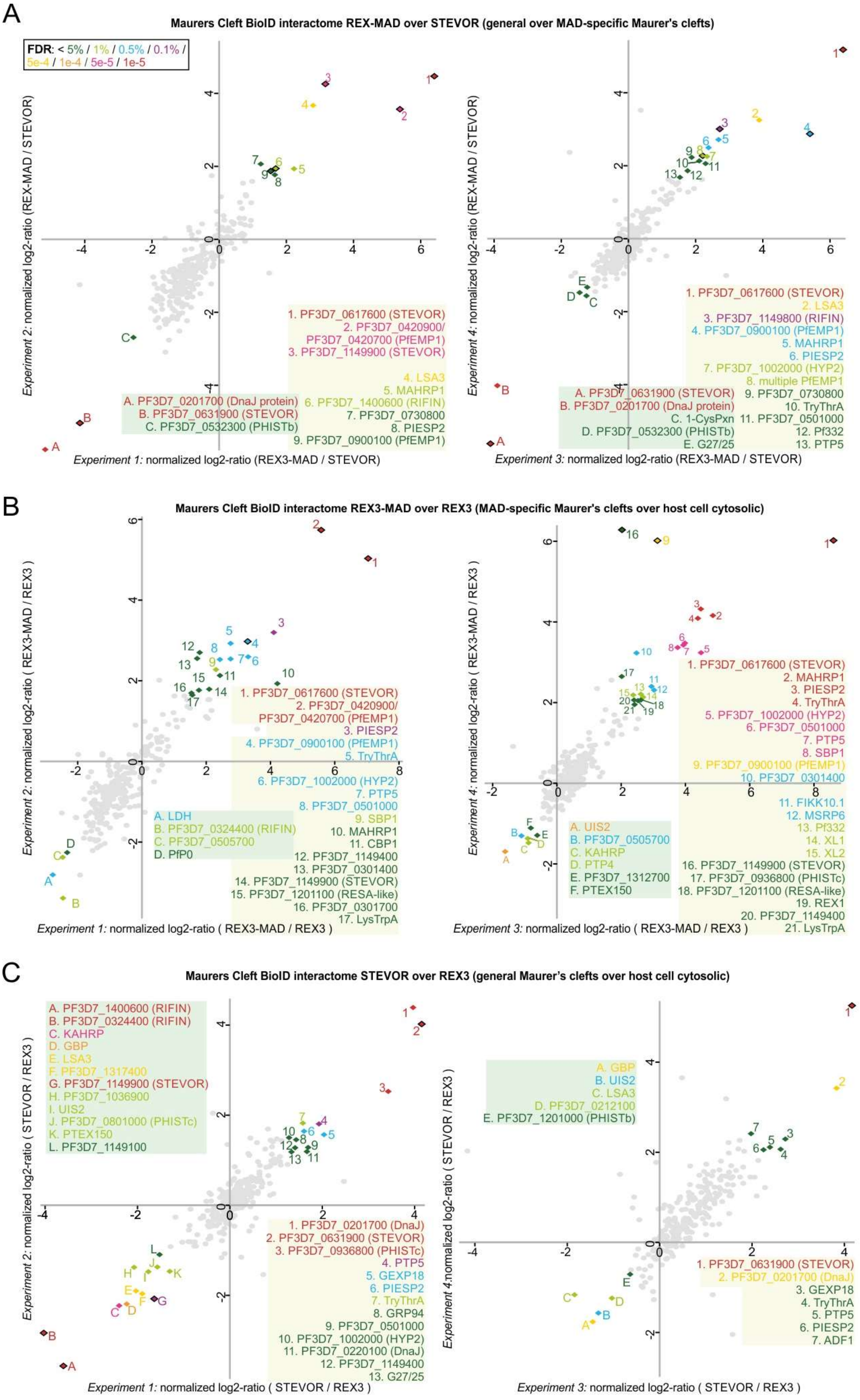


**Figure S1: Bio-ID scatterplots**

(**A-C**) Scatterplots of quantitative Bio-ID experiments of all four replicates (two experiments, each with 2 technical replicates, including the plots from Fig. 4 for side-by-side comparison of experiments) of the MSRP6 attachment domain (MAD) interactome compared to the STEVOR Maurer's cleft proteome (A), the

MAD interactome compared to the host cell cytosolic proteome (B), and the Maurer's cleft proteome compared to the host cytosolic proteome (C). Interactors of either REX3$^{trunc}$-MAD- (REX3-MAD), REX3$^{trunc}$-(REX3) or STEVOR$^{1-260}$- (STEVOR) BirA*-GFP fusion proteins were biotin labelled by BirA*, purified, Trypsin/LysC digested, dimethyl-labelled and light-medium-heavy labelled sample pools were measured by LC-MS/MS. Normalized ratios were calculated for proteins identified by at least two peptides and normalized log2-ratios of replicate experiments were plotted. Proteins enriched or depleted with an FDR below 5% in both replicates were labelled with the following colour code: FDR (in the least significant experiment) of <5% dark green, <1% light green, <0.5% blue, <0.1% purple, $<5e^{-4}$ yellow, $<1e^{-4}$ orange, $<5e^{-5}$ pink, $<1e^{-5}$ red. Significant proteins are numbered and gene-IDs or short unique names are given. Proteins belonging to variantly expressed multigene families (STEVOR, RIFIN, PfEMP1) that might differ in expression between the cell lines are likely false-positives and are shown as rhomboids with black frames in the plots. See also Figure 4 and Table S1.

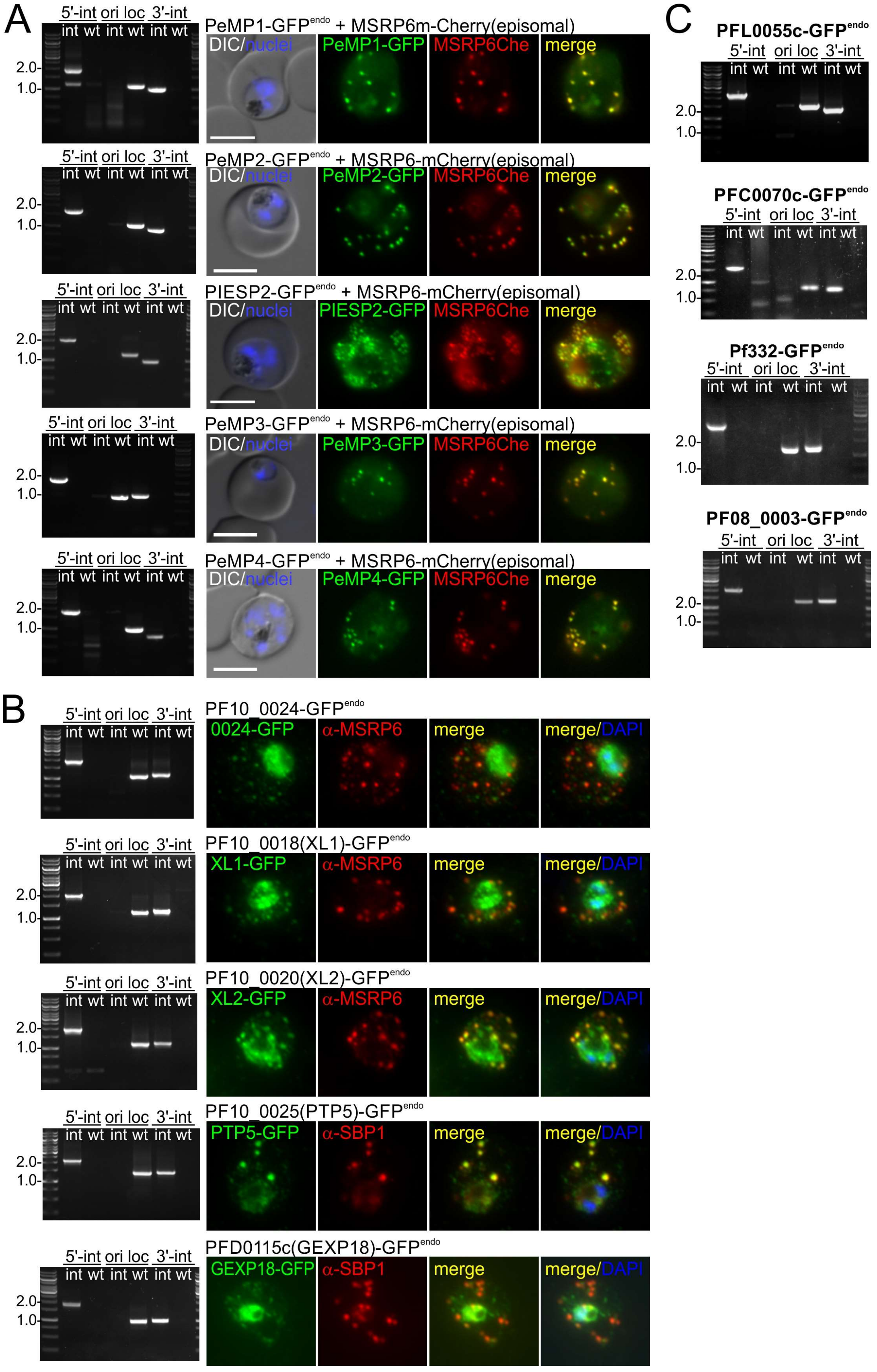


**Figure S2: Endogenous GFP tagging and Maurer's clefts localisation of MS hits**

(**A-C**) Agarose gels showing PCR products from genomic DNA to confirm correct integration of SLI plasmids to generate the indicated cell lines expressing endogenously GFP (and 2xFKBP) tagged target proteins. The primers used are listed in Table S3 and generate PCR product across the 5'- (5' int) and 3'- (3' int) prime integration junction or demonstrate absence of the original locus (ori-loc); Int, modified cell line; wt, 3D7 parental parasites. Marker is 1kb ladder with relevant bands marked in kbp. Fluorescence microscopy images show the signal of the corresponding endogenously GFP-tagged protein in live cells with an episomally co-expressed Maurer's clefts marker (MSRP6 or SBP1mCherry) (A) or IFA of acetone fixed cells detecting a Maurer's clefts marker (MSRP6 or SBP1) (B). No microscopy images are shown for the candidates in (C) but co-localised with a Maurer's clefts marker are found Figure S9B for Pf332-GFP$^{endo}$ or was done previously with an episomally expressed copy in the case of PF08_0003 (Heiber et al., 2013). Nuclei were stained with DAPI; DIC, differential interference contrast; merge, merge of red and green signal; scale bar, 5 µm.

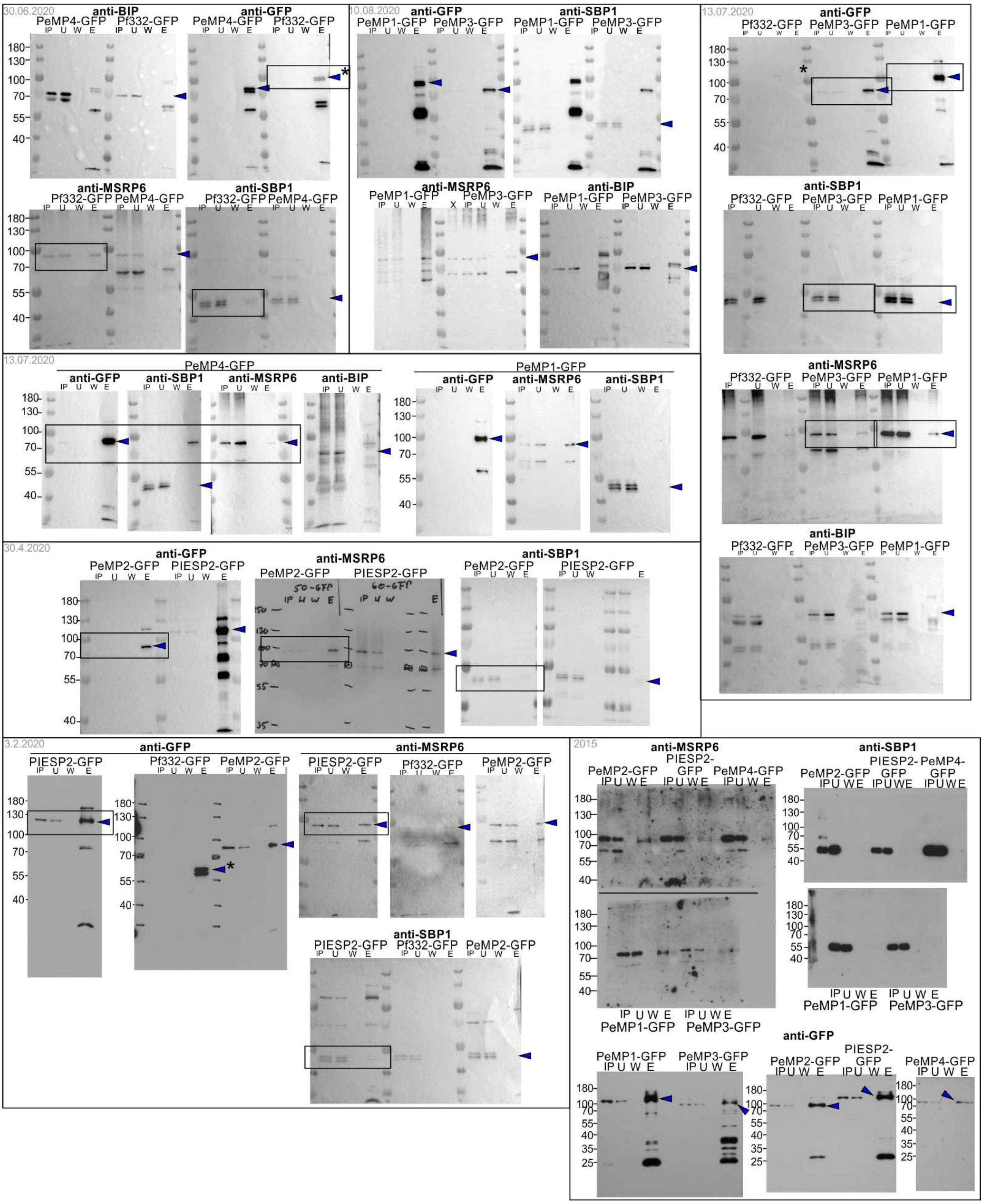


**Figure S3: Co-Immunoprecipitation blots**

Full blots of all co-immunoprecipitations using extracts from the cell lines with the indicated endogenously tagged proteins as baits are shown (used for quantification in Figure 5E). Blue arrows show relevant bands, boxes the areas shown in the main figure (Figure 5D). Asterisks indicate a degradation product of Pf332-GFP, the full length protein is too large to enter the gel. Smaller bands in eluates detected with anti-GFP show the frequently observed degradation products of GFP fusion proteins in *P. falciparum* parasites. Panels are labelled as in Figure 5D.

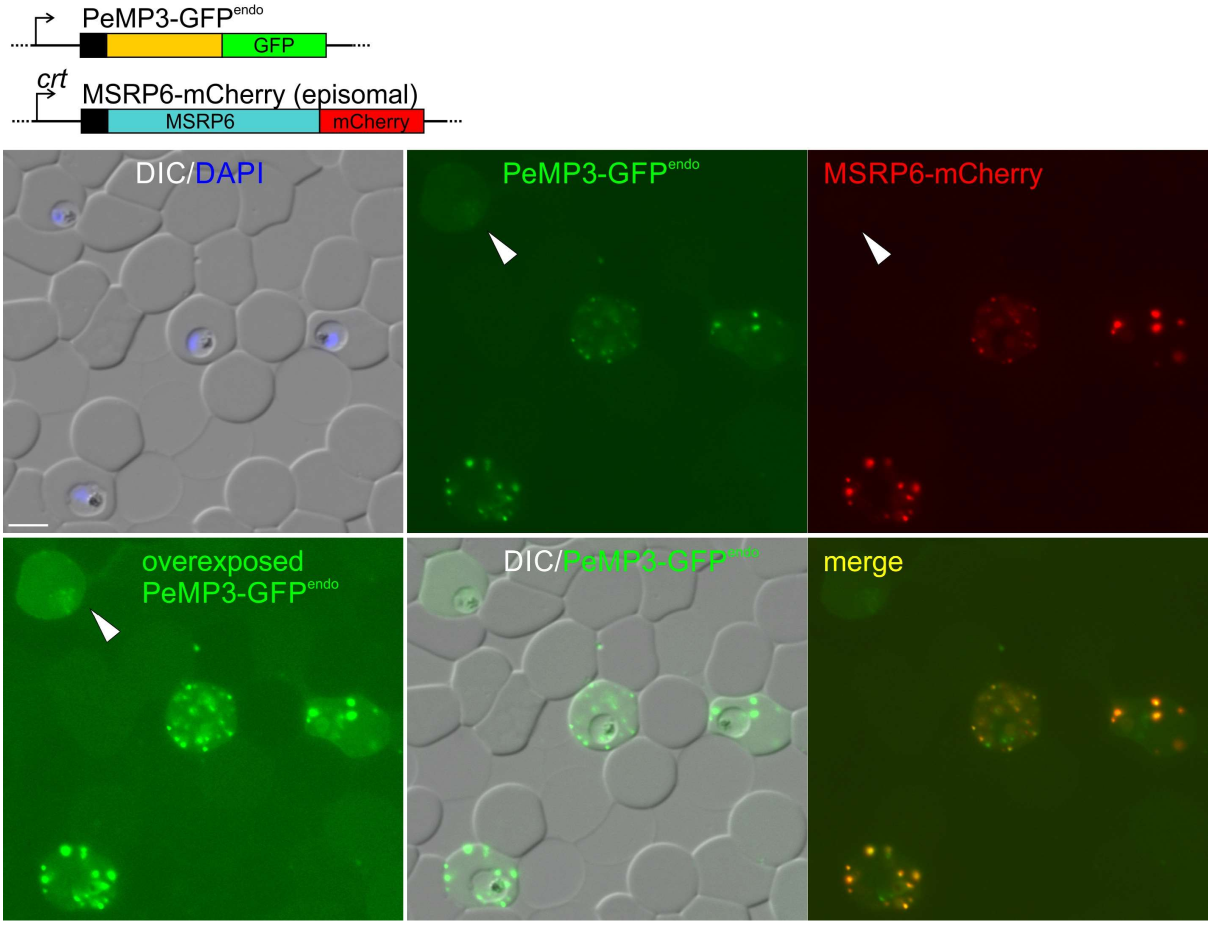


**Figure S4: Maurer's cleft recruitment of PeMP3 in young trophozoites that overexpress MSRP6**
Microscopy images of a larger area showing PeMP3-GFP$^{endo}$ young trophozoite parasites overexpressing MSRP6-mCherry. An example of the rare cells without MSRP6-mCherry (arrow, likely corresponding to cells lacking the episomal plasmid) showed little PeMP3 recruited to the Maurer's clefts even though the development stage was similar to the cells expressing MSRP6-mCherry. Nuclei were stained with DAPI; DIC, differential interference contrast; merge, merge of red and green signal; scale bar, 5 µm.

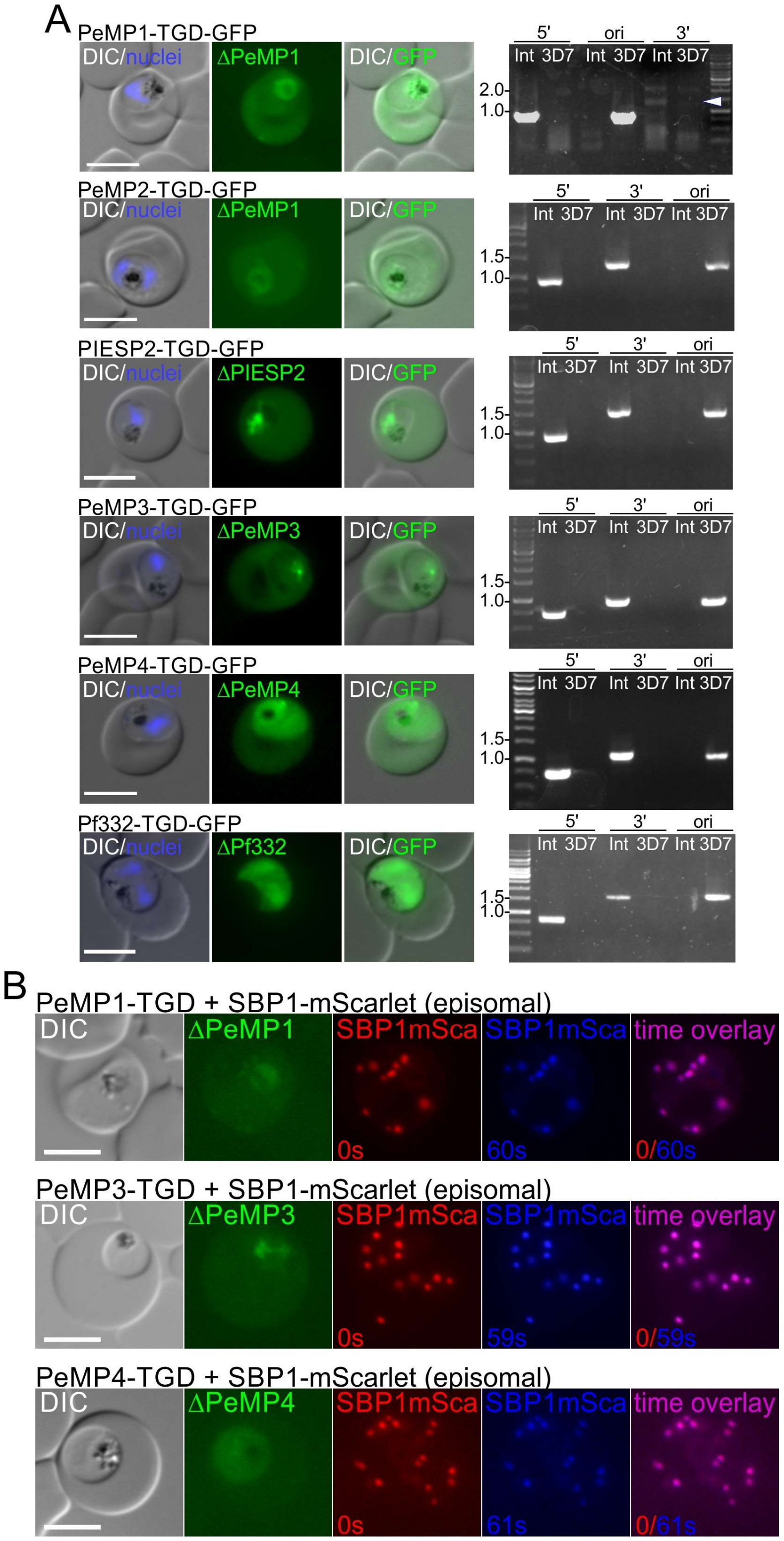


**Figure S5: Generation of TGD lines using SLI and cell lines showing no Maurer's clefts anchoring defect**
(**A**) Fluorescence microscopy images of live cells of parasites with disruptions in the indicated targets (targeted gene disruption, TGD; in image panels indicated by Δ). The disrupted product is fused with GFP, showing its loss of export or loss of Maurer's clefts localisation. Agarose gels show PCR products from genomic DNA, with primers (Table S3) that generate products across the 5'- (5' int) and 3'- (3' int) prime integration junction to show correct integration of the SLI plasmid or demonstrate absence of the original locus (ori-loc); Int, modified cell line; 3D7, wt parental parasites. Marker: 1kb ladder, relevant bands are marked in kbp. (**B**) Short-term time-lapse images of the indicated parasites co-expressing episomal SBP1-mScarlet. Labels as in Figure 6. DAPI, nuclei; DIC, differential interference contrast; scale bar, 5 µm.

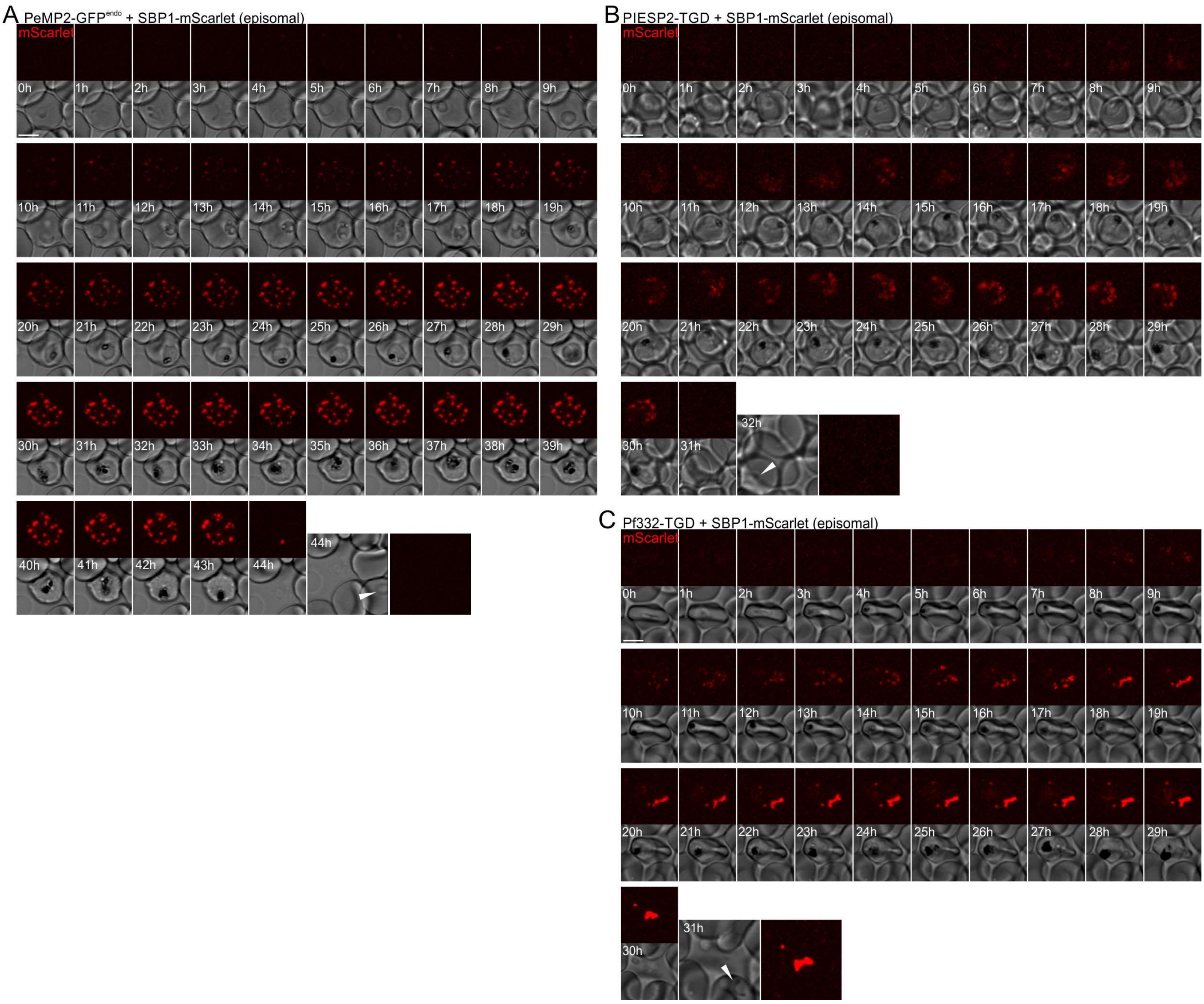


**Figure S6: Full time lapse data shown in Fig. 7E-G**

All time points of long term time lapse experiments in Fig. 7E-G showing PeMP2-GFP$^{endo}$ (**A**, control), PIESP2-TGD (**B**) and Pf332-TGD (**C**) parasites.

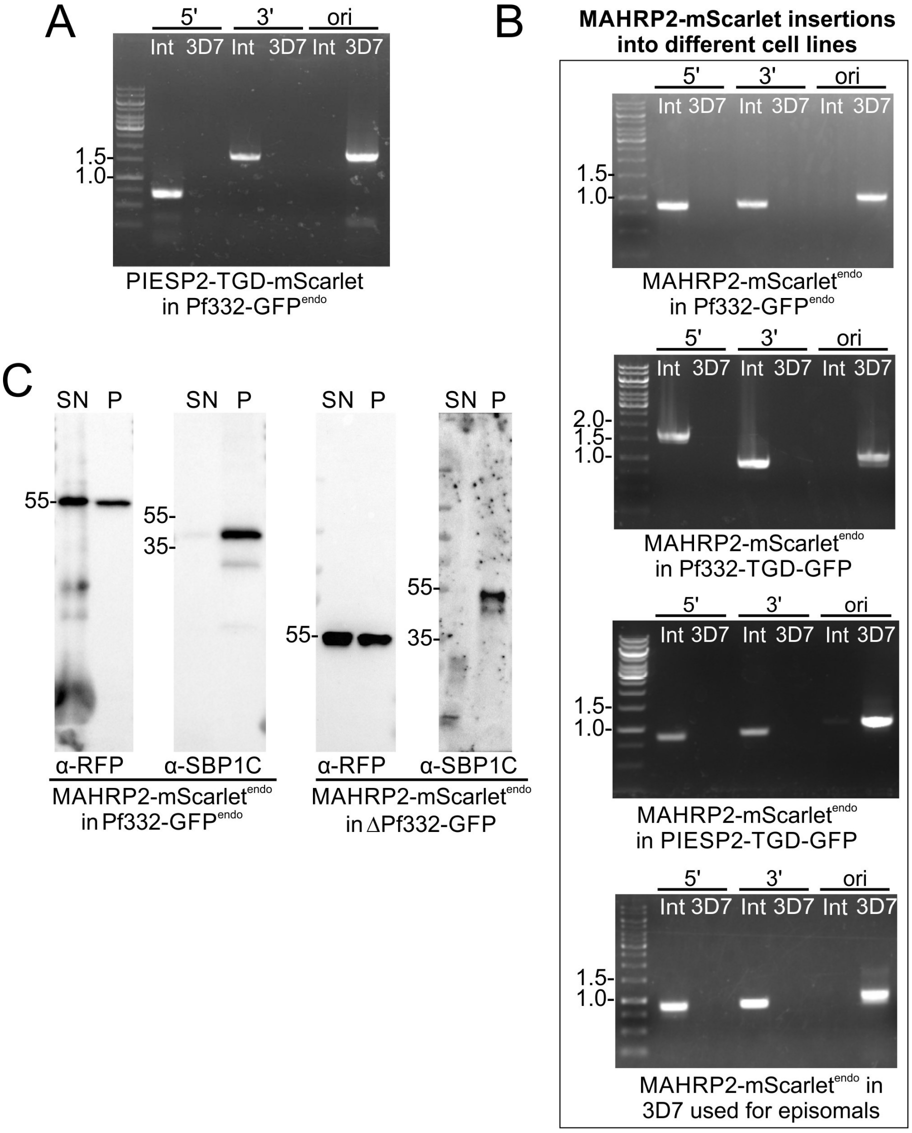


**Figure S7: Confirmation of correct plasmid insertion for double integration cell lines and MAHRP2 solubility**

(**A, B**) Agarose gels with PCR products confirming integration of plasmids into the genome of the Pf332-GFP$^{endo}$ parasites, resulting in disruption of the gene encoding PIESP2 (A) or into the cell lines indicated in (B), resulting in mScarlet tagging of endogenous MAHRP2 (MAHRP2-mScarlet$^{endo}$. Genomic DNA was used for the PCRs with primers (Table S3) that generate products across the 5'- (5') and 3'- (3') prime integration junction to show correct integration of the SLI plasmid or primers that demonstrate absence of the original locus (ori); Int, modified cell line; 3D7, wt parental parasites. Note that for the MAHRP2 integration into the Pf332-TGD line a different set of primers was used, resulting in a different banding patter. Marker: 1kb ladder, relevant bands are marked in kbp. (**C**) Western blots detecting the endogenously tagged MAHRP2-mScarlet in extracts separated into saponin soluble (host cell soluble pool) and pellet (includes the Maurer's clefts associated pool) fractions from the indicated cell lines. This experiment indicates that there are similar levels of free MAHRP2 before and after disruption of Pf332. SBP1 (SBP1C) was detected as Maurer's clefts associated control. Selected MW are indicated in kDa.

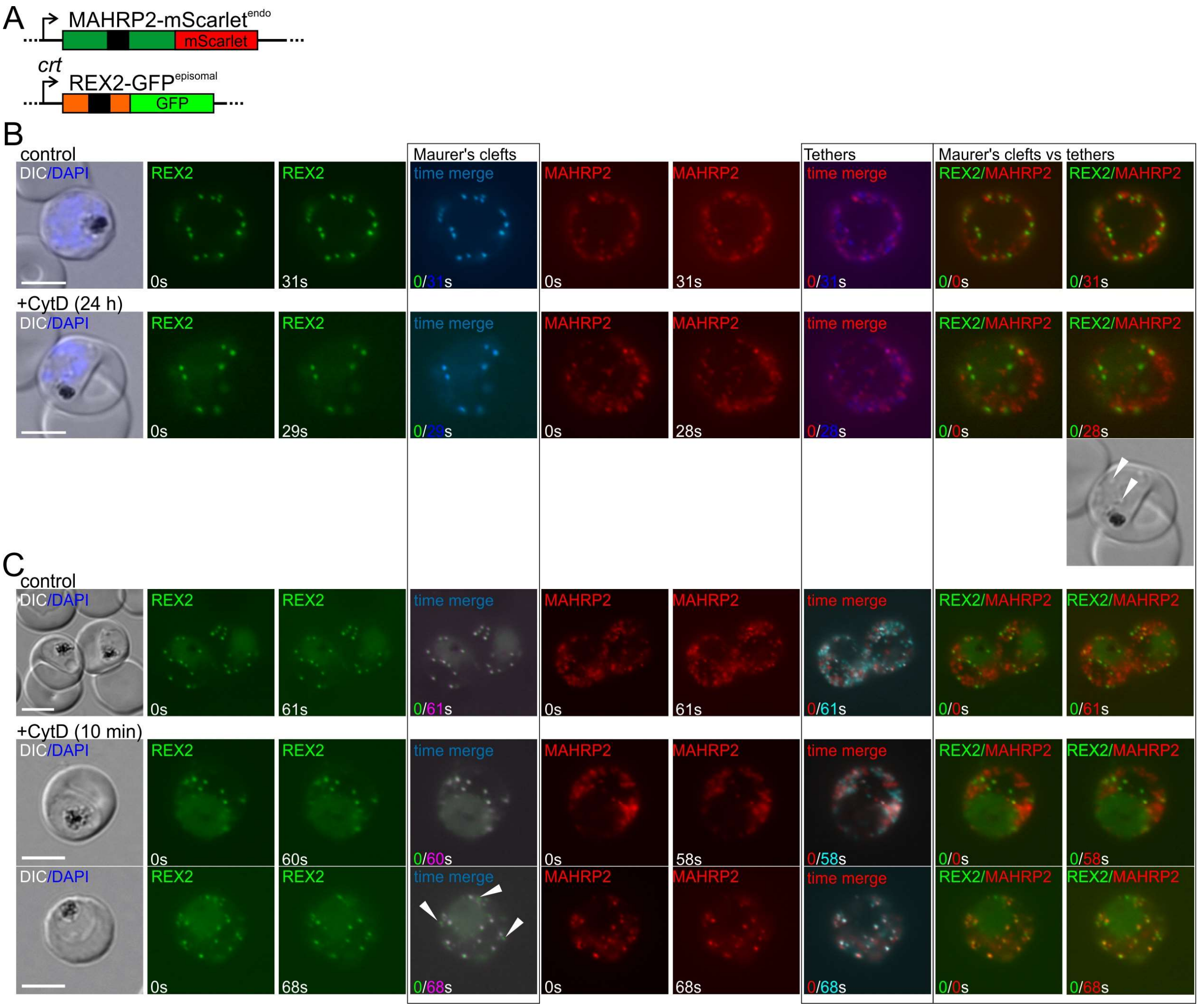


**Figure S8: Impact of cytochalasin D (CytD) on Maurer's clefts anchoring**

(**A**) Scheme of fluorescently tagged constructs in the cell line used for CytD experiments. Black box, hydrophobic region acting as transmembrane domain or signal peptide. (**B, C**) Fluorescence live microscopy images of the MAHRP2-mScarlet$^{endo}$ parasites expressing REX2-GFP (episomal) as a Maurer's clefts marker after long term (B, addition of CytD for 24 h starting in the ring stage before Maurer's clefts arrested) or short term (C, CytD treatment for 10 min). Images were taken after a short time interval to assess movement of Maurer's clefts (REX2 signal) and tethers (MAHRP2 signal) in time merges and overlays as indicated. Arrows in B show vesicles that had accumulated in the parasite, indicative of CytD effects. Controls and CytD treated cells are shown; two panels are shown for short term CytD treatment (C) to indicate the two phenotypes observed. Arrows in the time overlay in (C) highlight Maurer's clefts with evidence for movement. Scale bars, 5 µm.

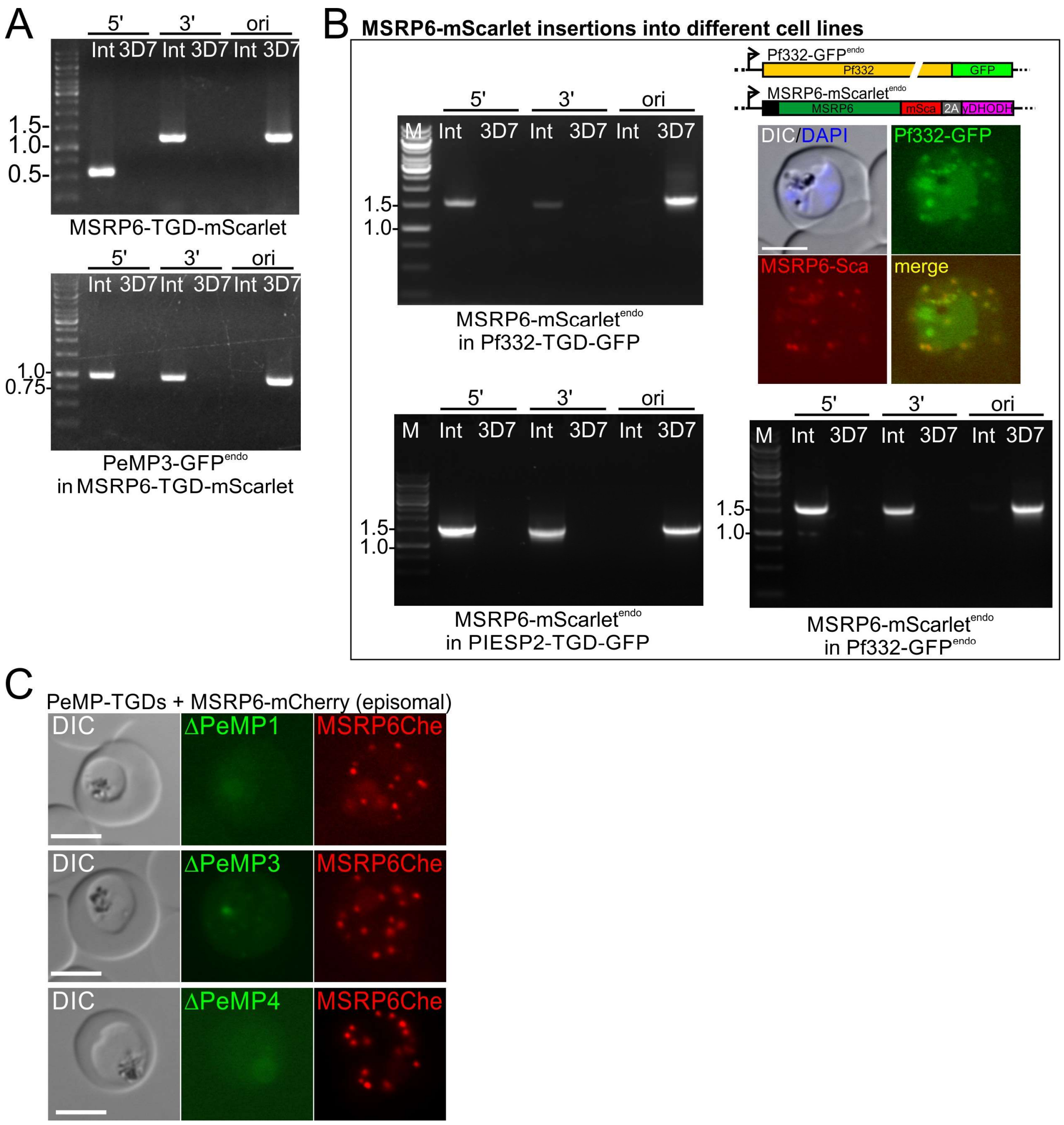


**Figure S9. Confirmation of MSRP6 integration lines and maintenance of MSRP6 at the Maurer's clefts in PeMP mutants.**

(**A, B**) Agarose gels with PCR products confirming plasmid integration that result in disruption of the gene encoding MSRP6 in 3D7 (top) or PeMP2-GFP$^{endo}$ (bottom) parasites (A) or integrations into MSRP6-mScarlet$^{endo}$ parasites (B) as indicated. Genomic DNA was used for the PCRs with primers (Table S3) that generate products across the 5'- (5') and 3'- (3') prime integration junction to show correct integration of the SLI plasmid or primers that demonstrate absence of the original locus (ori); Int, modified cell line; 3D7, wt parental parasites. Marker: 1kb ladder, relevant bands are marked in kbp. Image panels in B show fluorescence microscopy images of live cells expressing endogenously tagged Pf332 (with GFP) and MSRP6 (with mScarlet, MSRP6-mSca). (**C**) Episomally expressed MSRP6-mCherry (MSRP6Ch) in parasites with disrupted PeMPs is still Maurer's clefts associated. DAPI, nuclei; DIC, differential interference contrast; merge, merge of red and green channel; scale bar, 5 µm. TGD, targeted gene disruption (indicated as Δ in figure panels).

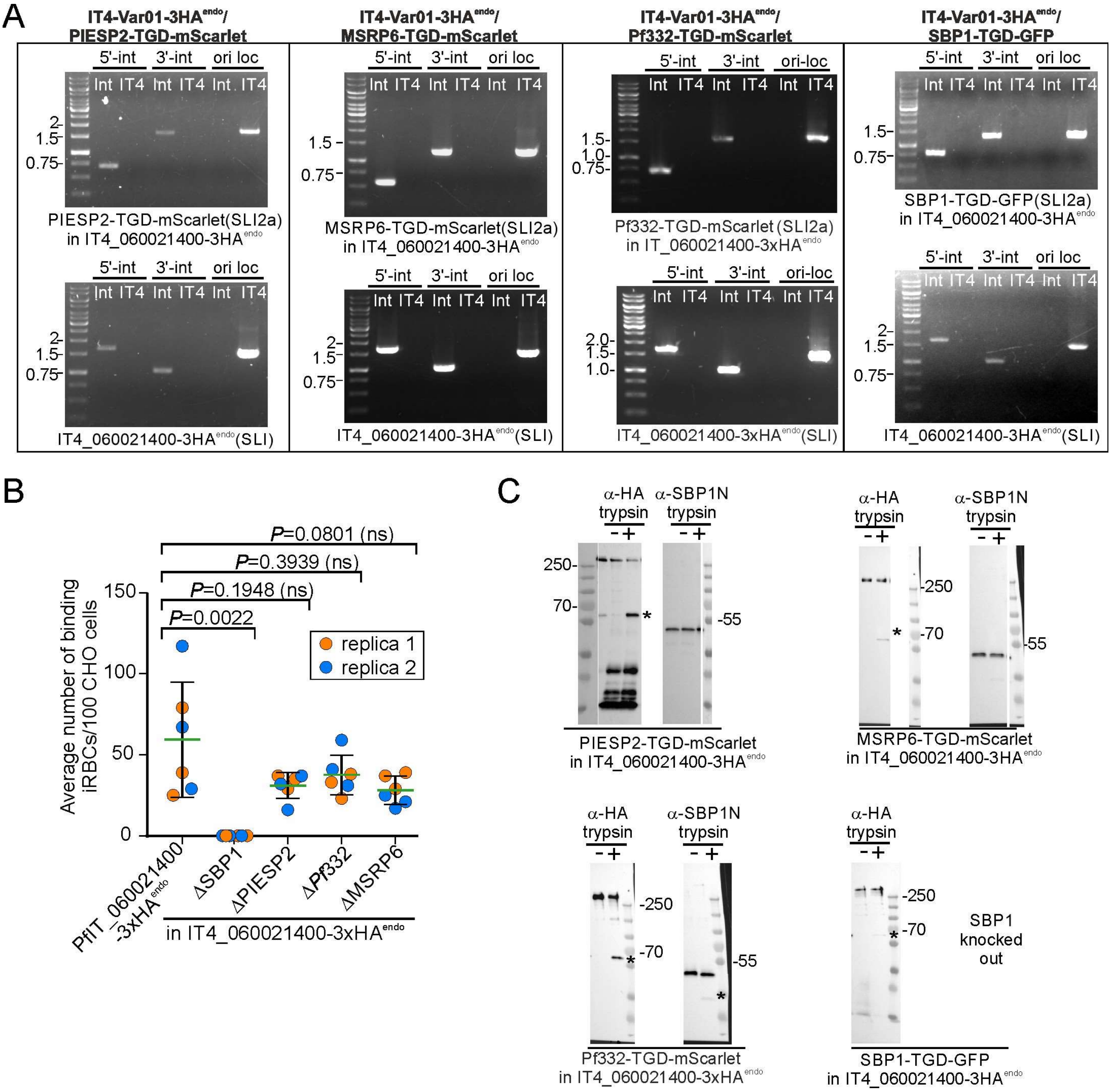


**Figure S10. Generation of TGDs in IT4var01-HA$^{act}$ parasites, initial binding experiments and replicas of immunoblots showing PfEMP1 surface exposure**

(**A**) Agarose gels show PCRs from genomic DNA with primers (Table S3) that generate products across the 5'- (5' int) and 3'- (3' int) integration junction to show correct insertion of the SLI plasmid into the genome and demonstrate absence of the original locus (ori-loc); Int, modified cell line; IT4, wt parental parasites. Marker: 1kb ladder, relevant bands marked in kbp. The top gel shows the targeted gene disruption (TGD) of the indicated gene product using the SLI2a plasmid. The bottom gel shows maintenance of the correct integration to activate and HA-tag IT4-Var01 (IT4_060021400) after the second integration had been obtained. (**B**) Initial binding experiments with iRBCs of the indicated cell lines to ICAM1 expressing CHO cells. Two experiments with technical replicas. Error bars (SD), mean (green lines) and P-values (Mann-Whitney-test) are shown; ns, not significant. (**C**) Replicas of blots shown in Figure 10D. Labelling as in Figure 10. Molecular weight standard is shown with relevant bands marked in kDa.
