## Supplementary material for "An unusual trafficking domain in MSRP6 defines a complex needed for Maurer’s clefts anchoring and maintenance in *P. falciparum* infected red blood cells": Data S1 (plasmids)

Constructs for Export domains (episomal expression)

Restriction site

GFP

>pArl1-crt-**Insert**-GFP

ggtacc**Insert**cctaggatgagtaaaggagaagaacttttcactggagttgtcccaattcttgttgaattagatggtgatgttaatgggcacaaattttctgtcagtggagagggtgaaggtgatgcaacatacggaaaacttacccttaaatttatttgcactactggaaaactaccagttccatggccaacacttgtcactactttcgcgtatggtcttcaatgctttgcgagatacccagatcatatgaaacagcatgactttttcaagagtgccatgcccgaaggttatgtacaggaaagaactatatttttcaaagatgacgggaactacaagacacgtgctgaagtcaagtttgaaggtgatacccttgttaatagaatcgagttaaaaggtattgattttaaagaagatggaaacattcttggacacaaattggaatacaactataactcacacaatgtatacatcatggcagacaaacaaaagaatggaatcaaagttaacttcaaaattagacacaacattgaagatggaagcgttcaactagcagaccattatcaacaaaatactccaattggcgatggccctgtccttttaccagacaaccattacctgtccacacaatctgccctttcgaaagatcccaacgaaaagagagaccacatggtccttcttgagtttgtaacagctgctgggattacacatggcatggatgagctctacaaaTAActcgagggatatggcagcttaatgttcgtttttcttatttatatatttataccaattgattgtatttataactgtaaaaatgtgtatgttgtgtgcatatttttttttgtgcatgcacatgcatgtaaatagctaaaattatgaacattttattttttgttcagaaaaaaaaaactttacacacataaaatggctagtatgaatagccatattttatataaattaaatcctatgaatttatgaccatattaaaaatttagatatttatggaacataatatgtttgaaacaataagacaaaattattattattattattatttttactgttataattatgtgtctccttcaatgattcataaatagttggacttgatttttaaaatgtttataatatgattagcatagttaaataaaaaaagttgaaaaattaaaaaaaaacatataaacacaaatgatggtttttccttcaatttcgatatcaatttatagaaacaaaatatatacttgtataattttatttttttatataaatcattacatatataattatacaatattttttctaagagataattatatattaatatatataaaaaaaggtgttttttttttttttttttatttttatttttattttatggtaatattttattttccttattttataaattatattagtttatatgtgattaattttatatattatcaatttatatatttttaaatgcttacttaattatctttttttttttttttttttttttttcccctctttttatattaatttatttttgaaaaaattgatatatatatatatatataatatatatatatacatgtagtagtattaaacaatgtataatatatataaataatatatttatatatttcatttcaattttaattttttttggttttttttttttttctttttgtcatatttaaaaaaaattatattcatataagttatgcattttttataaacattattcaatatatgtataatataatatatatatatatattaatgtattattccaatgtgcatgataaaagaaaaaaataatatttataaaaaaaaagaaaaataaaacaaaaaaagaaaaaaaaaaaaaaaaaaaaaaaaatacaaaaataaataatataatttataattatatattcttgtcacaataaaaatatatatatatatatatatatttataatatgtatattttaaactagaaaaggaataactaatattttatttattatcattcaagatttatattttataataataaatacctaatagaaatatatcaggatccatgcatggttcgctaaactgcatcgtcgctgtgtcccagaacatgggcatcggcaagaacggggactacccctggccaccgctcaggaacgaatttagatatttccagagaatgaccacaacctcttcagtagaaggtaaacagaatctggtgattatgggtaagaagacctggttctccattcctgagaagaatcgacctttaaagggtagaattaatttagttctcagcagagaactcaaggaacctccacaaggagctcattttctttccagaagtctagatgatgccttaaaacttactgaacaaccagaattagcaaataaagtagacatggtctggatagttggtggcagttctgtttataaggaagccatgaatcacccaggccatcttaaactatttgtgacaaggatcatgcaagactttgaaagtgacacgttttttccagaaattgatttggagaaatataaacttctgccagaatacccaggtgttctctctgatgtccaggaggagaaaggcattaagtacaaatttgaagtatatgagaagaatgattaagcttatttaataatagattaaaaatattataaaaataaaaacataaacacagaaattacaaaaaaaatacatatgaattttttttttgtaatcttccttataaatatagaataatgaatcatataaaacatatcattattcatttatttacatttaaaattattgtttcagtatctttaatttattatgtatatataaaaataacttacaattttattaataaacaatatatgtttattaattcatgttttgtaatttatgggatagcgattttttttactgtctgtatttttcttttttaattatgttttaattgtattttatttttattattgttctttttatagtattattttaaaacaaaatgtattttctaagaacttataataataataaatataaattttaataaaaattatatttatcttttacaatatgaacataaagtacaacattaatatatagcttttaatatttttattcctaatcatgtaaatcttaaatttttctttttaaacatatgttaaatatttatttctcattatatataagaacatatttattaaatctagaattctatagtgagtcgtattacaattcactggccgtcgttttacaacgtcgtgactgggaaaaccctggcgttacccaacttaatcgccttgcagcacatccccctttcgccagctggcgtaatagcgaagaggcccgcaccgatcgcccttcccaacagttgcgcagcctgaatggcgaatggcgcctgatgcggtattttctccttacgcatctgtgcggtatttcacaccgcatatggtgcactctcagtacaatctgctctgatgccgcatagttaagccagccccgacacccgccaacacccgctgacgcgccctgacgggcttgtctgctcccggcatccgcttacagacaagctgtgaccgtctccgggagctgcatgtgtcagaggttttcaccgtcatcaccgaaacgcgcgagacgaaagggcctcgtgatacgcctatttttataggttaatgtcatgataataatggtttcttagacgtcaggtggcacttttcggggaaatgtgcgcggaacccctatttgtttatttttctaaatacattcaaatatgtatccgctcatgagacaataaccctgataaatgcttcaataatattgaaaaaggaagagtatgagtattcaacatttccgtgtcgcccttattcccttttttgcggcattttgccttcctgtttttgctcacccagaaacgctggtgaaagtaaaagatgctgaagatcagttgggtgcacgagtgggttacatcgaactggatctcaacagcggtaagatccttgagagttttcgccccgaagaacgttttccaatgatgagcacttttaaagttctgctatgtggcgcggtattatcccgtattgacgccgggcaagagcaactcggtcgccgcatacactattctcagaatgacttggttgagtactcaccagtcacagaaaagcatcttacggatggcatgacagtaagagaattatgcagtgctgccataaccatgagtgataacactgcggccaacttacttctgacaacgatcggaggaccgaaggagctaaccgcttttttgcacaacatgggggatcatgtaactcgccttgatcgttgggaaccggagctgaatgaagccataccaaacgacgagcgtgacaccacgatgcctgtagcaatgccaacaacgttgcgcaaactattaactggcgaactacttactctagcttcccggcaacaattaatagactggatggaggcggataaagttgcaggaccacttctgcgctcggcccttccggctggctggtttattgctgataaatctggagccggtgagcgtgggtctcgcggtatcattgcagcactggggccagatggtaagccctcccgtatcgtagttatctacacgacggggagtcaggcaactatggatgaacgaaatagacagatcgctgagataggtgcctcactgattaagcattggtaactgtcagaccaagtttactcatatatactttagattgatttaaaacttcatttttaatttaaaaggatctaggtgaagatcctttttgataatctcatgaccaaaatcccttaacgtgagttttcgttccactgagcgtcagaccccgtagaaaagatcaaaggatcttcttgagatcctttttttctgcgcgtaatctgctgcttgcaaacaaaaaaaccaccgctaccagcggtggtttgtttgccggatcaagagctaccaactctttttccgaaggtaactggcttcagcagagcgcagataccaaatactgtccttctagtgtagccgtagttaggccaccacttcaagaactctgtagcaccgcctacatacctcgctctgctaatcctgttaccagtggctgctgccagtggcgataagtcgtgtcttaccgggttggactcaagacgatagttaccggataaggcgcagcggtcgggctgaacggggggttcgtgcacacagcccagcttggagcgaacgacctacaccgaactgagatacctacagcgtgagctatgagaaagcgccacgcttcccgaagggagaaaggcggacaggtatccggtaagcggcagggtcggaacaggagagcgcacgagggagcttccagggggaaacgcctggtatctttatagtcctgtcgggtttcgccacctctgacttgagcgtcgatttttgtgatgctcgtcaggggggcggagcctatcgaaaaacgccagcaacgcggcctttttacggttcctggccttttgctggccttttgctcacatgttctttcctgcgttatcccctgattctgtggataaccgtattaccgcctttgagtgagctgataccgctcgccgcagccgaacgaccgagcgcagcgagtcagtgagcgaggaagcggaagagcgcccaatacgcaaaccgcctctccccgcgcgttggccgattcattaatgcagctggcacgacaggtttcccgactggaaagcgggcagtgagcgcaacgcaattaatgtgagttagctcactcattaggcaccccaggctttacactttatgcttccggctcgtatgttgtgtggaattgtgagcggataacaatttcacacaggaaacagctatgaccatgattacgccaagctatttaggtgacactatagaatactcgcggccgctaacgtaacagacttaggaggagatcttagtagttgagtgattctatatacatatacataaataaattacataatattataaattttttttatattatattagaatgtttactatataaaaataaatattttcttatatatttttttattttttcatatgaaaaaagaaattttttatatatttttttattatattttataaataaaaaagaacttatacctattattattattatatataaaaaatatatatttttttataaattcctatttttccgattatttatttattttttttttttattgtttaaaaatatataaaaaaaattcttatatatttaatattataagtccataaatatatataatttatataatttatatatttccatactttttttattatatatcatgagataaaataaatttcatatagaaatcgtatttatttattaatttgatatatattaataaataaatattaataataatatgttatatatatatatatttatttaattattatataggaaactatatatatatatatatattatatttttttttttgaaatataataatatataatattattcctaaaaatatctatattcttatacctgaacccttttttttttttttttttttttttgactttcgattgttcattctgtttattgtataaatatataaatatatatatattgtatattttattacatatattttttttttttgaaagttaagaattttaacttaataaaaaaagtacatatatttttaataaatgtcctccattatataaattgttatataaaagattttatataatttaaatagaattcatttataaacaaatttgtttataaaaatatatttatgtatatataatataaatatatatatttatatatatatatatatatatattactatatatatttttttttttttttccttttttttactttcccaagttgtactgcttctaagcttttttaataaacatatataatttgtacaaatattttagattatatacatgatgtatatttgaatatattttctatatatttgtggttccatttttgtatattatatataatatatttatatatatattgatatgtcaatatttgtataacacatgaagtttttgtttttttttttttttttttttaatggagaatatttaataatatatgaaaaaaattttatataaatatatatatatatatatatatatatatatatatatatatatatatgtatatatatatatttatatatacatgtatgttttttaaaaagttaaataattctatagattattttcattgtcttcacatatatgacataaatattttaaaatcgacattccgatatattatatttttagactataatatccgttaataataaatacacgcagtcatattatttattatacattcatttattattttgttttttttaatttcttacatataactcgaccccgggat

**Inserts used:**

>PF08_0005

ATGAACATGTACGTAATCTATTACTACTTTTTAATTTTAATTTTTATAAATTCCTGCCATCTATACTTATCTTCCGAAAACCAAAAAAAAACTATTGCTACTGTTCATAATAACACAAGAACGAATACATTAAAAAAAAATAATAGTAATAATAATAATACAAATGATATATTTGGTTTAAGCCCCTCTGAAGTACCAAATTTAATAGATGATGAAGAAGAATATGAAATACTTGAAGGAGTCAAAAATAATTCTGATTTAACAAGATCAACTAACATAGACCACCCCTCTCCTTCTTCTACTATGATAGACCTTGATAATATAATTACAGAAATCTCAAAATTAAAAAAAAAAAAATTAAAAAAAGAAATGAACAATAAATTAGAGCAACAAACAAATCAAAATAACAATACTATTCATCATAATAATGAAAATGAAATAAATACCTTCACACAAAATATTAAAACAAATTCAAATGAACTCAAAAATCAAGATTCATCAAATAGTCTTATTAGTACTAATAGTAATACGATGGATGAATTATTGTTATACAGTACTAACTCAGAAGATAATTTAGATATTTCTTTTGGTGAACTTCAATTATATGAGAACAGTGATGAAGATACAAGTGATTATGAATATGTTAATGAAGATTATTCCGTGAATCATATTTTTTCAAATGATACAGAAGAATCATTTAATATTTTAGAAGATGTTGAAAATATATCATTATCATCAAGTAATAGATATCAATATAGTCCTATCGGACCATATAAAAAAAAACACACAAAGGTTTTTGAAAATTTTAAAATGACAAGAAATTATGACGAATTTCTTAAACTCTACAATTTAAAAGATTCTTCAGATAATCAAGAAGAATATTATGAATTGTTAGCAGGAGAACCTTTTAAACTTAATAGTTACTATTATAGAGATGTAAAGTATGAAAATGTTAAAAAATATATATTTAAAGAAATTTATGATAATATTCAAAATATAAGTACAGAAAATAAAATAATTGTATCAAAAAAAGAGGAACTCTTTTTTTATTTTATCAAAAGTTTATTGAAAAATAATTTTATATGCTTATCCTATGAAGAAGAGGAAAATTTATTGAGTGAATCTAAACTCTTACTAGAAGCTATGATATCTAAAAAAATACAA

>PF08_0005Δ1

ATGAACATGTACGTAATCTATTACTACTTTTTAATTTTAATTTTTATAAATTCCTGCCATCTAAATAATGAAAATGAAATAAATACCTTCACACAAAATATTAAAACAAATTCAAATGAACTCAAAAATCAAGATTCATCAAATAGTCTTATTAGTACTAATAGTAATACGATGGATGAATTATTGTTATACAGTACTAACTCAGAAGATAATTTAGATATTTCTTTTGGTGAACTTCAATTATATGAGAACAGTGATGAAGATACAAGTGATTATGAATATGTTAATGAAGATTATTCCGTGAATCATATTTTTTCAAATGATACAGAAGAATCATTTAATATTTTAGAAGATGTTGAAAATATATCATTATCATCAAGTAATAGATATCAATATAGTCCTATCGGACCATATAAAAAAAAACACACAAAGGTTTTTGAAAATTTTAAAATGACAAGAAATTATGACGAATTTCTTAAACTCTACAATTTAAAAGATTCTTCAGATAATCAAGAAGAATATTATGAATTGTTAGCAGGAGAACCTTTTAAACTTAATAGTTACTATTATAGAGATGTAAAGTATGAAAATGTTAAAAAATATATATTTAAAGAAATTTATGATAATATTCAAAATATAAGTACAGAAAATAAAATAATTGTATCAAAAAAAGAGGAACTCTTTTTTTATTTTATCAAAAGTTTATTGAAAAATAATTTTATATGCTTATCCTATGAAGAAGAGGAAAATTTATTGAGTGAATCTAAACTCTTACTAGAAGCTATGATATCTAAAAAAATACAA

>PF08_0005Δ2

ATGAACATGTACGTAATCTATTACTACTTTTTAATTTTAATTTTTATAAATTCCTGCCATCTATACTTATCTTCCGAAAACCAAAAAAAAACTATTGCTACTGTTCATAATAACACAAGAACGAATACATTAAAAAAAAATAATAGTAATAATAATAATACAAATGATATATTTGGTTTAAGCCCCTCTGAAGTACCAAATTTAATAGATGATGAAGAAGAATATGAAATACTTGAAGGAGTCAAAAATAATTCTGATTTAACAAGATCAACTAACATAGACCACCCCTCTCCTTCTTCTACTATGATAGACCTTGATAATATAATTACAGAAATCTCAAAATTAAAAAAAAAAAAATTAAAAAAAGAAATGAACAATAAATTAGAGCAACAAACAAATCAAAATAACAATACTATTCATCATGAAAATTTTAAAATGACAAGAAATTATGACGAATTTCTTAAACTCTACAATTTAAAAGATTCTTCAGATAATCAAGAAGAATATTATGAATTGTTAGCAGGAGAACCTTTTAAACTTAATAGTTACTATTATAGAGATGTAAAGTATGAAAATGTTAAAAAATATATATTTAAAGAAATTTATGATAATATTCAAAATATAAGTACAGAAAATAAAATAATTGTATCAAAAAAAGAGGAACTCTTTTTTTATTTTATCAAAAGTTTATTGAAAAATAATTTTATATGCTTATCCTATGAAGAAGAGGAAAATTTATTGAGTGAATCTAAACTCTTACTAGAAGCTATGATATCTAAAAAAATACAA

>PF08_0005Δ3

ATGAACATGTACGTAATCTATTACTACTTTTTAATTTTAATTTTTATAAATTCCTGCCATCTATACTTATCTTCCGAAAACCAAAAAAAAACTATTGCTACTGTTCATAATAACACAAGAACGAATACATTAAAAAAAAATAATAGTAATAATAATAATACAAATGATATATTTGGTTTAAGCCCCTCTGAAGTACCAAATTTAATAGATGATGAAGAAGAATATGAAATACTTGAAGGAGTCAAAAATAATTCTGATTTAACAAGATCAACTAACATAGACCACCCCTCTCCTTCTTCTACTATGATAGACCTTGATAATATAATTACAGAAATCTCAAAATTAAAAAAAAAAAAATTAAAAAAAGAAATGAACAATAAATTAGAGCAACAAACAAATCAAAATAACAATACTATTCATCATAATAATGAAAATGAAATAAATACCTTCACACAAAATATTAAAACAAATTCAAATGAACTCAAAAATCAAGATTCATCAAATAGTCTTATTAGTACTAATAGTAATACGATGGATGAATTATTGTTATACAGTACTAACTCAGAAGATAATTTAGATATTTCTTTTGGTGAACTTCAATTATATGAGAACAGTGATGAAGATACAAGTGATTATGAATATGTTAATGAAGATTATTCCGTGAATCATATTTTTTCAAATGATACAGAAGAATCATTTAATATTTTAGAAGATGTTGAAAATATATCATTATCATCAAGTAATAGATATCAATATAGTCCTATCGGACCATATAAAAAAAAACACACAAAGGTTTTT

>PF08_0005Δ1a

ATGAACATGTACGTAATCTATTACTACTTTTTAATTTTAATTTTTATAAATTCCTGCCATCTAAATAATAGTAATAATAATAATACAAATGATATATTTGGTTTAAGCCCCTCTGAAGTACCAAATTTAATAGATGATGAAGAAGAATATGAAATACTTGAAGGAGTCAAAAATAATTCTGATTTAACAAGATCAACTAACATAGACCACCCCTCTCCTTCTTCTACTATGATAGACCTTGATAATATAATTACAGAAATCTCAAAATTAAAAAAAAAAAAATTAAAAAAAGAAATGAACAATAAATTAGAGCAACAAACAAATCAAAATAACAATACTATTCATCATAATAATGAAAATGAAATAAATACCTTCACACAAAATATTAAAACAAATTCAAATGAACTCAAAAATCAAGATTCATCAAATAGTCTTATTAGTACTAATAGTAATACGATGGATGAATTATTGTTATACAGTACTAACTCAGAAGATAATTTAGATATTTCTTTTGGTGAACTTCAATTATATGAGAACAGTGATGAAGATACAAGTGATTATGAATATGTTAATGAAGATTATTCCGTGAATCATATTTTTTCAAATGATACAGAAGAATCATTTAATATTTTAGAAGATGTTGAAAATATATCATTATCATCAAGTAATAGATATCAATATAGTCCTATCGGACCATATAAAAAAAAACACACAAAGGTTTTTGAAAATTTTAAAATGACAAGAAATTATGACGAATTTCTTAAACTCTACAATTTAAAAGATTCTTCAGATAATCAAGAAGAATATTATGAATTGTTAGCAGGAGAACCTTTTAAACTTAATAGTTACTATTATAGAGATGTAAAGTATGAAAATGTTAAAAAATATATATTTAAAGAAATTTATGATAATATTCAAAATATAAGTACAGAAAATAAAATAATTGTATCAAAAAAAGAGGAACTCTTTTTTTATTTTATCAAAAGTTTATTGAAAAATAATTTTATATGCTTATCCTATGAAGAAGAGGAAAATTTATTGAGTGAATCTAAACTCTTACTAGAAGCTATGATATCTAAAAAAATACAA

>PF08_0005Δ1b

ATGAACATGTACGTAATCTATTACTACTTTTTAATTTTAATTTTTATAAATTCCTGCCATCTATACTTATCTTCCGAAAACCAAAAAAAAACTATTGCTACTGTTCATAATAACACAAGAACGAATACATTAAAAAAATCTCCTTCTTCTACTATGATAGACCTTGATAATATAATTACAGAAATCTCAAAATTAAAAAAAAAAAAATTAAAAAAAGAAATGAACAATAAATTAGAGCAACAAACAAATCAAAATAACAATACTATTCATCATAATAATGAAAATGAAATAAATACCTTCACACAAAATATTAAAACAAATTCAAATGAACTCAAAAATCAAGATTCATCAAATAGTCTTATTAGTACTAATAGTAATACGATGGATGAATTATTGTTATACAGTACTAACTCAGAAGATAATTTAGATATTTCTTTTGGTGAACTTCAATTATATGAGAACAGTGATGAAGATACAAGTGATTATGAATATGTTAATGAAGATTATTCCGTGAATCATATTTTTTCAAATGATACAGAAGAATCATTTAATATTTTAGAAGATGTTGAAAATATATCATTATCATCAAGTAATAGATATCAATATAGTCCTATCGGACCATATAAAAAAAAACACACAAAGGTTTTTGAAAATTTTAAAATGACAAGAAATTATGACGAATTTCTTAAACTCTACAATTTAAAAGATTCTTCAGATAATCAAGAAGAATATTATGAATTGTTAGCAGGAGAACCTTTTAAACTTAATAGTTACTATTATAGAGATGTAAAGTATGAAAATGTTAAAAAATATATATTTAAAGAAATTTATGATAATATTCAAAATATAAGTACAGAAAATAAAATAATTGTATCAAAAAAAGAGGAACTCTTTTTTTATTTTATCAAAAGTTTATTGAAAAATAATTTTATATGCTTATCCTATGAAGAAGAGGAAAATTTATTGAGTGAATCTAAACTCTTACTAGAAGCTATGATATCTAAAAAAATACAA

>PF08_0005Δ1c

ATGAACATGTACGTAATCTATTACTACTTTTTAATTTTAATTTTTATAAATTCCTGCCATCTATACTTATCTTCCGAAAACCAAAAAAAAACTATTGCTACTGTTCATAATAACACAAGAACGAATACATTAAAAAAAAATAATAGTAATAATAATAATACAAATGATATATTTGGTTTAAGCCCCTCTGAAGTACCAAATTTAATAGATGATGAAGAAGAATATGAAATACTTGAAGGAGTCAAAAATAATTCTGATTTAACAAGATCAACTAACATAGACCACCCCAATAATGAAAATGAAATAAATACCTTCACACAAAATATTAAAACAAATTCAAATGAACTCAAAAATCAAGATTCATCAAATAGTCTTATTAGTACTAATAGTAATACGATGGATGAATTATTGTTATACAGTACTAACTCAGAAGATAATTTAGATATTTCTTTTGGTGAACTTCAATTATATGAGAACAGTGATGAAGATACAAGTGATTATGAATATGTTAATGAAGATTATTCCGTGAATCATATTTTTTCAAATGATACAGAAGAATCATTTAATATTTTAGAAGATGTTGAAAATATATCATTATCATCAAGTAATAGATATCAATATAGTCCTATCGGACCATATAAAAAAAAACACACAAAGGTTTTTGAAAATTTTAAAATGACAAGAAATTATGACGAATTTCTTAAACTCTACAATTTAAAAGATTCTTCAGATAATCAAGAAGAATATTATGAATTGTTAGCAGGAGAACCTTTTAAACTTAATAGTTACTATTATAGAGATGTAAAGTATGAAAATGTTAAAAAATATATATTTAAAGAAATTTATGATAATATTCAAAATATAAGTACAGAAAATAAAATAATTGTATCAAAAAAAGAGGAACTCTTTTTTTATTTTATCAAAAGTTTATTGAAAAATAATTTTATATGCTTATCCTATGAAGAAGAGGAAAATTTATTGAGTGAATCTAAACTCTTACTAGAAGCTATGATATCTAAAAAAATACAA

>PF08_0005-1ab

ATGAACATGTACGTAATCTATTACTACTTTTTAATTTTAATTTTTATAAATTCCTGCCATCTATACTTATCTTCCGAAAACCAAAAAAAAACTATTGCTACTGTTCATAATAACACAAGAACGAATACATTAAAAAAAAATAATAGTAATAATAATAATACAAATGATATATTTGGTTTAAGCCCCTCTGAAGTACCAAATTTAATAGATGATGAAGAAGAATATGAAATACTTGAAGGAGTCAAAAATAATTCTGATTTAACAAGATCAACTAACATAGACCACCCCgctggtgcaggagccggtgctggtgcaggagccggtgccggt

>PF08_0005-1a

ATGAACATGTACGTAATCTATTACTACTTTTTAATTTTAATTTTTATAAATTCCTGCCATCTATACTTATCTTCCGAAAACCAAAAAAAAACTATTGCTACTGTTCATAATAACACAAGAACGAATACATTAAAAAAAgctggtgcaggagccggtgctggtgcaggagccggtgccggt

>MSRP6

ATGAAAAGCAAAAAAATAATATGTTCATCTTGCTTATTTTTAATATTTTTAAGTGTAATATTTTGTAGTGAACCAGATACAAATTCATTTGATGAAAATGTAAAGAAGAATGAAGTTTTTAATGCCTTAAATGAACATTTAGAAAGTATAAGTAATATCGTAAAAGTAAATATTATGGATGCCCTTTCAAATAACCCTTCGTTAATAAAAAAGACATACGAAGCTGTTGAAATAAATGATGATGATTATGTTCTGGAATATGTTGGGGATTCGACAGGAAAATATGGAGAGGGCTCTATATTCTATGATGAAAGTAAGAAATATAATTATAGAAAGTATTTAGATATAGAAAATGAATTACGAAATATGAAAGGTGAAGATGATGATGATTTTGATGAAGATGATGATGATTTTGATGAAGATGATGAAGATGATGAAGATGATGAAGATGATGAAGATGATGTAGATGATGAAGATGATCTAGATGTTGAAGATAATGTAGATGATGAATATGATGATGAGCATAATCATAATTATAATGATGACAAATTGAGTGAAAATCCTGAGAAGTATTCAAATTATAATAAAAATATACACGAAGATAAGAAAAAAGATAATTTGAATGAACCACATTTTAAACAAACTCATTATATATATTCATCAAATCACGATAATAATGAAACATCTAGATTTCCAAAAAAAAATGTACCTAAATATGATGAAAAATTAAATAATGAATTTAAAACATATTTAAGAAGACCTGAAAAGAAAGAAGAAACAAAAGAATATCCTAACAATGGATGTTCTGTAGTTCAAATATCTATAGTAACTAATGAAGATTTTTTAAAAAAAGTTAGAGAAAGAAATAAAAAGAGAAATAATAAAAAAAGAACAAATATTTATGATAGTGATGAAGAATCAGAAAGTTCAGAAGAAACAAGTAAAGATCCCTATTCATCGGGTCCATATACAGTTGATCATAAAAATGAAAATTGTTTTCCTTACAAGCAAAATTCAGATATTCATCCAAATATAAATAATGATAATAATGATAATAATGAAAATAATGATAATAATGAAAATAATGATAATAATATACCTGTTAATAATAAAAAAAATTCTCATACTCCTCATGTACCAAAAGATACCAAAAATAATATAGAAAAAAATCATAGTAACTATAATTACAAACCAATTAATCAAGAAAATTTATCAATGTATTTTAATAATAATAATAAGAAGAAGAATAAAAAGAATAGTCAAAAAAATAATCAAAAGAATAATCAAAAGAATAATCAAAAGCATAACCAAGATCTTTATAATATAAAAAAAAATAATGATCTACCTCAATATGAAGAATCTAATAAAACTTATAAAAAAATTTTTTCAAATGAACATGAACCACGTCCGATGATAACAACATCCAGATATGAAAATAAAATATCAAATGATAATAACAACAATAACTATAATTTGAATGAACAAGAAAGAAAACTTATTAAAAATATGATAGATATATTTATTTATACATACAAGTTAAATTATACAAACAGTAAATCTATATCTAAATTATTTAAAAATAATTTACTTAAAAAAAACTTCCGTACTTATTTCACAAATTATATATATACACTTTTTAATTATGGAAAGACATATAATTTTCTAACACCTTATAATAAAGATAATGATCATATGTATAGACAATTATTTGATGAAGCTGTACAAATGATGGATCTACTTATTAATAAAATGGACTTAGCTTTAAATCCCACGAAATTA

>MSRP6Δ1

ATGAAAAGCAAAAAAATAATATGTTCATCTTGCTTATTTTTAATATTTTTAAGTGTAATATTTTGTGTAAAAGTAAATATTATGGATGCCCTTTCAAATAACCCTTCGTTAATAAAAAAGACATACGAAGCTGTTGAAATAAATGATGATGATTATGTTCTGGAATATGTTGGGGATTCGACAGGAAAATATGGAGAGGGCTCTATATTCTATGATGAAAGTAAGAAATATAATTATAGAAAGTATTTAGATATAGAAAATGAATTACGAAATATGAAAGGTGAAGATGATGATGATTTTGATGAAGATGATGATGATTTTGATGAAGATGATGAAGATGATGAAGATGATGAAGATGATGAAGATGATGTAGATGATGAAGATGATCTAGATGTTGAAGATAATGTAGATGATGAATATGATGATGAGCATAATCATAATTATAATGATGACAAATTGAGTGAAAATCCTGAGAAGTATTCAAATTATAATAAAAATATACACGAAGATAAGAAAAAAGATAATTTGAATGAACCACATTTTAAACAAACTCATTATATATATTCATCAAATCACGATAATAATGAAACATCTAGATTTCCAAAAAAAAATGTACCTAAATATGATGAAAAATTAAATAATGAATTTAAAACATATTTAAGAAGACCTGAAAAGAAAGAAGAAACAAAAGAATATCCTAACAATGGATGTTCTGTAGTTCAAATATCTATAGTAACTAATGAAGATTTTTTAAAAAAAGTTAGAGAAAGAAATAAAAAGAGAAATAATAAAAAAAGAACAAATATTTATGATAGTGATGAAGAATCAGAAAGTTCAGAAGAAACAAGTAAAGATCCCTATTCATCGGGTCCATATACAGTTGATCATAAAAATGAAAATTGTTTTCCTTACAAGCAAAATTCAGATATTCATCCAAATATAAATAATGATAATAATGATAATAATGAAAATAATGATAATAATGAAAATAATGATAATAATATACCTGTTAATAATAAAAAAAATTCTCATACTCCTCATGTACCAAAAGATACCAAAAATAATATAGAAAAAAATCATAGTAACTATAATTACAAACCAATTAATCAAGAAAATTTATCAATGTATTTTAATAATAATAATAAGAAGAAGAATAAAAAGAATAGTCAAAAAAATAATCAAAAGAATAATCAAAAGAATAATCAAAAGCATAACCAAGATCTTTATAATATAAAAAAAAATAATGATCTACCTCAATATGAAGAATCTAATAAAACTTATAAAAAAATTTTTTCAAATGAACATGAACCACGTCCGATGATAACAACATCCAGATATGAAAATAAAATATCAAATGATAATAACAACAATAACTATAATTTGAATGAACAAGAAAGAAAACTTATTAAAAATATGATAGATATATTTATTTATACATACAAGTTAAATTATACAAACAGTAAATCTATATCTAAATTATTTAAAAATAATTTACTTAAAAAAAACTTCCGTACTTATTTCACAAATTATATATATACACTTTTTAATTATGGAAAGACATATAATTTTCTAACACCTTATAATAAAGATAATGATCATATGTATAGACAATTATTTGATGAAGCTGTACAAATGATGGATCTACTTATTAATAAAATGGACTTAGCTTTAAATCCCACGAAATTA

>MSRP6Δ2

ATGAAAAGCAAAAAAATAATATGTTCATCTTGCTTATTTTTAATATTTTTAAGTGTAATATTTTGTAGTGAACCAGATACAAATTCATTTGATGAAAATGTAAAGAAGAATGAAGTTTTTAATGCCTTAAATGAACATTTAGAAAGTATAAGTAATATCACAGTTGATCATAAAAATGAAAATTGTTTTCCTTACAAGCAAAATTCAGATATTCATCCAAATATAAATAATGATAATAATGATAATAATGAAAATAATGATAATAATGAAAATAATGATAATAATATACCTGTTAATAATAAAAAAAATTCTCATACTCCTCATGTACCAAAAGATACCAAAAATAATATAGAAAAAAATCATAGTAACTATAATTACAAACCAATTAATCAAGAAAATTTATCAATGTATTTTAATAATAATAATAAGAAGAAGAATAAAAAGAATAGTCAAAAAAATAATCAAAAGAATAATCAAAAGAATAATCAAAAGCATAACCAAGATCTTTATAATATAAAAAAAAATAATGATCTACCTCAATATGAAGAATCTAATAAAACTTATAAAAAAATTTTTTCAAATGAACATGAACCACGTCCGATGATAACAACATCCAGATATGAAAATAAAATATCAAATGATAATAACAACAATAACTATAATTTGAATGAACAAGAAAGAAAACTTATTAAAAATATGATAGATATATTTATTTATACATACAAGTTAAATTATACAAACAGTAAATCTATATCTAAATTATTTAAAAATAATTTACTTAAAAAAAACTTCCGTACTTATTTCACAAATTATATATATACACTTTTTAATTATGGAAAGACATATAATTTTCTAACACCTTATAATAAAGATAATGATCATATGTATAGACAATTATTTGATGAAGCTGTACAAATGATGGATCTACTTATTAATAAAATGGACTTAGCTTTAAATCCCACGAAATTA

>MSRP6Δ3

ATGAAAAGCAAAAAAATAATATGTTCATCTTGCTTATTTTTAATATTTTTAAGTGTAATATTTTGTAGTGAACCAGATACAAATTCATTTGATGAAAATGTAAAGAAGAATGAAGTTTTTAATGCCTTAAATGAACATTTAGAAAGTATAAGTAATATCGTAAAAGTAAATATTATGGATGCCCTTTCAAATAACCCTTCGTTAATAAAAAAGACATACGAAGCTGTTGAAATAAATGATGATGATTATGTTCTGGAATATGTTGGGGATTCGACAGGAAAATATGGAGAGGGCTCTATATTCTATGATGAAAGTAAGAAATATAATTATAGAAAGTATTTAGATATAGAAAATGAATTACGAAATATGAAAGGTGAAGATGATGATGATTTTGATGAAGATGATGATGATTTTGATGAAGATGATGAAGATGATGAAGATGATGAAGATGATGAAGATGATGTAGATGATGAAGATGATCTAGATGTTGAAGATAATGTAGATGATGAATATGATGATGAGCATAATCATAATTATAATGATGACAAATTGAGTGAAAATCCTGAGAAGTATTCAAATTATAATAAAAATATACACGAAGATAAGAAAAAAGATAATTTGAATGAACCACATTTTAAACAAACTCATTATATATATTCATCAAATCACGATAATAATGAAACATCTAGATTTCCAAAAAAAAATGTACCTAAATATGATGAAAAATTAAATAATGAATTTAAAACATATTTAAGAAGACCTGAAAAGAAAGAAGAAACAAAAGAATATCCTAACAATGGATGTTCTGTAGTTCAAATATCTATAGTAACTAATGAAGATTTTTTAAAAAAAGTTAGAGAAAGAAATAAAAAGAGAAATAATAAAAAAAGAACAAATATTTATGATAGTGATGAAGAATCAGAAAGTTCAGAAGAAACAAGTAAAGATCCCTATTCATCGGGTCCATAT

>ETRAMP10.1SP-MSRP6

ATGAAAAGCAAAAAAATAATATGTTCATCTTGCTTATTTTTAATATTTTTAAGTGTAATATTTTGTAGTGAACCAGATACAAATTCATTTGATGAAAATGTAAAGAAGAATGAAGTTTTTAATGCCTTAAATGAACATTTAGAAAGTATAAGTAATATCGTAAAAGTAAATATTATGGATGCCCTTTCAAATAACCCTTCGTTAATAAAAAAGACATACGAAGCTGTTGAAATAAATGATGATGATTATGTTCTGGAATATGTTGGGGATTCGACAGGAAAATATGGAGAGGGCTCTATATTCTATGATGAAAGTAAGAAATATAATTATAGAAAGTATTTAGATATAGAAAATGAATTACGAAATATGAAAGGTGAAGATGATGATGATTTTGATGAAGATGATGATGATTTTGATGAAGATGATGAAGATGATGAAGATGATGAAGATGATGAAGATGATGTAGATGATGAAGATGATCTAGATGTTGAAGATAATGTAGATGATGAATATGATGATGAGCATAATCATAATTATAATGATGACAAATTGAGTGAAAATCCTGAGAAGTATTCAAATTATAATAAAAATATACACGAAGATAAGAAAAAAGATAATTTGAATGAACCACATTTTAAACAAACTCATTATATATATTCATCAAATCACGATAATAATGAAACATCTAGATTTCCAAAAAAAAATGTACCTAAATATGATGAAAAATTAAATAATGAATTTAAAACATATTTAAGAAGACCTGAAAAGAAAGAAGAAACAAAAGAATATCCTAACAATGGATGTTCTGTAGTTCAAATATCTATAGTAACTAATGAAGATTTTTTAAAAAAAGTTAGAGAAAGAAATAAAAAGAGAAATAATAAAAAAAGAACAAATATTTATGATAGTGATGAAGAATCAGAAAGTTCAGAAGAAACAAGTAAAGATCCCTATTCATCGGGTCCATATACAGTTGATCATAAAAATGAAAATTGTTTTCCTTACAAGCAAAATTCAGATATTCATCCAAATATAAATAATGATAATAATGATAATAATGAAAATAATGATAATAATGAAAATAATGATAATAATATACCTGTTAATAATAAAAAAAATTCTCATACTCCTCATGTACCAAAAGATACCAAAAATAATATAGAAAAAAATCATAGTAACTATAATTACAAACCAATTAATCAAGAAAATTTATCAATGTATTTTAATAATAATAATAAGAAGAAGAATAAAAAGAATAGTCAAAAAAATAATCAAAAGAATAATCAAAAGAATAATCAAAAGCATAACCAAGATCTTTATAATATAAAAAAAAATAATGATCTACCTCAATATGAAGAATCTAATAAAACTTATAAAAAAATTTTTTCAAATGAACATGAACCACGTCCGATGATAACAACATCCAGATATGAAAATAAAATATCAAATGATAATAACAACAATAACTATAATTTGAATGAACAAGAAAGAAAACTTATTAAAAATATGATAGATATATTTATTTATACATACAAGTTAAATTATACAAACAGTAAATCTATATCTAAATTATTTAAAAATAATTTACTTAAAAAAAACTTCCGTACTTATTTCACAAATTATATATATACACTTTTTAATTATGGAAAGACATATAATTTTCTAACACCTTATAATAAAGATAATGATCATATGTATAGACAATTATTTGATGAAGCTGTACAAATGATGGATCTACTTATTAATAAAATGGACTTAGCTTTAAATCCCACGAAATTA

>MSRP6trunc(Domain1+part of domain 2)

ATGAAAAGCAAAAAAATAATATGTTCATCTTGCTTATTTTTAATATTTTTAAGTGTAATATTTTGTAGTGAACCAGATACAAATTCATTTGATGAAAATGTAAAGAAGAATGAAGTTTTTAATGCCTTAAATGAACATTTAGAAAGTATAAGTAATATCGTAAAAGTAAATATTATGGATGCCCTTTCAAATAACCCTTCGTTAATAAAAAAGACATACGAAGCTGTTGAAATAAATGATGATGATTATGTTCTGGAATATGTTGGGGATTCGACAGGAAAATATGGAGAGGGCTCTATATTCTATGATGAAAGTAAGAAATATAATTATAGAAAGTATTTAGATATAGAAAATGAATTACGAAATATGAAAGGATCAGGAGGTTCA

>MSRP6trunc(Domain1)

ATGAAAAGCAAAAAAATAATATGTTCATCTTGCTTATTTTTAATATTTTTAAGTGTAATATTTTGTAGTGAACCAGATACAAATTCATTTGATGAAAATGTAAAGAAGAATGAAGTTTTTAATGCCTTAAATGAACATTTAGAAAGTATAAGTAATATC

>PFB0115wtrunc(aa1-76)

ATGTTAAAGAAATATATTATATTAATATATATCGGTGTAATTCTAAATTTCATAACTAAAAATAATAATGTAGTGTCTGTTCCTGAGCCCTTTTTATCACAAAACAAAGATTCTTTTGAAGAAAAAAAATATACGTATGGGGATAATTTACAATTGGGGGCATCAACTATAAATACCCCTAAAACACAATCACAAGAAAATAAAGATATAAATAAAGAAACAAAAAAT

>MSRP6Δ3ab

ATGAAAAGCAAAAAAATAATATGTTCATCTTGCTTATTTTTAATATTTTTAAGTGTAATATTTTGTAGTGAACCAGATACAAATTCATTTGATGAAAATGTAAAGAAGAATGAAGTTTTTAATGCCTTAAATGAACATTTAGAAAGTATAAGTAATATCGTAAAAGTAAATATTATGGATGCCCTTTCAAATAACCCTTCGTTAATAAAAAAGACATACGAAGCTGTTGAAATAAATGATGATGATTATGTTCTGGAATATGTTGGGGATTCGACAGGAAAATATGGAGAGGGCTCTATATTCTATGATGAAAGTAAGAAATATAATTATAGAAAGTATTTAGATATAGAAAATGAATTACGAAATATGAAAGGTGAAGATGATGATGATTTTGATGAAGATGATGATGATTTTGATGAAGATGATGAAGATGATGAAGATGATGAAGATGATGAAGATGATGTAGATGATGAAGATGATCTAGATGTTGAAGATAATGTAGATGATGAATATGATGATGAGCATAATCATAATTATAATGATGACAAATTGAGTGAAAATCCTGAGAAGTATTCAAATTATAATAAAAATATACACGAAGATAAGAAAAAAGATAATTTGAATGAACCACATTTTAAACAAACTCATTATATATATTCATCAAATCACGATAATAATGAAACATCTAGATTTCCAAAAAAAAATGTACCTAAATATGATGAAAAATTAAATAATGAATTTAAAACATATTTAAGAAGACCTGAAAAGAAAGAAGAAACAAAAGAATATCCTAACAATGGATGTTCTGTAGTTCAAATATCTATAGTAACTAATGAAGATTTTTTAAAAAAAGTTAGAGAAAGAAATAAAAAGAGAAATAATAAAAAAAGAACAAATATTTATGATAGTGATGAAGAATCAGAAAGTTCAGAAGAAACAAGTAAAGATCCCTATTCATCGGGTCCATATCTACCTCAATATGAAGAATCTAATAAAACTTATAAAAAAATTTTTTCAAATGAACATGAACCACGTCCGATGATAACAACATCCAGATATGAAAATAAAATATCAAATGATAATAACAACAATAACTATAATTTGAATGAACAAGAAAGAAAACTTATTAAAAATATGATAGATATATTTATTTATACATACAAGTTAAATTATACAAACAGTAAATCTATATCTAAATTATTTAAAAATAATTTACTTAAAAAAAACTTCCGTACTTATTTCACAAATTATATATATACACTTTTTAATTATGGAAAGACATATAATTTTCTAACACCTTATAATAAAGATAATGATCATATGTATAGACAATTATTTGATGAAGCTGTACAAATGATGGATCTACTTATTAATAAAATGGACTTAGCTTTAAATCCCACGAAATTA

>MSRP6Δ3bc

ATGAAAAGCAAAAAAATAATATGTTCATCTTGCTTATTTTTAATATTTTTAAGTGTAATATTTTGTAGTGAACCAGATACAAATTCATTTGATGAAAATGTAAAGAAGAATGAAGTTTTTAATGCCTTAAATGAACATTTAGAAAGTATAAGTAATATCGTAAAAGTAAATATTATGGATGCCCTTTCAAATAACCCTTCGTTAATAAAAAAGACATACGAAGCTGTTGAAATAAATGATGATGATTATGTTCTGGAATATGTTGGGGATTCGACAGGAAAATATGGAGAGGGCTCTATATTCTATGATGAAAGTAAGAAATATAATTATAGAAAGTATTTAGATATAGAAAATGAATTACGAAATATGAAAGGTGAAGATGATGATGATTTTGATGAAGATGATGATGATTTTGATGAAGATGATGAAGATGATGAAGATGATGAAGATGATGAAGATGATGTAGATGATGAAGATGATCTAGATGTTGAAGATAATGTAGATGATGAATATGATGATGAGCATAATCATAATTATAATGATGACAAATTGAGTGAAAATCCTGAGAAGTATTCAAATTATAATAAAAATATACACGAAGATAAGAAAAAAGATAATTTGAATGAACCACATTTTAAACAAACTCATTATATATATTCATCAAATCACGATAATAATGAAACATCTAGATTTCCAAAAAAAAATGTACCTAAATATGATGAAAAATTAAATAATGAATTTAAAACATATTTAAGAAGACCTGAAAAGAAAGAAGAAACAAAAGAATATCCTAACAATGGATGTTCTGTAGTTCAAATATCTATAGTAACTAATGAAGATTTTTTAAAAAAAGTTAGAGAAAGAAATAAAAAGAGAAATAATAAAAAAAGAACAAATATTTATGATAGTGATGAAGAATCAGAAAGTTCAGAAGAAACAAGTAAAGATCCCTATTCATCGGGTCCATATACAGTTGATCATAAAAATGAAAATTGTTTTCCTTACAAGCAAAATTCAGATATTCATCCAAATATAAATAATGATAATAATGATAATAATGAAAATAATGATAATAATGAAAATAATGATAATAATATACCTGTTAATAATAAAAAAAATTCTCATACTCCTCATGTACCAAAAGATACCAAAAATAATATAGAAAAAAATACAAACAGTAAATCTATATCTAAATTATTTAAAAATAATTTACTTAAAAAAAACTTCCGTACTTATTTCACAAATTATATATATACACTTTTTAATTATGGAAAGACATATAATTTTCTAACACCTTATAATAAAGATAATGATCATATGTATAGACAATTATTTGATGAAGCTGTACAAATGATGGATCTACTTATTAATAAAATGGACTTAGCTTTAAATCCCACGAAATTA

>MSRP6Δ3cd

ATGAAAAGCAAAAAAATAATATGTTCATCTTGCTTATTTTTAATATTTTTAAGTGTAATATTTTGTAGTGAACCAGATACAAATTCATTTGATGAAAATGTAAAGAAGAATGAAGTTTTTAATGCCTTAAATGAACATTTAGAAAGTATAAGTAATATCGTAAAAGTAAATATTATGGATGCCCTTTCAAATAACCCTTCGTTAATAAAAAAGACATACGAAGCTGTTGAAATAAATGATGATGATTATGTTCTGGAATATGTTGGGGATTCGACAGGAAAATATGGAGAGGGCTCTATATTCTATGATGAAAGTAAGAAATATAATTATAGAAAGTATTTAGATATAGAAAATGAATTACGAAATATGAAAGGTGAAGATGATGATGATTTTGATGAAGATGATGATGATTTTGATGAAGATGATGAAGATGATGAAGATGATGAAGATGATGAAGATGATGTAGATGATGAAGATGATCTAGATGTTGAAGATAATGTAGATGATGAATATGATGATGAGCATAATCATAATTATAATGATGACAAATTGAGTGAAAATCCTGAGAAGTATTCAAATTATAATAAAAATATACACGAAGATAAGAAAAAAGATAATTTGAATGAACCACATTTTAAACAAACTCATTATATATATTCATCAAATCACGATAATAATGAAACATCTAGATTTCCAAAAAAAAATGTACCTAAATATGATGAAAAATTAAATAATGAATTTAAAACATATTTAAGAAGACCTGAAAAGAAAGAAGAAACAAAAGAATATCCTAACAATGGATGTTCTGTAGTTCAAATATCTATAGTAACTAATGAAGATTTTTTAAAAAAAGTTAGAGAAAGAAATAAAAAGAGAAATAATAAAAAAAGAACAAATATTTATGATAGTGATGAAGAATCAGAAAGTTCAGAAGAAACAAGTAAAGATCCCTATTCATCGGGTCCATATACAGTTGATCATAAAAATGAAAATTGTTTTCCTTACAAGCAAAATTCAGATATTCATCCAAATATAAATAATGATAATAATGATAATAATGAAAATAATGATAATAATGAAAATAATGATAATAATATACCTGTTAATAATAAAAAAAATTCTCATACTCCTCATGTACCAAAAGATACCAAAAATAATATAGAAAAAAATCATAGTAACTATAATTACAAACCAATTAATCAAGAAAATTTATCAATGTATTTTAATAATAATAATAAGAAGAAGAATAAAAAGAATAGTCAAAAAAATAATCAAAAGAATAATCAAAAGAATAATCAAAAGCATAACCAAGATCTTTATAATATAAAAAAAAATAATGAT

>Rex3-3cd

ATGCAAACCCGTAAATATAATAAGATGTTGTCAAAAGTTGAAACGAAACAATTTATATACATTCTTTTTTTCCTTTGCTTATATCTTAATACGTTCAACTATAAATATACCACCTCATACGAAGGGAGCAGTTTCAGGCAATTATCTGAACCAGTTGTAGAAGAACAGGATTTAAAAAAAACAAATGCAGAGTCATCACATATAGAAGCAACAACTTCACAAGCAACAACTTCACAAGCAACAACTTCACAAGCAACAACTTCACAAGCAACAACTTCACAAGCAACAACTTCACAAGCAACATCTTCACAAGAATCAGATGAACAAGGACTAACGGCACCATCATTAAATTTAGAAGAAACCCAATCAAACAAAGTACGTAATAAAATTTTTAACTTCCCTATACCATCCGCAGAAGGTACAGTATCAAAAGAATTTAAAAACCAACCAAAAACTGAATATGAAAAAAAATTATTTGAAGAATGGCAACACTTAAATATGTTTGAACATTCTAATTGGGTAAATATAACTGTACAAAGCTGTCAGGTTTTAGTACAAGGATTAACATCATTAGATGATTATGATGCTAAATTTAAAAGTTGGTCTGCTATGGTAGAACTATTAGGAGAATTCCGTATTACTTTATTTAACGAAAGTAATAATATGTTCGAAGCACTATTAAATGAATTAAGAGAAGCAAGAAAAGAAAATCCAAATGAAAACTTAACACCAGAAGAAGAAGAAAAATGGGATCTAATAAAACAAACTAAGCTAGAAAAAGATATTGAATGGAAAATTTATCAAATATTAACATGGAAATATTGGAATCTAAAAGAATTCCCAGGTGTTGATATACCTGATCCTAGTGTACCCTCTCTAGATTTTGATGCCACATACGATGTTCTTGGGGATATTCTTGAAGATGACGAAGATGAAGAAGATGATGAAGATACGGCTAAACCAAGTACAAGTTCTTCACTACCTCAATATGAAGAATCTAATAAAACTTATAAAAAAATTTTTTCAAATGAACATGAACCACGTCCGATGATAACAACATCCAGATATGAAAATAAAATATCAAATGATAATAACAACAATAACTATAATTTGAATGAACAAGAAAGAAAACTTATTAAAAATATGATAGATATATTTATTTATACATACAAGTTAAATTATACAAACAGTAAATCTATATCTAAATTATTTAAAAATAATTTACTTAAAAAAAACTTCCGTACTTATTTCACAAATTATATATATACACTTTTTAATTATGGAAAGACATATAATTTTCTAACACCTTATAATAAAGATAATGATCATATGTATAGACAATTATTTGATGAAGCTGTACAAATGATGGATCTACTTATTAATAAAATGGACTTAGCTTTAAATCCCACGAAATTA

>Rex3trunc-3c

ATGCAAACCCGTAAATATAATAAGATGTTGTCAAAAGTTGAAACGAAACAATTTATATACATTCTTTTTTTCCTTTGCTTATATCTTAATACGTTCAACTATAAATATACCACCTCATACGAAGGGAGCAGTTTCAGGCAATTATCTGAACCAGTTGTAGAAGAACAGGATTTAAAAAAAACAAATGCAGAGTCATCACATATAGAAGCAACAACTTCACAAGCAACAACTTCACAAGCAACAACTTCACAAGCAACAACTTCACAAGCAACAACTTCACAAGCAACAACTTCACAAGCAACATCTTCACAAGAATCAGATGAACAAGGACTAACGGCACCATCATTAAATTTAGAAGAAACCCAATCAAACAAAGTACGTAATAAAATTTTTAACTTCCCTATACCATCCGCAGAAGGTACAGTATCAAAAGAATTTAAAAACCAACCAAAAACTGAATATGAAAAAAAATTATTTGAAGAATGGCAACACTTAAATATGTTTGAACATTCTAATTGGGTAAATATAACTGTACAAAGCTGTCAGGTTTTAGTACAAGGATTAACATCATTAGATGATTATGATGCTAAATTTAAAAGTTGGTCTGCTATGGTAGAACTATTAGGAGAATTCCGTATTACTTTATTTAACGAAAGTAATAATATGTTCGAAGCACTATTAAATGAATTAAGAGAAGCAAGAAAAGAAAATCCAAATGAAAACTTAACACCAGAAGAAGAAGAAAAATGGGATCTAATAAAACAAACTAAGCTAGAAAAAGATATTGAATGGAAAATTTATCAAATATTAACATGGAAATATTGGAATCTAAAAGAATTCCCAGGTGTTGATATACCTGATCCTAGTGTACCCTCTCTAGATTTTGATGCCACATACGATGTTCTTGGGGATATTCTTGAAGATGACGAAGATGAAGAAGATGATGAAGATACGGCTAAACCAAGTACAAGTTCTTCACTACCTCAATATGAAGAATCTAATAAAACTTATAAAAAAATTTTTTCAAATGAACATGAACCACGTCCGATGATAACAACATCCAGATATGAAAATAAAATATCAAATGATAATAACAACAATAACTATAATTTGAATGAACAAGAAAGAAAACTTATTAAAAATATGATAGATATATTTATTTATACATACAAGTTAAATTAT

>Rex3trunc-3d

ATGCAAACCCGTAAATATAATAAGATGTTGTCAAAAGTTGAAACGAAACAATTTATATACATTCTTTTTTTCCTTTGCTTATATCTTAATACGTTCAACTATAAATATACCACCTCATACGAAGGGAGCAGTTTCAGGCAATTATCTGAACCAGTTGTAGAAGAACAGGATTTAAAAAAAACAAATGCAGAGTCATCACATATAGAAGCAACAACTTCACAAGCAACAACTTCACAAGCAACAACTTCACAAGCAACAACTTCACAAGCAACAACTTCACAAGCAACAACTTCACAAGCAACATCTTCACAAGAATCAGATGAACAAGGACTAACGGCACCATCATTAAATTTAGAAGAAACCCAATCAAACAAAGTACGTAATAAAATTTTTAACTTCCCTATACCATCCGCAGAAGGTACAGTATCAAAAGAATTTAAAAACCAACCAAAAACTGAATATGAAAAAAAATTATTTGAAGAATGGCAACACTTAAATATGTTTGAACATTCTAATTGGGTAAATATAACTGTACAAAGCTGTCAGGTTTTAGTACAAGGATTAACATCATTAGATGATTATGATGCTAAATTTAAAAGTTGGTCTGCTATGGTAGAACTATTAGGAGAATTCCGTATTACTTTATTTAACGAAAGTAATAATATGTTCGAAGCACTATTAAATGAATTAAGAGAAGCAAGAAAAGAAAATCCAAATGAAAACTTAACACCAGAAGAAGAAGAAAAATGGGATCTAATAAAACAAACTAAGCTAGAAAAAGATATTGAATGGAAAATTTATCAAATATTAACATGGAAATATTGGAATCTAAAAGAATTCCCAGGTGTTGATATACCTGATCCTAGTGTACCCTCTCTAGATTTTGATGCCACATACGATGTTCTTGGGGATATTCTTGAAGATGACGAAGATGAAGAAGATGATGAAGATACGGCTAAACCAAGTACAAGTTCTTCAACAAACAGTAAATCTATATCTAAATTATTTAAAAATAATTTACTTAAAAAAAACTTCCGTACTTATTTCACAAATTATATATATACACTTTTTAATTATGGAAAGACATATAATTTTCTAACACCTTATAATAAAGATAATGATCATATGTATAGACAATTATTTGATGAAGCTGTACAAATGATGGATCTACTTATTAATAAAATGGACTTAGCTTTAAATCCCACGAAATTA

>MSRP6-SP-3cd

ATGAAAAGCAAAAAAATAATATGTTCATCTTGCTTATTTTTAATATTTTTAAGTGTAATATTTTGTCTACCTCAATATGAAGAATCTAATAAAACTTATAAAAAAATTTTTTCAAATGAACATGAACCACGTCCGATGATAACAACATCCAGATATGAAAATAAAATATCAAATGATAATAACAACAATAACTATAATTTGAATGAACAAGAAAGAAAACTTATTAAAAATATGATAGATATATTTATTTATACATACAAGTTAAATTATACAAACAGTAAATCTATATCTAAATTATTTAAAAATAATTTACTTAAAAAAAACTTCCGTACTTATTTCACAAATTATATATATACACTTTTTAATTATGGAAAGACATATAATTTTCTAACACCTTATAATAAAGATAATGATCATATGTATAGACAATTATTTGATGAAGCTGTACAAATGATGGATCTACTTATTAATAAAATGGACTTAGCTTTAAATCCCACGAAATTAGGATCAGGAGGTTCA

>Rex3trunc-BirA*

ATGCAAACCCGTAAATATAATAAGATGTTGTCAAAAGTTGAAACGAAACAATTTATATACATTCTTTTTTTCCTTTGCTTATATCTTAATACGTTCAACTATAAATATACCACCTCATACGAAGGGAGCAGTTTCAGGCAATTATCTGAACCAGTTGTAGAAGAACAGGATTTAAAAAAAACAAATGCAGAGTCATCACATATAGAAGCACTGCAGGGATCAGGAGGTTCATCCGGTGGATCTGGATCCTCTATGAAAGATAATACAGTACCATTAAAATTAATAGCTTTATTAGCTAATGGTGAATTTCATTCAGGTGAACAATTAGGTGAAACATTAGGTATGTCAAGAGCTGCTATAAATAAACATATACAAACATTAAGAGATTGGGGTGTAGATGTATTTACAGTACCAGGTAAAGGTTATTCATTACCAGAACCAATACAATTATTAAATGCTGAAAAAATATTATCACAATTAGATGATGGTTCAGTAGCTGTATTACCAGTAATAGATTCAACAAATCAATATTTATTAGATAGAATAGGTGAATTAAAATCAGGTGATGCTTGTGTAGCTGAATATCAACATGCTGGTAGAGGTGGTAGAGGTAGAAAATGGTTTTCACCATTTGGTGCTAATTTATATTTATCAATGTTTTGGAGATTAGAACAAGGTCCAGCTGCTGCTATAGGTTTATCATTAGTAATAGGTATAGTAATGGCTGAAGTATTAAGAAAATTAGGTGCTGATAAAGTAAGAGTAAAATGGCCAAATGATTTATATTTACAAGATAGAAAATTAGCTGGTATATTAGTAGAATTAACAGGTAAAACAGGTGATGCTGCTCAAATAGTAATAGGTGCTGGTATAAATATGGCTATGAGAAGAGTAGAAGAATCAGTAGTAAATCAAGGTTGGATAACATTACAAGAAGCTGGTATAAATTTAGATAGAAATACATTAGCTGCTATGTTAATAAGAGAATTAAGAGCTGCTTTAGAATTATTTGAACAAGAAGGTTTAGCTCCATATTTATCAAGATGGGAAAAATTAGATAATTTTATAAATAGACCAGTAAAATTAATAATAGGTGATAAAGAAATATTTGGTATATCAAGAGGTATAGATAAACAAGGTGCTTTATTATTAGAACAAGATGGTATAATAAAACCATGGATGGGTGGTGAAATATCATTAAGATCAGCTGAAAAAGGTTCAGGATCTGGTAGTGGATCAGGTTCTGGAAGTGGTTCTGGAAGT

>Rec3trunc-MAD-BirA*

ATGCAAACCCGTAAATATAATAAGATGTTGTCAAAAGTTGAAACGAAACAATTTATATACATTCTTTTTTTCCTTTGCTTATATCTTAATACGTTCAACTATAAATATACCACCTCATACGAAGGGAGCAGTTTCAGGCAATTATCTGAACCAGTTGTAGAAGAACAGGATTTAAAAAAAACAAATGCAGAGTCATCACATATAGAAGCACTGCAGCTACCTCAATATGAAGAATCTAATAAAACTTATAAAAAAATTTTTTCAAATGAACATGAACCACGTCCGATGATAACAACATCCAGATATGAAAATAAAATATCAAATGATAATAACAACAATAACTATAATTTGAATGAACAAGAAAGAAAACTTATTAAAAATATGATAGATATATTTATTTATACATACAAGTTAAATTATACAAACAGTAAATCTATATCTAAATTATTTAAAAATAATTTACTTAAAAAAAACTTCCGTACTTATTTCACAAATTATATATATACACTTTTTAATTATGGAAAGACATATAATTTTCTAACACCTTATAATAAAGATAATGATCATATGTATAGACAATTATTTGATGAAGCTGTACAAATGATGGATCTACTTATTAATAAAATGGACTTAGCTTTAAATCCCACGAAATTACTGCAGGGATCAGGAGGTTCATCCGGTGGATCTGGATCCTCTATGAAAGATAATACAGTACCATTAAAATTAATAGCTTTATTAGCTAATGGTGAATTTCATTCAGGTGAACAATTAGGTGAAACATTAGGTATGTCAAGAGCTGCTATAAATAAACATATACAAACATTAAGAGATTGGGGTGTAGATGTATTTACAGTACCAGGTAAAGGTTATTCATTACCAGAACCAATACAATTATTAAATGCTGAAAAAATATTATCACAATTAGATGATGGTTCAGTAGCTGTATTACCAGTAATAGATTCAACAAATCAATATTTATTAGATAGAATAGGTGAATTAAAATCAGGTGATGCTTGTGTAGCTGAATATCAACATGCTGGTAGAGGTGGTAGAGGTAGAAAATGGTTTTCACCATTTGGTGCTAATTTATATTTATCAATGTTTTGGAGATTAGAACAAGGTCCAGCTGCTGCTATAGGTTTATCATTAGTAATAGGTATAGTAATGGCTGAAGTATTAAGAAAATTAGGTGCTGATAAAGTAAGAGTAAAATGGCCAAATGATTTATATTTACAAGATAGAAAATTAGCTGGTATATTAGTAGAATTAACAGGTAAAACAGGTGATGCTGCTCAAATAGTAATAGGTGCTGGTATAAATATGGCTATGAGAAGAGTAGAAGAATCAGTAGTAAATCAAGGTTGGATAACATTACAAGAAGCTGGTATAAATTTAGATAGAAATACATTAGCTGCTATGTTAATAAGAGAATTAAGAGCTGCTTTAGAATTATTTGAACAAGAAGGTTTAGCTCCATATTTATCAAGATGGGAAAAATTAGATAATTTTATAAATAGACCAGTAAAATTAATAATAGGTGATAAAGAAATATTTGGTATATCAAGAGGTATAGATAAACAAGGTGCTTTATTATTAGAACAAGATGGTATAATAAAACCATGGATGGGTGGTGAAATATCATTAAGATCAGCTGAAAAAGGTTCAGGATCTGGTAGTGGATCAGGTTCTGGAAGTGGTTCTGGAAGT>

>STEVOR1-260-GFB-BirA* (cloned between KpnI and XhoI)

ATGAAGATGTATTACCTTAAAATGTTATTGTTTAACTTTTTAATAAATGTTTTAGTATTACCACATTATGAAAATTATCAAAATAACCATTATAACATAAGGCTCATACCAAATAACACATACAGAATAACGATAAAATCAAGACTCTTAGCACAAACCCAAATCCATAATCCACATTATCATAATGATCCAGAACTCAAAGAAATAATTGATAAATTGAACGAAGACGCAATAAAGAAATACCAACAAACTCATGATCCATATGAACAATTACAAGAAGTAGTAGAAAAAAATGGAACAAAATATAAAGGTGGTAATGATGCAGAACCTATGTCAACGCTAGAAAAAGAATTATTGCAAACATATGAAGAAGTGTTTGGTAACGAAAGTGATATGTTGAAGTCGGGAATGAATACAAATGTTGATGAAAAATCTTCAACATGTGGATGTACTGATATTAATGGTGTGAAATTAGCAAAAACAAAAGGAAGAGATAAGTATTTAAAACACTTAAAACATAGATGTACCCGTGGTATATGTTTTTGCTCAGTTGGTAGTGCATTGTTAACTATGTTCGGTTTGGCAGTTGCAAAAAAAGCTGCCGTTGATGCCATCCTTCCTGTATATGTAGCAGCGATTAAAAAGTGTGTATCCTCCAGTTCATTATTTCATATATTTCATGGTGGTTCATTAACTACAGCTCTTAAAGCAACCGAAGCGTGTGCAAGTGTTGCAGGTCCAGATATAGTCATACCTGCTACAGGTGCTGCTATAGGTGCACCTAGGATGAGTAAAGGAGAAGAACTTTTCACTGGAGTTGTCCCAATTCTTGTTGAATTAGATGGTGATGTTAATGGGCACAAATTTTCTGTCAGTGGAGAGGGTGAAGGTGATGCAACATACGGAAAACTTACCCTTAAATTTATTTGCACTACTGGAAAACTACCAGTTCCATGGCCAACACTTGTCACTACTTTCGCGTATGGTCTTCAATGCTTTGCGAGATACCCAGATCATATGAAACAGCATGACTTTTTCAAGAGTGCCATGCCCGAAGGTTATGTACAGGAAAGAACTATATTTTTCAAAGATGACGGGAACTACAAGACACGTGCTGAAGTCAAGTTTGAAGGTGATACCCTTGTTAATAGAATCGAGTTAAAAGGTATTGATTTTAAAGAAGATGGAAACATTCTTGGACACAAATTGGAATACAACTATAACTCACACAATGTATACATCATGGCAGACAAACAAAAGAATGGAATCAAAGTTAACTTCAAAATTAGACACAACATTGAAGATGGAAGCGTTCAACTAGCAGACCATTATCAACAAAATACTCCAATTGGCGATGGCCCTGTCCTTTTACCAGACAACCATTACCTGTCCACACAATCTGCCCTTTCGAAAGATCCCAACGAAAAGAGAGACCACATGGTCCTTCTTGAGTTTGTAACAGCTGCTGGGATTACACATGGCATGGATGAGCTCTACAAAGCTAGCTCAGGATCTGGTAGTGGATCAGGTTCTGGAAGTGGTTCTGGAAGTATGAAAGATAATACAGTACCATTAAAATTAATAGCTTTATTAGCTAATGGTGAATTTCATTCAGGTGAACAATTAGGTGAAACATTAGGTATGTCAAGAGCTGCTATAAATAAACATATACAAACATTAAGAGATTGGGGTGTAGATGTATTTACAGTACCAGGTAAAGGTTATTCATTACCAGAACCAATACAATTATTAAATGCTGAAAAAATATTATCACAATTAGATGATGGTTCAGTAGCTGTATTACCAGTAATAGATTCAACAAATCAATATTTATTAGATAGAATAGGTGAATTAAAATCAGGTGATGCTTGTGTAGCTGAATATCAACATGCTGGTAGAGGTGGTAGAGGTAGAAAATGGTTTTCACCATTTGGTGCTAATTTATATTTATCAATGTTTTGGAGATTAGAACAAGGTCCAGCTGCTGCTATAGGTTTATCATTAGTAATAGGTATAGTAATGGCTGAAGTATTAAGAAAATTAGGTGCTGATAAAGTAAGAGTAAAATGGCCAAATGATTTATATTTACAAGATAGAAAATTAGCTGGTATATTAGTAGAATTAACAGGTAAAACAGGTGATGCTGCTCAAATAGTAATAGGTGCTGGTATAAATATGGCTATGAGAAGAGTAGAAGAATCAGTAGTAAATCAAGGTTGGATAACATTACAAGAAGCTGGTATAAATTTAGATAGAAATACATTAGCTGCTATGTTAATAAGAGAATTAAGAGCTGCTTTAGAATTATTTGAACAAGAAGGTTTAGCTCCATATTTATCAAGATGGGAAAAATTAGATAATTTTATAAATAGACCAGTAAAATTAATAATAGGTGATAAAGAAATATTTGGTATATCAAGAGGTATAGATAAACAAGGTGCTTTATTATTAGAACAAGATGGTATAATAAAACCATGGATGGGTGGTGAAATATCATTAAGATCAGCTGAAAAAGGTTAA

SLI integration constructs (2xFKBP-GFP)

2xFKBP

GFP

T2A

Neomycin-R

>pArl1-**Insert**-2xFKBP-GFP-T2A-NeoR

gcggccgc**Insert**cctaggTCAGGATTGAGATCAAGATCTGCTGCTGCTGGTGCTGGTGGTGCTGCTAGAGCTGCTctgcagAGAGGAGTACAAGTTGAAACAATATCACCAGGAGATGGTCGTACATTTCCAAAAAGAGGTCAAACTTGTGTTGTACATTATACTGGAATGCTTGAAGATGGAAAGAAATTTGATTCATCTCGTGATAGAAATAAACCATTTAAATTTATGCTAGGTAAACAAGAAGTAATACGAGGTTGGGAAGAAGGAGTTGCTCAAATGAGTGTAGGTCAAAGAGCAAAACTTACTATATCTCCAGATTATGCTTATGGTGCAACTGGACATCCAGGTATAATTCCACCTCATGCAACTCTTGTATTTGATGTGGAGCTTCTAAAACTAGAAACTAGAGGTGTTCAGGTTGAAACAATTTCACCTGGAGATGGCAGAACCTTTCCTAAAAGAGGACAGACTTGCGTAGTTCATTATACAGGCATGCTAGAGGATGGTAAGAAATTTGATTCTAGTCGAGATAGAAATAAGCCATTCAAGTTTATGCTAGGTAAACAGGAAGTAATAAGAGGTTGGGAAGAGGGTGTAGCACAGATGTCAGTTGGACAAAGAGCAAAGTTAACAATATCACCAGATTATGCATACGGTGCAACAGGCCATCCTGGCATCATCCCTCCACATGCAACTTTAGTATTCGACGTTGAATTGTTAAAGTTAGAGACAacgcgtGCTAGAGGTGCTGCTGCTGGTGCTGGAGGTGCAGGTAGACGTACGATGAGTAAAGGAGAAGAACTTTTCACTGGAGTTGTCCCAATTCTTGTTGAATTAGATGGTGATGTTAATGGGCACAAATTTTCTGTCAGTGGAGAGGGTGAAGGTGATGCAACATACGGAAAACTTACCCTTAAATTTATTTGCACTACTGGAAAACTACCTGTTCCATGGCCAACACTTGTCACTACTTTCGCGTATGGTCTTCAATGCTTTGCGAGATACCCAGATCATATGAAACAGCATGACTTTTTCAAGAGTGCCATGCCCGAAGGTTATGTACAGGAAAGAACTATATTTTTCAAAGATGACGGGAACTACAAGACACGTGCTGAAGTCAAGTTTGAAGGTGATACCCTTGTTAATAGAATCGAGTTAAAAGGTATTGATTTTAAAGAAGATGGAAACATTCTTGGACACAAATTGGAATACAACTATAACTCACACAATGTATACATCATGGCAGACAAACAAAAGAATGGAATCAAAGTTAACTTCAAAATTAGACACAACATTGAAGATGGAAGCGTTCAACTAGCAGACCATTATCAACAAAATACTCCAATTGGCGATGGCCCTGTCCTTTTACCAGACAACCATTACCTGTCCACACAATCTGCCCTTTCGAAAGATCCCAACGAAAAGAGAGACCACATGGTCCTTCTTGAGTTTGTAACAGCTGCTGGGATTACACATGGCATGGATGAGCTCTACAAAGTCGACGGAGAAGGAAGAGGAAGTTTATTAACATGTGGAGATGTAGAAGAAAATCCAGGACCAATGATTGAACAAGATGGATTGCACGCAGGTTCTCCGGCCGCTTGGGTGGAGAGGCTATTCGGCTATGACTGGGCACAACAGACAATCGGCTGCTCTGATGCCGCCGTGTTCCGGCTGTCAGCGCAGGGGCGCCCGGTTCTTTTTGTCAAGACCGACCTGTCCGGTGCCCTGAATGAACTGCAGGACGAGGCAGCGCGGCTATCGTGGCTGGCCACGACGGGCGTTCCTTGCGCAGCTGTGCTCGACGTTGTCACTGAAGCGGGAAGGGACTGGCTGCTATTGGGCGAAGTGCCGGGGCAGGATCTCCTGTCATCTCACCTTGCTCCTGCCGAGAAAGTATCCATCATGGCTGATGCAATGCGGCGGCTGCATACGCTTGATCCGGCTACCTGCCCATTCGACCACCAAGCGAAACATCGCATCGAGCGAGCACGTACTCGGATGGAAGCCGGTCTTGTCGATCAGGATGATCTGGACGAAGAGCATCAGGGGCTCGCGCCAGCCGAACTGTTCGCCAGGCTCAAGGCGCGCATGCCCGACGGCGAGGATCTCGTCGTGACCCATGGCGATGCCTGCTTGCCGAATATCATGGTGGAAAATGGCCGCTTTTCTGGATTCATCGACTGTGGCCGGCTGGGTGTGGCGGACCGCTATCAGGACATAGCGTTGGCTACCCGTGATATTGCTGAAGAGCTTGGCGGCGAATGGGCTGACCGCTTCCTCGTGCTTTACGGTATCGCCGCTCCCGATTCGCAGCGCATCGCCTTCTATCGCCTTCTTGACGAGTTCTTCTAACTCGAGggatatggcagcttaatgttcgtttttcttatttatatatttataccaattgattgtatttataactgtaaaaatgtgtatgttgtgtgcatatttttttttgtgcatgcacatgcatgtaaatagctaaaattatgaacattttattttttgttcagaaaaaaaaaactttacacacataaaatggctagtatgaatagccatattttatataaattaaatcctatgaatttatgaccatattaaaaatttagatatttatggaacataatatgtttgaaacaataagacaaaattattattattattattatttttactgttataattatgtgtctccttcaatgattcataaatagttggacttgatttttaaaatgtttataatatgattagcatagttaaataaaaaaagttgaaaaattaaaaaaaaacatataaacacaaatgatggtttttccttcaatttcgatatcaatttatagaaacaaaatatatacttgtataattttatttttttatataaatcattacatatataattatacaatattttttctaagagataattatatattaatatatataaaaaaaggtgttttttttttttttttttatttttatttttattttatggtaatattttattttccttattttataaattatattagtttatatgtgattaattttatatattatcaatttatatatttttaaatgcttacttaattatctttttttttttttttttttttttttcccctctttttatattaatttatttttgaaaaaattgatatatatatatatatataatatatatatatacatgtagtagtattaaacaatgtataatatatataaataatatatttatatatttcatttcaattttaattttttttggttttttttttttttctttttgtcatatttaaaaaaaattatattcatataagttatgcattttttataaacattattcaatatatgtataatataatatatatatatatattaatgtattattccaatgtgcatgataaaagaaaaaaataatatttataaaaaaaaagaaaaataaaacaaaaaaagaaaaaaaaaaaaaaaaaaaaaaaaatacaaaaataaataatataatttataattatatattcttgtcacaataaaaatatatatatatatatatatatttataatatgtatattttaaactagaaaaggaataactaatattttatttattatcattcaagatttatattttataataataaatacctaatagaaatatatcaggatccatgcatggttcgctaaactgcatcgtcgctgtgtcccagaacatgggcatcggcaagaacggggactacccctggccaccgctcaggaacgaatttagatatttccagagaatgaccacaacctcttcagtagaaggtaaacagaatctggtgattatgggtaagaagacctggttctccattcctgagaagaatcgacctttaaagggtagaattaatttagttctcagcagagaactcaaggaacctccacaaggagctcattttctttccagaagtctagatgatgccttaaaacttactgaacaaccagaattagcaaataaagtagacatggtctggatagttggtggcagttctgtttataaggaagccatgaatcacccaggccatcttaaactatttgtgacaaggatcatgcaagactttgaaagtgacacgttttttccagaaattgatttggagaaatataaacttctgccagaatacccaggtgttctctctgatgtccaggaggagaaaggcattaagtacaaatttgaagtatatgagaagaatgattaagcttatttaataatagattaaaaatattataaaaataaaaacataaacacagaaattacaaaaaaaatacatatgaattttttttttgtaatcttccttataaatatagaataatgaatcatataaaacatatcattattcatttatttacatttaaaattattgtttcagtatctttaatttattatgtatatataaaaataacttacaattttattaataaacaatatatgtttattaattcatgttttgtaatttatgggatagcgattttttttactgtctgtatttttcttttttaattatgttttaattgtattttatttttattattgttctttttatagtattattttaaaacaaaatgtattttctaagaacttataataataataaatataaattttaataaaaattatatttatcttttacaatatgaacataaagtacaacattaatatatagcttttaatatttttattcctaatcatgtaaatcttaaatttttctttttaaacatatgttaaatatttatttctcattatatataagaacatatttattaaatctagaattctatagtgagtcgtattacaattcactggccgtcgttttacaacgtcgtgactgggaaaaccctggcgttacccaacttaatcgccttgcagcacatccccctttcgccagctggcgtaatagcgaagaggcccgcaccgatcgcccttcccaacagttgcgcagcctgaatggcgaatggcgcctgatgcggtattttctccttacgcatctgtgcggtatttcacaccgcatatggtgcactctcagtacaatctgctctgatgccgcatagttaagccagccccgacacccgccaacacccgctgacgcgccctgacgggcttgtctgctcccggcatccgcttacagacaagctgtgaccgtctccgggagctgcatgtgtcagaggttttcaccgtcatcaccgaaacgcgcgagacgaaagggcctcgtgatacgcctatttttataggttaatgtcatgataataatggtttcttagacgtcaggtggcacttttcggggaaatgtgcgcggaacccctatttgtttatttttctaaatacattcaaatatgtatccgctcatgagacaataaccctgataaatgcttcaataatattgaaaaaggaagagtatgagtattcaacatttccgtgtcgcccttattcccttttttgcggcattttgccttcctgtttttgctcacccagaaacgctggtgaaagtaaaagatgctgaagatcagttgggtgcacgagtgggttacatcgaactggatctcaacagcggtaagatccttgagagttttcgccccgaagaacgttttccaatgatgagcacttttaaagttctgctatgtggcgcggtattatcccgtattgacgccgggcaagagcaactcggtcgccgcatacactattctcagaatgacttggttgagtactcaccagtcacagaaaagcatcttacggatggcatgacagtaagagaattatgcagtgctgccataaccatgagtgataacactgcggccaacttacttctgacaacgatcggaggaccgaaggagctaaccgcttttttgcacaacatgggggatcatgtaactcgccttgatcgttgggaaccggagctgaatgaagccataccaaacgacgagcgtgacaccacgatgcctgtagcaatgccaacaacgttgcgcaaactattaactggcgaactacttactctagcttcccggcaacaattaatagactggatggaggcggataaagttgcaggaccacttctgcgctcggcccttccggctggctggtttattgctgataaatctggagccggtgagcgtgggtctcgcggtatcattgcagcactggggccagatggtaagccctcccgtatcgtagttatctacacgacggggagtcaggcaactatggatgaacgaaatagacagatcgctgagataggtgcctcactgattaagcattggtaactgtcagaccaagtttactcatatatactttagattgatttaaaacttcatttttaatttaaaaggatctaggtgaagatcctttttgataatctcatgaccaaaatcccttaacgtgagttttcgttccactgagcgtcagaccccgtagaaaagatcaaaggatcttcttgagatcctttttttctgcgcgtaatctgctgcttgcaaacaaaaaaaccaccgctaccagcggtggtttgtttgccggatcaagagctaccaactctttttccgaaggtaactggcttcagcagagcgcagataccaaatactgtccttctagtgtagccgtagttaggccaccacttcaagaactctgtagcaccgcctacatacctcgctctgctaatcctgttaccagtggctgctgccagtggcgataagtcgtgtcttaccgggttggactcaagacgatagttaccggataaggcgcagcggtcgggctgaacggggggttcgtgcacacagcccagcttggagcgaacgacctacaccgaactgagatacctacagcgtgagctatgagaaagcgccacgcttcccgaagggagaaaggcggacaggtatccggtaagcggcagggtcggaacaggagagcgcacgagggagcttccagggggaaacgcctggtatctttatagtcctgtcgggtttcgccacctctgacttgagcgtcgatttttgtgatgctcgtcaggggggcggagcctatcgaaaaacgccagcaacgcggcctttttacggttcctggccttttgctggccttttgctcacatgttctttcctgcgttatcccctgattctgtggataaccgtattaccgcctttgagtgagctgataccgctcgccgcagccgaacgaccgagcgcagcgagtcagtgagcgaggaagcggaagagcgcccaatacgcaaaccgcctctccccgcgcgttggccgattcattaatgcagctggcacgacaggtttcccgactggaaagcgggcagtgagcgcaacgcaattaatgtgagttagctcactcattaggcaccccaggctttacactttatgcttccggctcgtatgttgtgtggaattgtgagcggataacaatttcacacaggaaacagctatgaccatgattacgccaagctatttaggtgacactatagaatactc

**Inserts used:**

>PIESP2

TAACAAATCAACATAACTATAAAGAAGGTCCCTCATATGAAGATAAAAAAAATATGTACAAAGAAATATTGAAAGGATATTATAATGTATTTTTTGAAAATTATGCAAACGACACAGAATCAAATGTACATAATAAACCTGAGGAAGTTCATAAACATGAGGAAATTCATAAACATAGGAAACTTCATAAACATGAAGAAGTTCATAAACCTGAGGAATTTCATAAACCTGAGGAATTTCATAAACATGAGAAAGTTCATAAACATGAAGAAGTTCATAAACCTGAGGAAGTTCATAAACATGAGGAAAATCATAAACATGAGGAAAATCATAAACCTCAAATGGTAGGTCAAGCACCTCCAGAAAAAGAGATACGCCAAGAATCAAGAACTCTAATACTTGGTTCATTTCCCCAAGCAGGTGAAATATTAAGAGAGGATTTATGGAACAAAGAGGATAACAAATTTAGTTACGCACTTGACCCTAATGATTATGCATCTATAGAAGATAAACTTTTAGGATCTATATTTGGATACTTTAAAAAAAATCATGACAATTTGGTTAAACATTTGTTACAACAAATTAATACTTACAAACATAAATATATGGAACTTAAAGAACAATATATTAATGAAGTTATGAAACTTAAAAAAATATATAACAAAAGCATCATGGTCATATTTATAGCATCTTGTATTTCAATATTAGGACCTGTAATGTTACACATGCATCAAAATAATCCAGAAGAATTTTTTGCGACCATATTAAGTTTTTCTATATCATTAGGTCTTCATAATTTATTACTAACT

>PeMP1

TAAGGAGTTCTTTCCAAGAGTTATGATATGAATAAGAAGAATTTGAATAATGTAGGATTCCGAAATAACAGAATTTTGTCTAGTAAGGAAAACCGTGAAGAACCAGGAACTAGTGGTACAACATCAGGTGCTAATGGTAACAAGGCTAATATTCCTGACCTTGATAATTTTATAAATCAGACAACATCTTCATGTGGTAATTTGGGTAATTTGTTAGCACCATTACTTAAGGGTATTAAGTCTAATTTATTTTCAAAATATGTTAATGAAGATGGTACAACTAAACCAAATGCTCCTACAGCGCAGAGTTTAACCTCTATAGTTAAAGGAGTTATGCATGATATTAAGCCACAGATGGAAGCATTAAAAGAAGGTATAAAGAAAAATATTGAAGAAGAAGGTGAAGAAAGTATTAAGAAAGATTGTGATGCTTTGAAAGATGGATTTAAGAATATTATGAAAAGTTTTGGAGGTGATTTTTCTAACACTGTACAAGGTGCAACAGGAATACCGGGTAAATTATTCAAAGGGTTTTTAAATGAAGATGAAAGTAAAACTACAGAAAATGCAGATATTAAGGATGAAATAGATATATTATTAGAAGATGACAACACTGATGATGATGATGATGATGATGGTGATGATGATGATGATAATAATGATGATGATGACGATGAAAAAGAAAAAGAAAAAGAAAAGGAAAATGAAAAGGGAAAGGGAAAGGGAAAAGGAAAGAAAAAGGAAAACAAAGAAACCTCAGATGACAATAATAAAAATGCTGGTACTTCTACATTGAGAGGAAAAAAAAAAAAATTT

>PF10_0024

TAAGAACCAATTTTCGGATTTGATGTCATGTATGATTTATTTAAAGATAATTTTTTAGAACAAATTGGTTATAATGATGCACAAAAAAATGAATTAAAAGTTTCTATGAAATCGTTTTTTCAAAGTTCAAATAGTCAAGAATATACCTATTCTAGACGAGAAGATAATATTCATAGTATTAGTAAATCAAATGTGACAAACGTGAAAAGTGTGAAATTTGAAGAACCACAAGATGAGAAAAATTTAATTACAATTGAAGAAATAAAAGCAGACAATAATGATTCAAAAGAAAAAAAAAATGAAATAACTAATATAAACGTGGATGAAAAAAAGGATTTGGAAAGTGTAATTTCCAGTGAAGAAGAAAAAACACCAACTAGTAATAATATAAGGAAAAAGAGAAAGACCAAAAGGTCTCATAAGAATAAAGACAGTAACACATCTTATGATAATATAGAATTGAATAAAATATTAAAAGATGATAATATAGAAGTAAATAAAACATTAAAAGAAGATAATATAGAAGTAAATAATACATTAGAAGAAGATAATATAGAAGTGAATAATACATTAAAAGAAGATAATATAGAAGTTGAAAAACTAATTGATGAAAACAGAGAGTTAATTAAAAAAGAATCTATATATAAAAATAAATTAAAAGAAGATAATATAGAAGTTGAAAAACTAATTGATGAAAACAGAGAGTTAATTAAAAAAGAATCTATATATAAAAATAAATTAAAAGAAGATAATATAGAAGTTGAAGATCTAATTGATGAAAACAGAAAGTTAATTAAAAAAGAATCTATATATAAAAATAAATTAGAAGAAGAACCTGTTGAAAAAGGAATAAAAAGGAAAAATTATCTTTTTCTTATACTTGAAAAGGCAAAAGAAATTCCCCCAGGTGTTATTGTAGGATTATTTTCTATTTTTTACGTTGCACTAGGAGAAAATAGATCATCTTTAATTTTATTAATTGTAATATTTATTTTAACAGTACATACATATTTGGAAGTAAAAAGAATATGTCGATTCTTAAATATAGAT

>PF8_0003

TAAGGTCAAACCTAGAAGATACAGATCAAATCAGAGTATTAAATCACAAAGATAGAAAAGAATGGCTTAGATGGAAAGAAAGAGTTACTAGAGAAAAACTCGAATGGAAACATTGGGTAGAAATGAAAGAAAATATGAATATATATAATAAGTGGAAAAAATGGATAAAATGGAAGAAAAATAAATTAGCTAATTTTAATGAATGGTCAAAAAATTTTATAGAAAAATGGATACGAGAAAAACAATGGAACAATTGGATTAATGAAAGAAAAAAATATACATCTCAAAGAAAAAGTTTAGAACAACAATTTGGAGATAATATGGATAAAATGAACAAATTAAAAAAAAAAAAAATTTTGAAATTCTTCCCACTATTTAATTATAAAAGTGATTTGGAATCAATAATGGAAGAAGATGAAAACGAATACAACAGTTTTGATGAAAACGAAGAAGAAAACGATGAAAAAACAGGAGATGTTAATGTAGGAAAGACGGAAGCTTTGAATGTAGCAAAGACCGAAGGTTTGAATGTAGGAAAGACGGAAGATTTGAATGTAGCAAAGACAGAAGATTTGAATGTAGCAAAGACGGCAGATTTGAATGCAGAAAAAACAACAGATTTGAATTCAGAAAAAACAGCAGATTTGAATTCAGACAAAACAACAGATTTGAATCCAGAAAAAACAACAAATTTTAATACATACAAAGCAACAGATTTGAATGCAAACAAAACAGCAGATTTAAATTCAGACAAAACAACAGATTTGAATTCAGACAAAACAACAAATTTTAATACATACAGAACAACAGATTTGAATTCAGACAAAACAACAAATTTTAATACATACAAAACAGATTTGTATGCAGAAAAAACGACAGATGTTAATCTAGGAAAAACAACAAACCATAATGTAGCAAAAACAACAGATCAAAAGGTAGTAAAACATTCGTTAGATCACGAAGTAAGACAAATGATAGACCAAAAAGTAGCACAAATAATGAACCATGACTTAGAATCAACAGCAGAACAAAAAGCAGAAAAAAAAGGAGGAAAAGCAAAAGCGAAGACAAAAGTAAGAACAGTTGATGATGACGGAAATGAAATTAATGTT

>PeMP2

TAAccctttacattaatttcttagAACGTATCATCCAAATCGATATATAATAAAAATAAATTCCATAACACGTTCAATAGAAGAGATACAAGAGTTTTAGCAGAGCAAGAAGATCAATACATAAGGAACCCAAATAATTCTAATTATCCTGATAGAGACCTTGACATCTGTAATTGGGATGAACCTCATAATCCTGAAAAAAATCCTTGTGCCATTCCACAAGACGATTTATCCAATGAAGGCATAGAAATAATTGATTATGTTGATAAAAAAGAAGAAGAGAAACTTAAAAATGATTTGAAAATACATTTTGGCAAACAACACTATGCATTAATTAAGGATATATATAAACCTATACAATCTTATGCAAACAATTTTTCAAAGGTCGTAGATCATTATGTTGGTAGTTTAAAAAGAATTAACAATGCATTCAATTTAGATATTTTACGTACTGTTATATGCACAGCATTCCTTATACAACCAATAGTTTTATTAATGAAACTTCACCCTGAACTTCTTGTTCACATACCAATCAATATTATAATATCCATGGTTGTATTTAAATTAATGCATAAATCACTACGTAACGAACAAGAATCAAATAAA

>Pf332

TAAGTGAGAAATCTAAAATTGTGGCTTCAAGTAGCAAAACAACTAATGAAGAATATATTATTTTAAGCAGAAAAGATTTAATATACAATATAATTGTAATGGTTCATATGATGGTATTAGATCAATTTAAGTACGATGAACTTAAATATGCTAAGAAAAGATTTTTAAATAGATCAATTGATAAATTTATAAAAGAAAAAAAAATAAAAGACAAAGAAACAGTTTTGGATTATTATATAGATGATATTATAAAACGTATTGTAGAAGATACTTCACATGTTAATAATATTAAGGAAAAGGCAATAGATCATTACACTAGTCATGATTGGTTTAGATTATTACGCCAAGGGAATAAAGCTCAAAATTCCATTTCTCAGGAAGTAGACATATTAACGGAAAAGTATAAAAAATTAATTTCAGAGGAAGACAATAAAAAAGAAAATAACAATAATAATGATGATGAAGTAAAACCAGAACTAAAAGACAATATAAAGTGCGAAGAAAATGTCTCATTCAATATATTGAGAAGATCAAATAAAAAAGATCAATTACCGTTGGAAGAAAAAAAAAATAAAAATGGTGATTTGAACAATACATTAACTGAAAGTGATGGAAAAAAAATAAATATAAATGAAGCAATAGAAGAAAGTAAAAATGGTAATGAAGATAAAAATTCGGGAAATATGGAAAAATCTAGAAAACCTAGGGATAGGAGAACTGAAAGAAAGGAAAAGGATCAAGATTTAAGAGTTCAATTACTTCAAGATTATGAAAATATATTCGAACAAATTCAAAATATGGAAAATCAAAATAAAGATAAGAATAAAAAGGAAACTAAAAATATATTAAAAACATCAATTGATTTAGATAAAAATCTATTAAGAGAATATAAAAAGGAGAAAAATATAATAAACCAAAGTGAGAAGGAATTTAGTGTAAATGAGAAC

>PFC0070c

TAACAAAGTTTGTGGTATAAATTCCAAGAACCATATAATAATATATTAAATGTTAGATATTGTAGATTGTTAGAAGAACATGCAGAATATTTTGGTGAGGTGAAATCGAATGTAATCGTTTTAGTTAGTCAAAATTATCTTGATAATATGATAAATGAAGAAGATGATGAAATGTATGATGATATGGAAGATGAATATTATGATGTAAATGATGAAATTTTATTAGATGATGAAAGAACGAAAACATTTAATGTAGAAGATGAAGAAGTATTAAATGAAGAATTTTCAGATGCTTTAGATGATAATAATTTTTATATAAATAATTATGTGAAGGAAGATGTAGTTGTAGATGATATAAATAAAAATGCTGGAGCTGTGGAAAAAGAGAATTTAAATGATAAATGGTTAAATTTGATGAAAGAAAAATTAAAGGATATAAATTATTATAAAGAAAAATATGAAAATAAAAGACATAATTATAATGATAGTGTTAAAAAGTTAGAAGAAAAATATCATTTAACATTTCTATGGGTCTTGGGTTACATATTTTATTTTATTCCATCCCTTGCTAAATTATTTCCTATATTATATCATTTTTGTATATATAAATTTAGTCAAATGATGTCAAATCCA

>PeMP3

TAAATCTTTACGAACTTCAAAATATTATCCATAGAAATACTAGTGTTTTTATTGCTACTATTACATATTgtaagaataataattaatataaattaatctatatatatatatattatattttgttataattatatattaatataatgaaaacgcaaaattttccttactaaatttaaatataatttttttatacttcttataacatgcagGATAAGCCTAAAAATGTTAATCTTGTAAAAACGGATATATACAATGGATTTGTTCAGGAAAATGCAAGATATTTAGCGGAACAATATATTAATAAAAAAATGGATCAAACTTTACAACATAGAGCACAACTCAGAACAGCCCAAAATGTATATAGAAAAAAGCAACAAAAAAAAAAAAAAAAGAAATTACCAAAATCGGAAAGAAGAAATGCAGAAGAAAGAGCATTAGCAATAAATGATTATATAGATTACTTAAAGGAAACCTGCAACGGAAGTGAAAAACTCATGAGACTTAAATATATTTCATACCTAATTACATATATGGATATAACATATAAACAAAAAGAAAAGCTTCTAGATCTCGTTGAGCAATATATATATAGTAACATTGAACATGAGAAAAAAAAATTATCTAAATTTATAGATCCTTATATAAATAAAGAAGACGATGAAAAATCTAAAATTAATTTTCAAAATATTCTTTATGAAAAAAAAAGGATAGAATCCTTAAAATATTCTGAATTAGATTTGTTTTTATACACTATATTTCCCTACATG

>PTP5

TAACCATAGTGAAACTTCATCACATCAAGGGGAAAAGAATAATGAAAATGAAACTGAAAAGAAAACTGATCAAAATGAAACTGGAACAAAAAAACCTTCTAAATATACAATGAATCTTGATTCTCCTCTGTTAAAAGGTTCAAGTGAAGGTGAAACTTCATCCAAAAAAGCACAAGAGAAAAGTGTAGAACCAACTAAAAAACCTTCTAAATATACAATGAATCTTGATTCTCCTTTGTTAAAAGGTTCAAGTGGAAGTGAGACATCATCTACCAAAGAATCAAATGATAGTAATGGAGCAACCAAAAAGCCATCAAAATATACAATGAATCTTGATTCTCCTTTGTTAAAAGGTTCAAGTGGGGGTAAAACATCATCTAATAAAGAATCAAATGAAAGTAATGAACCAACTAAAAAGCCATCTAAATATACAATGAATCTTGATTCTCCTTTGTTAAAAGGTTCAAGTGGAGGTGAGACATCATCTAACAAAGAATCAAATGAAAATAATGAACCAACTAAAAAGCCATCAAAATATACAATGAATCTTGATTCTCCTTTGTTAAAAGGTTCAAGTGAAGGTGAAACATCATCTAACAAAGAATCAAATGAAAGTAATGAACCAACTAAAAAGCCATCTAAATATACAATGAATCTTGATTCTCCTTTGTTAAAAGGTTCAAGTGGAGGTGAGACATCATCTAACAAAGAATCAAATGAAAATAATGAACCAACTAAAAAGCCATCAAAATATACAATGAATCTTGATTCTCCTTTGTTAAAAGGTTCAAGTGAAGGTGAAACATCATCTAACAAAGAATCAAATGAAAGTAATGAACCAACTAAAAAGCCATCTAAATATACAATGAATCTTGATTCTCCTTTGTTAAAAGGTTCAAGTGAAGGTGAAACATCTTCTAACGAAAAGCAAGAGGAAAGTAATGTTGCAACTAAAAAACCTTCTAAATATACATTGAACCTTGATTCCCCTCTTTTAAAAAGCGAAACAAAATCAGATGTAAAACGTGGATCAAATAAATCATTTAGTCTAGAAAATATTGGAAAACTCGATATAGGTTCTCTACTTGCACAAAATTTAGAATTACTAAAAGGATTTGCGTTGAATTTTCAAACATTGAGTTTATTTTTCTTGGTGTTTGTTGGACAATTTTATCCTAAACATTTTCAGAAGGCTGCAATATTCATTGGAATGGTAAATGTATTTTTAGAATGTAAACGTTTATATGGTAGATCTCAAAAGAAATTAAAA

>PFL0055c

TAAATCATAGTGTTAATGCAGTATCACCCGTTAATACGACATATTATGAAATATTAAATGTAGATACGAAAGCAGAATTAAAAGAAATTAAAAGTAATTATTATCATTTATCATTACAGTATCATCCTGATTATAATATAGGTGATCGTATAGCTAAATTGAAATTTCGACTTGTCAGTGAAGCATATCAAGTATTATCTGATGACGAACGAAGAAGGATTTATAATAAGCAAGGATTAAAAGCAACAGAAAAAATGTTTTTAATGGAACCTGGTCTTTTATTTATGATAATGTTTAGTATAGATGAAATGTCTGATTATGTTGGTGATTTAAAATTGTTTTATTTTATTAAAGAAGCTTTTGAAAAGAAAAAACGAATAGAAGATATAGAATCTCCATTTGAAGATATGGATGCAAAAATGGAAAATGACCAAAGAAAAAGAGAAGTTGTTTTAGCTCTTTTGTTAAGAGAAAGAATACAACCATATGTAGATAATAATAAAGAATGGATGTGCGAGATGGAGAAGGAAATAAAAAGCTTACTAGAATCCTCTCATTCAAATGCTATTTTAGGATCCATAGGATGGACATATGAAAATGTTGCAACTAAATATTTAAGTGATATAAAATCAAAATGGAGATTAAAAGAACCAATGTCAAAATATGATGCATCTTTTAGACATGTAAATAGATCTAAAAGAACAAAAATGACTAAGTTGTCATCTAGGTTTTCTGGAATGTTTAGTTGTTCTTCTGCCTTAAAATCGCAAGAACCTTCTATGGGAGAAACAAGTTCAAGTGATAACGATCGTAATGAAGGAAGTGAAGGTATTTGTAATATGGATAATATGTTTTCCGAAGAAACATTATCCTTGTTAGAAATAGATGAGAATAAATATTTTGGATTTATGACAATGCATATTTTAACATTAATTTTATGGGATATAGAAGAAACTACTCAATATGCTGCTAGTAGAGTTTTAAGAGATGAGGGAGTTGATGAAAATACACGAATAAAACGAGCAGAGGCATTACAAATTCTAGGAAAATTAATGCAGAAATGGTCATTAAATGTTAAGAGTAAAAAGGAAATACGTGAAAAGGATATAATTGAAATTATGGAAAAGGCAAGAGCTTATAATATG

>XL1

TAAAGTTGGATTGAAAAATTAAATGAGAATGGTTTCACATTTATTGGAATAGATAATCAATCTCATGGTTTATCAGATGGAGCTCGAAATCAACGATGTTTTGTTGAAAATTTTGATAATTTTGTTTTCGATGCTGTTAAAGCGTTAGAAATGTTTATAAATGAATGCAAAGAGAAAAATGAATTAAAACCTATAATAATTATGGGAACATCAATGGGTGGATGTATAGCTTTAAGAACGATTGAAACTATATATAAATTAAATAAAAATTGGAAAGATAATATTAAATGTTTAGCTCTTATTTCACCTATGATAAGTATTGAGAATCAAAAGCGTAAATTAATTAATAGAATATTATATAGTATATGTAGATACGTTAAAAAATATTTTGAACTTTATGAAATGGATGTTTTATATGAGAGACCTAAATATCCTTGGATTAAAGGTGACACAGACATTGATCCTAATCATCATTCCGAAGGTTTGAAACTTGGAACAGCTGCAGAATGTGTATTCGCAGCTGATAAATGTTTAGTTCATTCAATATTAAAATATATAGAAGAGAGTGATATCGACATTATTATTCTTCAATCCAAATATGATACTACTGTTGATCCTACAGGTCCTATTGATTTTGTTAAAAAAATGATAGATTTGTACAACAAAAACCATCATGAAAATTCTAATGCTAAAGAGGATGATAATGAAAGTGAAAAAAAAGAATTAGAATCTCATAATGAAACGGAAGTGTCTAATAATGATCAAGAAAAAAAAAATATATCATTAAGAGATGAAAACATACCTAATGCTGATGACATTCAAAATGATATAAATGGATCATGTTATAATGTAGAGAATTATGTGCTTGTATCGGGAAATGATTTAACAGAACAACAAAAATTATGGCATTCATGTGATCATGGATATTATAAAAACTTTTTAATGAAAAAATTGGGTAAAAGCAACAATTTGAAAAGCAAAAATGGGGAAGATAGTTTTAAACGTTTAAGTGCTCATATATTGAATTATGGTTCCCACAGGTTACCCTGCGAACCAGATACAGAACGAAGTATAAATATTATAACTGACTGGATAAATAATATATTTGTA

>XL2

TAAGGTTTATCAGATGGGTTCAAAGATGAACGTTGTTTTGTCGATGATTTTGAAAACTTTGTTTTAGATGCTGTTCAAGGTTTAGAAATATTTATAAAAGAGTTGAAAGAAAAAAATGATGTAAAACCAATAATAATTATGGGTACGTCTATGGGTGGTTGTATAGCTTTAAAAATGTTTGAATACATACATAAATTAAATAAAGAATGGAAGTCACAGATTAAAAGTTTAGTACTTATTTCTCCTATGATTAGTGTGGAGAGACAGAAAAGTAAATTTACTAATAGAATTTTAATAAGTATAGGTAATATACTTAAGGGATTCCATCCTCTTCTAACGCTTGATGTAAAAGAAGGTGGTTCAAAATATCCATGGATTAAATATGATTCTGAAATCGATCCTTTTCAATATTCTGAAGGATTAAAAGTTGGAATGGCGTCCGAATGTGTAAAAGGTGCTGATAATTGTATGAAACATAATATTCTAAAATATATAGATAAAAGTAATGTTGATATTATGATTCTGCAGTCAAAATATGATGTGACGGTTGATCCTACAGGTACTGTTAATTTTATGAAAAAAATGATAGATTTATACAACAGTAAAATTGACAAAAATTCGAATAATAATAAGGATATTAATGATACAAAAAATAAAAAGAATGTAGAAAGAACTAATTTAAAAGGATTTAATAAAAATAAAGAGAAGAAACAAGTATCAGCTGATGACAAGGATGTATCTAATTCTAGTGAATTATTGGATTTGGATAATGAGCCATGTAAGCATTTAGAAGATTATACAGTTTTAGAAGGTAATGAATTAGAACCTGACAAACATTTATGGAAATCATGTAATCATGGATATTATAAGAATTTTAAAAAGAAAATAAAATCCGAAAAAAATAATAAAAATGATTCTGATAAGGAAGACAAGAAGTTTAAACACTTAAGTGCTCACATATTAAATTATGGTGCTCATAGTTTAGCATGTGAACCAGAAACAGCAGAAAGTGTTGATATTTTAGTTAATTGGCTTAATAATATATATCCA

>PeMP4

TAAttattagCATAACACCAATAAATATTGCAATGATAAACATCATTTTATAAATAACTCATATAATAAATTACTAAATAGATCATTAGCAGAAGTTAATGAAAACAATAGTTCGTCGATCAAAAATAAAGTAAATCAGAATGTAACTCGGTTAAGACGAGAAAAACCTTATATTGATGACAATAATAAAAAATGTGATAAAAATGAACATGATATAACACAAAAATTCACGTGTATCGATAATACATATAAGGAATCGAAAATTAAACCTACGAATTTAGAGCAGTTAATATTTAAAAAAAAAGAAGAAGAAAAACATTTTGACACAAGAAATGATTTATCATTTATTGAAAAAATTAAATTCGTATGTGATGCATTTGATGACCTTTTCATAGAAAAAATATTAGATTTTCATCTTCAAAAAAAGACTTCAGAATTCTTGAGAATAACAATGGAAACTCTTGTAGCACAATATTTAGCATTTTTCCCTTTTTCATTAGCAAGATTTAGATATTTAGTAGATCGAATAAACTTTTTTAATGTACGTCATTGTGGAAAAAAAAAAAAAGATATATTTTACATAACAGAAGATAAAAAAGATCTTTACGGTGTTAAA

>GEXP18

TAAATATATATAAGGAAGGATAAATTACGTTTTTGTTTTAGTTTTTATTTTTATGTGCAACTTTTTATAATATATTTGTTCATATGGACAGAGAATAAATATAATGAAgtaaaataaaaataacataaaatatatattatatatattatatatattatatatattatatatatattatatttatatatgtggtttatttttttatttttatttttatttttattttttttttagTATGGAGGAGATAAAAATTTGAATGGTAAATTAGATTTTAAGAGGTCCCAAAGATTGAAGGAATATAGGATATTAGTTGAATTTTCAAATAGTTATTATTATGATGAGCCGAAAGTAAGAATTATTAATTATGATGATGATAAGAATGAAGAATATCCTTCAAGATATAATAATAAGAATTTAGAAGAATCTTTTGAAAAATATGTTGATAATTATGAATATAATGTGAGGGATCATATATCAGAAAATGTAAAATTAGGAAATGATTCTATAGGGGATAACATAAGTATTGTTGATTTTAATGGAGAAAATAATGTGGAGGTAACTTTTCCTGACGATAATATAAATGAAAAGAATAGAAATAATGATAAATTGGATGAACTTAAATCATATATAAATACAAATGATGATGAAAATCAAAAAAAATTATGGAACGAGTATATAAATAATATTGATTATATAAAACCAAAAAGTATAGAATATAAATTTAAATATCATACATCTACAAAAAGAAATTTAAGAAGTTTTGTTAAATATGCAAAACGTTCTATTAGATTAGTATGTAAAATAATTAAATTCATTTTAATTACTATTTATCGTTTAATTAAATATATTATTGTTTGGATTTTTAACACAGCAGATTCTGTAGTAGAA

TGD contructs

GFP

Neomycin-R

>pArl1-**Insert**-GFP-T2A-NeoR

gcggccgc**Insert**acgcgtGCTAGAGGTGCTGCTGCTGGTGCTGGAGGTGCAGGTAGACGTACGATGAGTAAAGGAGAAGAACTTTTCACTGGAGTTGTCCCAATTCTTGTTGAATTAGATGGTGATGTTAATGGGCACAAATTTTCTGTCAGTGGAGAGGGTGAAGGTGATGCAACATACGGAAAACTTACCCTTAAATTTATTTGCACTACTGGAAAACTACCTGTTCCATGGCCAACACTTGTCACTACTTTCGCGTATGGTCTTCAATGCTTTGCGAGATACCCAGATCATATGAAACAGCATGACTTTTTCAAGAGTGCCATGCCCGAAGGTTATGTACAGGAAAGAACTATATTTTTCAAAGATGACGGGAACTACAAGACACGTGCTGAAGTCAAGTTTGAAGGTGATACCCTTGTTAATAGAATCGAGTTAAAAGGTATTGATTTTAAAGAAGATGGAAACATTCTTGGACACAAATTGGAATACAACTATAACTCACACAATGTATACATCATGGCAGACAAACAAAAGAATGGAATCAAAGTTAACTTCAAAATTAGACACAACATTGAAGATGGAAGCGTTCAACTAGCAGACCATTATCAACAAAATACTCCAATTGGCGATGGCCCTGTCCTTTTACCAGACAACCATTACCTGTCCACACAATCTGCCCTTTCGAAAGATCCCAACGAAAAGAGAGACCACATGGTCCTTCTTGAGTTTGTAACAGCTGCTGGGATTACACATGGCATGGATGAGCTCTACAAAGTCGACGGAGAAGGAAGAGGAAGTTTATTAACATGTGGAGATGTAGAAGAAAATCCAGGACCAATGATTGAACAAGATGGATTGCACGCAGGTTCTCCGGCCGCTTGGGTGGAGAGGCTATTCGGCTATGACTGGGCACAACAGACAATCGGCTGCTCTGATGCCGCCGTGTTCCGGCTGTCAGCGCAGGGGCGCCCGGTTCTTTTTGTCAAGACCGACCTGTCCGGTGCCCTGAATGAACTGCAGGACGAGGCAGCGCGGCTATCGTGGCTGGCCACGACGGGCGTTCCTTGCGCAGCTGTGCTCGACGTTGTCACTGAAGCGGGAAGGGACTGGCTGCTATTGGGCGAAGTGCCGGGGCAGGATCTCCTGTCATCTCACCTTGCTCCTGCCGAGAAAGTATCCATCATGGCTGATGCAATGCGGCGGCTGCATACGCTTGATCCGGCTACCTGCCCATTCGACCACCAAGCGAAACATCGCATCGAGCGAGCACGTACTCGGATGGAAGCCGGTCTTGTCGATCAGGATGATCTGGACGAAGAGCATCAGGGGCTCGCGCCAGCCGAACTGTTCGCCAGGCTCAAGGCGCGCATGCCCGACGGCGAGGATCTCGTCGTGACCCATGGCGATGCCTGCTTGCCGAATATCATGGTGGAAAATGGCCGCTTTTCTGGATTCATCGACTGTGGCCGGCTGGGTGTGGCGGACCGCTATCAGGACATAGCGTTGGCTACCCGTGATATTGCTGAAGAGCTTGGCGGCGAATGGGCTGACCGCTTCCTCGTGCTTTACGGTATCGCCGCTCCCGATTCGCAGCGCATCGCCTTCTATCGCCTTCTTGACGAGTTCTTCTAACTCGAGggatatggcagcttaatgttcgtttttcttatttatatatttataccaattgattgtatttataactgtaaaaatgtgtatgttgtgtgcatatttttttttgtgcatgcacatgcatgtaaatagctaaaattatgaacattttattttttgttcagaaaaaaaaaactttacacacataaaatggctagtatgaatagccatattttatataaattaaatcctatgaatttatgaccatattaaaaatttagatatttatggaacataatatgtttgaaacaataagacaaaattattattattattattatttttactgttataattatgtgtctccttcaatgattcataaatagttggacttgatttttaaaatgtttataatatgattagcatagttaaataaaaaaagttgaaaaattaaaaaaaaacatataaacacaaatgatggtttttccttcaatttcgatatcaatttatagaaacaaaatatatacttgtataattttatttttttatataaatcattacatatataattatacaatattttttctaagagataattatatattaatatatataaaaaaaggtgttttttttttttttttttatttttatttttattttatggtaatattttattttccttattttataaattatattagtttatatgtgattaattttatatattatcaatttatatatttttaaatgcttacttaattatctttttttttttttttttttttttttcccctctttttatattaatttatttttgaaaaaattgatatatatatatatatataatatatatatatacatgtagtagtattaaacaatgtataatatatataaataatatatttatatatttcatttcaattttaattttttttggttttttttttttttctttttgtcatatttaaaaaaaattatattcatataagttatgcattttttataaacattattcaatatatgtataatataatatatatatatatattaatgtattattccaatgtgcatgataaaagaaaaaaataatatttataaaaaaaaagaaaaataaaacaaaaaaagaaaaaaaaaaaaaaaaaaaaaaaaatacaaaaataaataatataatttataattatatattcttgtcacaataaaaatatatatatatatatatatatttataatatgtatattttaaactagaaaaggaataactaatattttatttattatcattcaagatttatattttataataataaatacctaatagaaatatatcaggatccatgcatggttcgctaaactgcatcgtcgctgtgtcccagaacatgggcatcggcaagaacggggactacccctggccaccgctcaggaacgaatttagatatttccagagaatgaccacaacctcttcagtagaaggtaaacagaatctggtgattatgggtaagaagacctggttctccattcctgagaagaatcgacctttaaagggtagaattaatttagttctcagcagagaactcaaggaacctccacaaggagctcattttctttccagaagtctagatgatgccttaaaacttactgaacaaccagaattagcaaataaagtagacatggtctggatagttggtggcagttctgtttataaggaagccatgaatcacccaggccatcttaaactatttgtgacaaggatcatgcaagactttgaaagtgacacgttttttccagaaattgatttggagaaatataaacttctgccagaatacccaggtgttctctctgatgtccaggaggagaaaggcattaagtacaaatttgaagtatatgagaagaatgattaagcttatttaataatagattaaaaatattataaaaataaaaacataaacacagaaattacaaaaaaaatacatatgaattttttttttgtaatcttccttataaatatagaataatgaatcatataaaacatatcattattcatttatttacatttaaaattattgtttcagtatctttaatttattatgtatatataaaaataacttacaattttattaataaacaatatatgtttattaattcatgttttgtaatttatgggatagcgattttttttactgtctgtatttttcttttttaattatgttttaattgtattttatttttattattgttctttttatagtattattttaaaacaaaatgtattttctaagaacttataataataataaatataaattttaataaaaattatatttatcttttacaatatgaacataaagtacaacattaatatatagcttttaatatttttattcctaatcatgtaaatcttaaatttttctttttaaacatatgttaaatatttatttctcattatatataagaacatatttattaaatctagaattctatagtgagtcgtattacaattcactggccgtcgttttacaacgtcgtgactgggaaaaccctggcgttacccaacttaatcgccttgcagcacatccccctttcgccagctggcgtaatagcgaagaggcccgcaccgatcgcccttcccaacagttgcgcagcctgaatggcgaatggcgcctgatgcggtattttctccttacgcatctgtgcggtatttcacaccgcatatggtgcactctcagtacaatctgctctgatgccgcatagttaagccagccccgacacccgccaacacccgctgacgcgccctgacgggcttgtctgctcccggcatccgcttacagacaagctgtgaccgtctccgggagctgcatgtgtcagaggttttcaccgtcatcaccgaaacgcgcgagacgaaagggcctcgtgatacgcctatttttataggttaatgtcatgataataatggtttcttagacgtcaggtggcacttttcggggaaatgtgcgcggaacccctatttgtttatttttctaaatacattcaaatatgtatccgctcatgagacaataaccctgataaatgcttcaataatattgaaaaaggaagagtatgagtattcaacatttccgtgtcgcccttattcccttttttgcggcattttgccttcctgtttttgctcacccagaaacgctggtgaaagtaaaagatgctgaagatcagttgggtgcacgagtgggttacatcgaactggatctcaacagcggtaagatccttgagagttttcgccccgaagaacgttttccaatgatgagcacttttaaagttctgctatgtggcgcggtattatcccgtattgacgccgggcaagagcaactcggtcgccgcatacactattctcagaatgacttggttgagtactcaccagtcacagaaaagcatcttacggatggcatgacagtaagagaattatgcagtgctgccataaccatgagtgataacactgcggccaacttacttctgacaacgatcggaggaccgaaggagctaaccgcttttttgcacaacatgggggatcatgtaactcgccttgatcgttgggaaccggagctgaatgaagccataccaaacgacgagcgtgacaccacgatgcctgtagcaatgccaacaacgttgcgcaaactattaactggcgaactacttactctagcttcccggcaacaattaatagactggatggaggcggataaagttgcaggaccacttctgcgctcggcccttccggctggctggtttattgctgataaatctggagccggtgagcgtgggtctcgcggtatcattgcagcactggggccagatggtaagccctcccgtatcgtagttatctacacgacggggagtcaggcaactatggatgaacgaaatagacagatcgctgagataggtgcctcactgattaagcattggtaactgtcagaccaagtttactcatatatactttagattgatttaaaacttcatttttaatttaaaaggatctaggtgaagatcctttttgataatctcatgaccaaaatcccttaacgtgagttttcgttccactgagcgtcagaccccgtagaaaagatcaaaggatcttcttgagatcctttttttctgcgcgtaatctgctgcttgcaaacaaaaaaaccaccgctaccagcggtggtttgtttgccggatcaagagctaccaactctttttccgaaggtaactggcttcagcagagcgcagataccaaatactgtccttctagtgtagccgtagttaggccaccacttcaagaactctgtagcaccgcctacatacctcgctctgctaatcctgttaccagtggctgctgccagtggcgataagtcgtgtcttaccgggttggactcaagacgatagttaccggataaggcgcagcggtcgggctgaacggggggttcgtgcacacagcccagcttggagcgaacgacctacaccgaactgagatacctacagcgtgagctatgagaaagcgccacgcttcccgaagggagaaaggcggacaggtatccggtaagcggcagggtcggaacaggagagcgcacgagggagcttccagggggaaacgcctggtatctttatagtcctgtcgggtttcgccacctctgacttgagcgtcgatttttgtgatgctcgtcaggggggcggagcctatcgaaaaacgccagcaacgcggcctttttacggttcctggccttttgctggccttttgctcacatgttctttcctgcgttatcccctgattctgtggataaccgtattaccgcctttgagtgagctgataccgctcgccgcagccgaacgaccgagcgcagcgagtcagtgagcgaggaagcggaagagcgcccaatacgcaaaccgcctctccccgcgcgttggccgattcattaatgcagctggcacgacaggtttcccgactggaaagcgggcagtgagcgcaacgcaattaatgtgagttagctcactcattaggcaccccaggctttacactttatgcttccggctcgtatgttgtgtggaattgtgagcggataacaatttcacacaggaaacagctatgaccatgattacgccaagctatttaggtgacactatagaatactc

**Inserts used:**

>ΔPIESP2

TAATTACTCTTTTTTGCAAAACTTGTCGTATTTACCTTTTTCTTTTGGCTTTTAAAATATGGGAAAACGgtaagacgtacatgatattcagaataatatatatatatatatatatgtatgtttttatagtacaattaatacgccttttataacgcttataaagaataaaaatctttatatatatatatatatatatatatttgtatgtatttataatttacctttaatcattgttatattaattttttattttttattttctttcagAGGTCATATCCCAAATCTGGCCATAAGGGACATACGAAATTAAATCAACCAGTAGTTAGAACATTAGCAGATTTTAATGACATGTTTGCAAACCAAAAAAATACATTTAATTTTCTAAAACATATAAATCATTATAAAAATGAACAAGATACAAATAATACACACACGCCAAATCATGATGAATATTCTCATAATTTGCCAAAAAATCACGAAGAGTCAAATGCAAATATGAACAATCATAATTCTTTCAATGACAAATCTGTTAATAAAAAAGAAGCTTTCGATCAATTTTTACAAACGTTATTAAACAATTATGAAATAATGCATAAAGAAGATGAAAGTAAAGAATCAAATCAACATAACTATAAAGAAGGTCCCTCATATGAA

>ΔPf332

TAATCTAATATAAATAACAAAGACTCTAGTACAGAATGGAATTGTAAAGAAGATGTGGGATGTGTTCCACCTAGGAGACAGAATTTGAATATGGAAAGGTTGGATAATGAAAATGAAGATTCTGTACCCGATTTCATGAAGAAAACTTTTTATCTTGCTGCTGCTGGAGAAGGAAAGAAGTTACGTGAGAAGCATGATGAGAGTTGTGATGAATTCTGTGACGCATGGAATAGAAGTTTAGCTGATTATAAAGATATATTTCAAGGAAAGGATATGTGGAATGATGGGAAATATGGTGAAGCGAAAAATCATATTAAGAATGCTTTTGGTGATATGAACAATAGAAAAACTATGTTAAATGAAATTGAGAAAGGAATTAAAGATGAAACGTTTAGTCGTGAAAATGGTTTAGACGTTTGTAAATCTCAATGTGAGGAAAGAAGTAGAGATGACACAGAAGATCAATTTTTGAGGTTTTTTGCAGAATGGGAAGAAGAATTTTGTGATGGG

>ΔPeMP1

TAAGTTCATTTTACCAAGATTTTAGCAGTTACCTGTTTTGCTGCTACTTTTCAATGTTCTAATAATgtaagtttataataaaatagattataaaaaaaaaaaaaaaaaaaagaaaaaaggaaaaggaaagaaaaatgcatatatatatatatatatatatatttatataatataatacaatacaatatgtagatattttaatttacctttttgttatatacttttttattttttttaattatttagGGAGTTCTTTCCAAGAGTTATGATATGAATAAGAAGAATTTGAATAATGTAGGATTCCGAAATAACAGAATTTTGTCTAGTAAGGAAAACCGTGAAGAACCAGGAACTAGTGGTACAACATCAGGTGCTAATGGTAACAAGGCTAATATTCCTGACCTTGATAATTTTATAAATCAGACAACATCTTCATGTGGTAATTTGGGTAATTTGTTAGCACCATTACTTAAGGGTATTAAGTCTAATTTATTTTCAAAATATGTTAATGAAGATGGTACAACTAAACCAAATGCTCCTACAGCGCAGAGTTTAACCTCTATAGTTAAAGGA

>ΔPeMP2

TAAAATAAAAAATCAATGCAAACTAAGAACTTTTTATCTGAAAGGAACTATGGAAGTATAGATCAAAATGTGAGGACTAAAAATAAAAGAAGATTAATGAAATTCCAAAGTAAGAGTAAAGCAAAATCGTTCCTTTTTTTATTGGAACTTATGGTATTCTCCCTTTTCATATGGATTTTAAAGAGTGCAAAGCATgtaagtttatattttctgtatatatatatatatatatatatatatatatttatttatttatttatttatttatttatttatttatttattgatttattcatgtacacctatatttgtattgctctacataaaacgataaaaaattatgttatccctacataaaaataatgattttgtatttctatccataataactacctatatataaacacatatttctttatttatccctttacattaatttcttagAACGTATCATCCAAATCGATATATAATAAAAATAAATTCCATAACACGTTCAATAGAAGAGATACAAGAGTTTTAGCAGAGCAAGAAGATCAATACATAAGGAACCCAAATAATTCTAATTATCCTGATAGAGACCTTGACATCTGTAATTGGGATGAACCTCATAATCCTGAAAAAAATCCTTGTGCCATTCCACAAGACGATTTATCCAATGAAGGCATAGAAATAATTGATTATGTT

>ΔPeMP3

TAAATCTTTACGAACTTCAAAATATTATCCATAGAAATACTAGTGTTTTTATTGCTACTATTACATATTgtaagaataataattaatataaattaatctatatatatatatattatattttgttataattatatattaatataatgaaaacgcaaaattttccttactaaatttaaatataatttttttatacttcttataacatgcagGATAAGCCTAAAAATGTTAATCTTGTAAAAACGGATATATACAATGGATTTGTTCAGGAAAATGCAAGATATTTAGCGGAACAATATATTAATAAAAAAATGGATCAAACTTTACAACATAGAGC

>ΔPeMP4

TAATTTATGAAAAAATTTATGAAGGCAAACAATAAACGATTCTATTACAGGTGTACCAATTATGTTTATAAAATCTCTAAAAATGAAGATACAAGTATATTATTATATAATTATATGTTGAATAATATACAACAATATTTCAAATTTCTTAAATATTTATTATTTTTTCATTTACTTTTATTTACATATATATACAACAAGgtagatataaaatattaaaacaaaagtgaaacatgatttattatataattataatttcaaatataaatgaaaaatatatatatatatatatatatatatgtatttcatttcaatattttaatatattatataaatgttatacattataatatttttatttttttttatttttattagCATAACACCAATAAATATTGCAATGATAAACATCATTTTATAAATAACTCATATAATAAATTACTAAATAGATCATTAGCAGAAGTTAATGAAAACAATAGTTCGTCGATCAAAAATAAAGTAAATCAGAATGTAACTCGGTTAAGACGAGAA

Episomal expression contructs for Maurer's clefts analysis

*MSRP6*

mCherry

>pArl2-crt-MSRP6-mCherry

ctcgag*ATGAAAAGCAAAAAAATAATATGTTCATCTTGCTTATTTTTAATATTTTTAAGTGTAATATTTTGTAGTGAACCAGATACAAATTCATTTGATGAAAATGTAAAGAAGAATGAAGTTTTTAATGCCTTAAATGAACATTTAGAAAGTATAAGTAATATCGTAAAAGTAAATATTATGGATGCCCTTTCAAATAACCCTTCGTTAATAAAAAAGACATACGAAGCTGTTGAAATAAATGATGATGATTATGTTCTGGAATATGTTGGGGATTCGACAGGAAAATATGGAGAGGGCTCTATATTCTATGATGAAAGTAAGAAATATAATTATAGAAAGTATTTAGATATAGAAAATGAATTACGAAATATGAAAGGTGAAGATGATGATGATTTTGATGAAGATGATGATGATTTTGATGAAGATGATGAAGATGATGAAGATGATGAAGATGATGAAGATGATGTAGATGATGAAGATGATCTAGATGTTGAAGATAATGTAGATGATGAATATGATGATGAGCATAATCATAATTATAATGATGACAAATTGAGTGAAAATCCTGAGAAGTATTCAAATTATAATAAAAATATACACGAAGATAAGAAAAAAGATAATTTGAATGAACCACATTTTAAACAAACTCATTATATATATTCATCAAATCACGATAATAATGAAACATCTAGATTTCCAAAAAAAAATGTACCTAAATATGATGAAAAATTAAATAATGAATTTAAAACATATTTAAGAAGACCTGAAAAGAAAGAAGAAACAAAAGAATATCCTAACAATGGATGTTCTGTAGTTCAAATATCTATAGTAACTAATGAAGATTTTTTAAAAAAAGTTAGAGAAAGAAATAAAAAGAGAAATAATAAAAAAAGAACAAATATTTATGATAGTGATGAAGAATCAGAAAGTTCAGAAGAAACAAGTAAAGATCCCTATTCATCGGGTCCATATACAGTTGATCATAAAAATGAAAATTGTTTTCCTTACAAGCAAAATTCAGATATTCATCCAAATATAAATAATGATAATAATGATAATAATGAAAATAATGATAATAATGAAAATAATGATAATAATATACCTGTTAATAATAAAAAAAATTCTCATACTCCTCATGTACCAAAAGATACCAAAAATAATATAGAAAAAAATCATAGTAACTATAATTACAAACCAATTAATCAAGAAAATTTATCAATGTATTTTAATAATAATAATAAGAAGAAGAATAAAAAGAATAGTCAAAAAAATAATCAAAAGAATAATCAAAAGAATAATCAAAAGCATAACCAAGATCTTTATAATATAAAAAAAAATAATGATCTACCTCAATATGAAGAATCTAATAAAACTTATAAAAAAATTTTTTCAAATGAACATGAACCACGTCCGATGATAACAACATCCAGATATGAAAATAAAATATCAAATGATAATAACAACAATAACTATAATTTGAATGAACAAGAAAGAAAACTTATTAAAAATATGATAGATATATTTATTTATACATACAAGTTAAATTATACAAACAGTAAATCTATATCTAAATTATTTAAAAATAATTTACTTAAAAAAAACTTCCGTACTTATTTCACAAATTATATATATACACTTTTTAATTATGGAAAGACATATAATTTTCTAACACCTTATAATAAAGATAATGATCATATGTATAGACAATTATTTGATGAAGCTGTACAAATGATGGATCTACTTATTAATAAAATGGACTTAGCTTTAAATCCCACGAAATTA*actagtacaagttatccatatgataatccagattatgcAccagttgcaacattaggtaccatggtgagcaagggcgaggaggataacatggccatcatCaaggagttcatgcgcttcaaggtgcacatggagggctccgtgaacggccacgagttcgaGatcgagggcgagggcgagggccgcccctacgagggcacccagaccgccaagctgaaggtGaccaagggtggccccctgcccttcgcctgggacatcctgtcccctcagttcatgtacggCtccaaggcctacgtgaagcaccccgccgacatccccgactacttgaagctgtccttcccCgagggcttcaagtgggagcgcgtgatgaacttcgaggacggcggcgtggtgaccgtgacCcaggactcctccctgcaggacggcgagttcatctacaaggtgaagctgcgcggcaccaaCttcccctccgacggccccgtaatgcagaagaagaccatgggctgggaggcctcctccgaGcggatgtaccccgaggacggcgccctgaagggcgagatcaagcagaggctgaagctgaaGgacggcggccactacgacgctgaggtcaagaccacctacaaggccaagaagcccgtgcaGctgcccggcgcctacaacgtcaacatcaagttggacatcacctcccacaacgaggactaCaccatcgtggaacagtacgaacgcgccgagggccgccactccaccggcggcatggacgagctgtacaagCCCGGGTCGAGGGATATGGCAGCTTAATGTTCGTTTTTCTTATTTATATATTTATACCAATTGATTGTATTTATAACTGTAAAAATGTGTATGTTGTGTGCATATTTTTTTTTGTGCATGCACATGCATGTAAATAGCTAAAATTATGAACATTTTATTTTTTGTTCAGAAAAAAAAAACTTTACACACATAAAATGGCTAGTATGAATAGCCATATTTTATATAAATTAAATCCTATGAATTTATGACCATATTAAAAATTTAGATATTTATGGAACATAATATGTTTGAAACAATAAGACAAAATTATTATTATTATTATTATTTTTACTGTTATAATTATGTGTCTCCTTCAATGATTCATAAATAGTTGGACTTGATTTTTAAAATGTTTATAATATGATTAGCATAGTTAAATAAAAAAAGTTGAAAAATTAAAAAAAAACATATAAACACAAATGATGGTTTTTCCTTCAATTTCGATATCAATTTATAGAAACAAAATATATACTTGTATAATTTTATTTTTTTATATAAATCATTACATATATAATTATACAATATTTTTTCTAAGAGATAATTATATATTAATATATATAAAAAAAGGTGTTTTTTTTTTTTTTTTTTATTTTTATTTTTATTTTATGGTAATATTTTATTTTCCTTATTTTATAAATTATATTAGTTTATATGTGATTAATTTTATATATTATCAATTTATATATTTTTAAATGCTTACTTAATTATCTTTTTTTTTTTTTTTTTTTTTTTTTCCCCTCTTTTTATATTAATTTATTTTTGAAAAAATTGATATATATATATATATATAATATATATATATACATGTAGTAGTATTAAACAATGTATAATATATATAAATAATATATTTATATATTTCATTTCAATTTTAATTTTTTTTGGTTTTTTTTTTTTTTCTTTTTGTCATATTTAAAAAAAATTATATTCATATAAGTTATGCATTTTTTATAAACATTATTCAATATATGTATAATATAATATATATATATATATTAATGTATTATTCCAATGTGCATGATAAAAGAAAAAAATAATATTTATAAAAAAAAAGAAAAATAAAACAAAAAAAGAAAAAAAAAAAAAAAAAAAAAAAAATACAAAAATAAATAATATAATTTATAATTATATATTCTTGTCACAATAAAAATATATATATATATATATATATTTATAATATGTATATTTTAAACTAGAAAAGGAATAACTAATATTTTATTTATTATCATTCAAGATTTATATTTTATAATAATAAATACCTAATAGAAATATATCAGGATCCATGGCCAAGCCTTTGTCTCAAGAAGAATCCACCCTCATTGAAAGAGCAACGGCTACAATCAACAGCATCCCCATCTCTGAAGACTACAGCGTCGCCAGCGCAGCTCTCTCTAGCGACGGCCGCATCTTCACTGGTGTCAATGTATATCATTTTACTGGGGGACCTTGTGCAGAACTCGTGGTGCTGGGCACTGCTGCTGCTGCGGCAGCTGGCAACCTGACTTGTATCGTCGCGATCGGAAATGAGAACAGGGGCATCTTGAGCCCCTGCGGACGGTGCCGACAGGTGCTTCTCGATCTGCATCCTGGGATCAAAGCGATAGTGAAGGACAGTGATGGACAGCCGACGGCAGTTGGGATTCGTGAATTGCTGCCCTCTGGTTATGTGTGGGAGGGCTAAAAGCTTATTTAATAATAGATTAAAAATATTATAAAAATAAAAACATAAACACAGAAATTACAAAAAAAATACATATGAATTTTTTTTTGTAATCTTCCTTATAAATATAGAATAATGAATCATATAAAACATATCATTATTCATTTATTTACATTTAAAATTATTGTTTCAGTATCTTTAATTTATTATGTATATATAAAAATAACTTACAATTTTATTAATAAACAATATATGTTTATTAATTCATGTTTTGTAATTTATGGGATAGCGATTTTTTTTACTGTCTGTATTTTTCTTTTTTAATTATGTTTTAATTGTATTTTATTTTTATTATTGTTCTTTTTATAGTATTATTTTAAAACAAAATGTATTTTCTAAGAACTTATAATAATAATAAATATAAATTTTAATAAAAATTATATTTATCTTTTACAATATGAACATAAAGTACAACATTAATATATAGCTTTTAATATTTTTATTCCTAATCATGTAAATCTTAAATTTTTCTTTTTAAACATATGTTAAATATTTATTTCTCATTATATATAAGAACATATTTATTAAATCTAGAATTCTATAGTGAGTCGTATTACAATTCACTGGCCGTCGTTTTACAACGTCGTGACTGGGAAAACCCTGGCGTTACCCAACTTAATCGCCTTGCAGCACATCCCCCTTTCGCCAGCTGGCGTAATAGCGAAGAGGCCCGCACCGATCGCCCTTCCCAACAGTTGCGCAGCCTGAATGGCGAATGGCGCCTGATGCGGTATTTTCTCCTTACGCATCTGTGCGGTATTTCACACCGCATATGGTGCACTCTCAGTACAATCTGCTCTGATGCCGCATAGTTAAGCCAGCCCCGACACCCGCCAACACCCGCTGACGCGCCCTGACGGGCTTGTCTGCTCCCGGCATCCGCTTACAGACAAGCTGTGACCGTCTCCGGGAGCTGCATGTGTCAGAGGTTTTCACCGTCATCACCGAAACGCGCGAGACGAAAGGGCCTCGTGATACGCCTATTTTTATAGGTTAATGTCATGATAATAATGGTTTCTTAGACGTCAGGTGGCACTTTTCGGGGAAATGTGCGCGGAACCCCTATTTGTTTATTTTTCTAAATACATTCAAATATGTATCCGCTCATGAGACAATAACCCTGATAAATGCTTCAATAATATTGAAAAAGGAAGAGTATGAGTATTCAACATTTCCGTGTCGCCCTTATTCCCTTTTTTGCGGCATTTTGCCTTCCTGTTTTTGCTCACCCAGAAACGCTGGTGAAAGTAAAAGATGCTGAAGATCAGTTGGGTGCACGAGTGGGTTACATCGAACTGGATCTCAACAGCGGTAAGATCCTTGAGAGTTTTCGCCCCGAAGAACGTTTTCCAATGATGAGCACTTTTAAAGTTCTGCTATGTGGCGCGGTATTATCCCGTATTGACGCCGGGCAAGAGCAACTCGGTCGCCGCATACACTATTCTCAGAATGACTTGGTTGAGTACTCACCAGTCACAGAAAAGCATCTTACGGATGGCATGACAGTAAGAGAATTATGCAGTGCTGCCATAACCATGAGTGATAACACTGCGGCCAACTTACTTCTGACAACGATCGGAGGACCGAAGGAGCTAACCGCTTTTTTGCACAACATGGGGGATCATGTAACTCGCCTTGATCGTTGGGAACCGGAGCTGAATGAAGCCATACCAAACGACGAGCGTGACACCACGATGCCTGTAGCAATGCCAACAACGTTGCGCAAACTATTAACTGGCGAACTACTTACTCTAGCTTCCCGGCAACAATTAATAGACTGGATGGAGGCGGATAAAGTTGCAGGACCACTTCTGCGCTCGGCCCTTCCGGCTGGCTGGTTTATTGCTGATAAATCTGGAGCCGGTGAGCGTGGGTCTCGCGGTATCATTGCAGCACTGGGGCCAGATGGTAAGCCCTCCCGTATCGTAGTTATCTACACGACGGGGAGTCAGGCAACTATGGATGAACGAAATAGACAGATCGCTGAGATAGGTGCCTCACTGATTAAGCATTGGTAACTGTCAGACCAAGTTTACTCATATATACTTTAGATTGATTTAAAACTTCATTTTTAATTTAAAAGGATCTAGGTGAAGATCCTTTTTGATAATCTCATGACCAAAATCCCTTAACGTGAGTTTTCGTTCCACTGAGCGTCAGACCCCGTAGAAAAGATCAAAGGATCTTCTTGAGATCCTTTTTTTCTGCGCGTAATCTGCTGCTTGCAAACAAAAAAACCACCGCTACCAGCGGTGGTTTGTTTGCCGGATCAAGAGCTACCAACTCTTTTTCCGAAGGTAACTGGCTTCAGCAGAGCGCAGATACCAAATACTGTCCTTCTAGTGTAGCCGTAGTTAGGCCACCACTTCAAGAACTCTGTAGCACCGCCTACATACCTCGCTCTGCTAATCCTGTTACCAGTGGCTGCTGCCAGTGGCGATAAGTCGTGTCTTACCGGGTTGGACTCAAGACGATAGTTACCGGATAAGGCGCAGCGGTCGGGCTGAACGGGGGGTTCGTGCACACAGCCCAGCTTGGAGCGAACGACCTACACCGAACTGAGATACCTACAGCGTGAGCTATGAGAAAGCGCCACGCTTCCCGAAGGGAGAAAGGCGGACAGGTATCCGGTAAGCGGCAGGGTCGGAACAGGAGAGCGCACGAGGGAGCTTCCAGGGGGAAACGCCTGGTATCTTTATAGTCCTGTCGGGTTTCGCCACCTCTGACTTGAGCGTCGATTTTTGTGATGCTCGTCAGGGGGGCGGAGCCTATCGAAAAACGCCAGCAACGCGGCCTTTTTACGGTTCCTGGCCTTTTGCTGGCCTTTTGCTCACATGTTCTTTCCTGCGTTATCCCCTGATTCTGTGGATAACCGTATTACCGCCTTTGAGTGAGCTGATACCGCTCGCCGCAGCCGAACGACCGAGCGCAGCGAGTCAGTGAGCGAGGAAGCGGAAGAGCGCCCAATACGCAAACCGCCTCTCCCCGCGCGTTGGCCGATTCATTAATGCAGCTGGCACGACAGGTTTCCCGACTGGAAAGCGGGCAGTGAGCGCAACGCAATTAATGTGAGTTAGCTCACTCATTAGGCACCCCAGGCTTTACACTTTATGCTTCCGGCTCGTATGTTGTGTGGAATTGTGAGCGGATAACAATTTCACACAGGAAACAGCTATGACCATGATTACGCCAAGCTATTTAGGTGACACTATAGAATACTCCGGCCGCTAACGTAACAGACTTAGGAGGAGATCTTAGTAGTTGAGTGATTCTATATACATATACATAAATAAATTACATAATATTATAAATTTTTTTTATATTATATTAGAATGTTTACTATATAAAAATAAATATTTTCTTATATATTTTTTTATTTTTTCATATGAAAAAAGAAATTTTTTATATATTTTTTTATTATATTTTATAAATAAAAAAGAACTTATACCTATTATTATTATTATATATAAAAAATATATATTTTTTTATAAATTCCTATTTTTCCGATTATTTATTTATTTTTTTTTTTTATTGTTTAAAAATATATAAAAAAAATTCTTATATATTTAATATTATAAGTCCATAAATATATATAATTTATATAATTTATATATTTCCATACTTTTTTTATTATATATCATGAGATAAAATAAATTTCATATAGAAATCGTATTTATTTATTAATTTGATATATATTAATAAATAAATATTAATAATAATATGTTATATATATATATATTTATTTAATTATTATATAGGAAACTATATATATATATATATATTATATTTTTTTTTTTGAAATATAATAATATATAATATTATTCCTAAAAATATCTATATTCTTATACCTGAACCCTTTTTTTTTTTTTTTTTTTTTTTTGACTTTCGATTGTTCATTCTGTTTATTGTATAAATATATAAATATATATATATTGTATATTTTATTACATATATTTTTTTTTTTTGAAAGTTAAGAATTTTAACTTAATAAAAAAAGTACATATATTTTTAATAAATGTCCTCCATTATATAAATTGTTATATAAAAGATTTTATATAATTTAAATAGAATTCATTTATAAACAAATTTGTTTATAAAAATATATTTATGTATATATAATATAAATATATATATTTATATATATATATATATATATATTACTATATATATTTTTTTTTTTTTTTCCTTTTTTTTACTTTCCCAAGTTGTACTGCTTCTAAGCTTTTTTAATAAACATATATAATTTGTACAAATATTTTAGATTATATACATGATGTATATTTGAATATATTTTCTATATATTTGTGGTTCCATTTTTGTATATTATATATAATATATTTATATATATATTGATATGTCAATATTTGTATAACACATGAAGTTTTTGTTTTTTTTTTTTTTTTTTTTTAATGGAGAATATTTAATAATATATGAAAAAAATTTTATATAAATATATATATATATATATATATATATATATATATATATATATATATGTATATATATATATTTATATATACATGTATGTTTTTTAAAAAGTTAAATAATTCTATAGATTATTTTCATTGTCTTCACATATATGACATAAATATTTTAAAATCGACATTCCGATATATTATATTTTTAGACTATAATATCCGTTAATAATAAATACACGCAGTCATATTATTTATTATACATTCATTTATTATTTTGTTTTTTTTAATTTCTTACATATAA

pArl1-SBP1-GFP (same backbone as in page 1)

XXXXX = hDHFR

XXXXX = recognition sequences for restriction enzymes

XXXXX = crt promotor

XXXXX = GFP sequence

>pArl1-crt-SBP1-GFP

ggatatggcagcttaatgttcgtttttcttatttatatatttataccaattgattgtatttataactgtaaaaatgtgtatgttgtgtgcatatttttttttgtgcatgcacatgcatgtaaatagctaaaattatgaacattttattttttgttcagaaaaaaaaaactttacacacataaaatggctagtatgaatagccatattttatataaattaaatcctatgaatttatgaccatattaaaaatttagatatttatggaacataatatgtttgaaacaataagacaaaattattattattattattatttttactgttataattatgtgtctccttcaatgattcataaatagttggacttgatttttaaaatgtttataatatgattagcatagttaaataaaaaaagttgaaaaattaaaaaaaaacatataaacacaaatgatggtttttccttcaatttcgatatcaatttatagaaacaaaatatatacttgtataattttatttttttatataaatcattacatatataattatacaatattttttctaagagataattatatattaatatatataaaaaaaggtgttttttttttttttttttatttttatttttattttatggtaatattttattttccttattttataaattatattagtttatatgtgattaattttatatattatcaatttatatatttttaaatgcttacttaattatctttttttttttttttttttttttttcccctctttttatattaatttatttttgaaaaaattgatatatatatatatatataatatatatatatacatgtagtagtattaaacaatgtataatatatataaataatatatttatatatttcatttcaattttaattttttttggttttttttttttttctttttgtcatatttaaaaaaaattatattcatataagttatgcattttttataaacattattcaatatatgtataatataatatatatatatatattaatgtattattccaatgtgcatgataaaagaaaaaaataatatttataaaaaaaaagaaaaataaaacaaaaaaagaaaaaaaaaaaaaaaaaaaaaaaaatacaaaaataaataatataatttataattatatattcttgtcacaataaaaatatatatatatatatatatatttataatatgtatattttaaactagaaaaggaataactaatattttatttattatcattcaagatttatattttataataataaatacctaatagaaatatatcaggatccatgcatggttcgctaaactgcatcgtcgctgtgtcccagaacatgggcatcggcaagaacggggactacccctggccaccgctcaggaacgaatttagatatttccagagaatgaccacaacctcttcagtagaaggtaaacagaatctggtgattatgggtaagaagacctggttctccattcctgagaagaatcgacctttaaagggtagaattaatttagttctcagcagagaactcaaggaacctccacaaggagctcattttctttccagaagtctagatgatgccttaaaacttactgaacaaccagaattagcaaataaagtagacatggtctggatagttggtggcagttctgtttataaggaagccatgaatcacccaggccatcttaaactatttgtgacaaggatcatgcaagactttgaaagtgacacgttttttccagaaattgatttggagaaatataaacttctgccagaatacccaggtgttctctctgatgtccaggaggagaaaggcattaagtacaaatttgaagtatatgagaagaatgattaagcttatttaataatagattaaaaatattataaaaataaaaacataaacacagaaattacaaaaaaaatacatatgaattttttttttgtaatcttccttataaatatagaataatgaatcatataaaacatatcattattcatttatttacatttaaaattattgtttcagtatctttaatttattatgtatatataaaaataacttacaattttattaataaacaatatatgtttattaattcatgttttgtaatttatgggatagcgattttttttactgtctgtatttttcttttttaattatgttttaattgtattttatttttattattgttctttttatagtattattttaaaacaaaatgtattttctaagaacttataataataataaatataaattttaataaaaattatatttatcttttacaatatgaacataaagtacaacattaatatatagcttttaatatttttattcctaatcatgtaaatcttaaatttttctttttaaacatatgttaaatatttatttctcattatatataagaacatatttattaaatctagaattctatagtgagtcgtattacaattcactggccgtcgttttacaacgtcgtgactgggaaaaccctggcgttacccaacttaatcgccttgcagcacatccccctttcgccagctggcgtaatagcgaagaggcccgcaccgatcgcccttcccaacagttgcgcagcctgaatggcgaatggcgcctgatgcggtattttctccttacgcatctgtgcggtatttcacaccgcatatggtgcactctcagtacaatctgctctgatgccgcatagttaagccagccccgacacccgccaacacccgctgacgcgccctgacgggcttgtctgctcccggcatccgcttacagacaagctgtgaccgtctccgggagctgcatgtgtcagaggttttcaccgtcatcaccgaaacgcgcgagacgaaagggcctcgtgatacgcctatttttataggttaatgtcatgataataatggtttcttagacgtcaggtggcacttttcggggaaatgtgcgcggaacccctatttgtttatttttctaaatacattcaaatatgtatccgctcatgagacaataaccctgataaatgcttcaataatattgaaaaaggaagagtatgagtattcaacatttccgtgtcgcccttattcccttttttgcggcattttgccttcctgtttttgctcacccagaaacgctggtgaaagtaaaagatgctgaagatcagttgggtgcacgagtgggttacatcgaactggatctcaacagcggtaagatccttgagagttttcgccccgaagaacgttttccaatgatgagcacttttaaagttctgctatgtggcgcggtattatcccgtattgacgccgggcaagagcaactcggtcgccgcatacactattctcagaatgacttggttgagtactcaccagtcacagaaaagcatcttacggatggcatgacagtaagagaattatgcagtgctgccataaccatgagtgataacactgcggccaacttacttctgacaacgatcggaggaccgaaggagctaaccgcttttttgcacaacatgggggatcatgtaactcgccttgatcgttgggaaccggagctgaatgaagccataccaaacgacgagcgtgacaccacgatgcctgtagcaatgccaacaacgttgcgcaaactattaactggcgaactacttactctagcttcccggcaacaattaatagactggatggaggcggataaagttgcaggaccacttctgcgctcggcccttccggctggctggtttattgctgataaatctggagccggtgagcgtgggtctcgcggtatcattgcagcactggggccagatggtaagccctcccgtatcgtagttatctacacgacggggagtcaggcaactatggatgaacgaaatagacagatcgctgagataggtgcctcactgattaagcattggtaactgtcagaccaagtttactcatatatactttagattgatttaaaacttcatttttaatttaaaaggatctaggtgaagatcctttttgataatctcatgaccaaaatcccttaacgtgagttttcgttccactgagcgtcagaccccgtagaaaagatcaaaggatcttcttgagatcctttttttctgcgcgtaatctgctgcttgcaaacaaaaaaaccaccgctaccagcggtggtttgtttgccggatcaagagctaccaactctttttccgaaggtaactggcttcagcagagcgcagataccaaatactgtccttctagtgtagccgtagttaggccaccacttcaagaactctgtagcaccgcctacatacctcgctctgctaatcctgttaccagtggctgctgccagtggcgataagtcgtgtcttaccgggttggactcaagacgatagttaccggataaggcgcagcggtcgggctgaacggggggttcgtgcacacagcccagcttggagcgaacgacctacaccgaactgagatacctacagcgtgagctatgagaaagcgccacgcttcccgaagggagaaaggcggacaggtatccggtaagcggcagggtcggaacaggagagcgcacgagggagcttccagggggaaacgcctggtatctttatagtcctgtcgggtttcgccacctctgacttgagcgtcgatttttgtgatgctcgtcaggggggcggagcctatcgaaaaacgccagcaacgcggcctttttacggttcctggccttttgctggccttttgctcacatgttctttcctgcgttatcccctgattctgtggataaccgtattaccgcctttgagtgagctgataccgctcgccgcagccgaacgaccgagcgcagcgagtcagtgagcgaggaagcggaagagcgcccaatacgcaaaccgcctctccccgcgcgttggccgattcattaatgcagctggcacgacaggtttcccgactggaaagcgggcagtgagcgcaacgcaattaatgtgagttagctcactcattaggcaccccaggctttacactttatgcttccggctcgtatgttgtgtggaattgtgagcggataacaatttcacacaggaaacagctatgaccatgattacgccaagctatttaggtgacactatagaatactcgcggccgctaacgtaacagacttaggaggagatcttagtagttgagtgattctatatacatatacataaataaattacataatattataaattttttttatattatattagaatgtttactatataaaaataaatattttcttatatatttttttattttttcatatgaaaaaagaaattttttatatatttttttattatattttataaataaaaaagaacttatacctattattattattatatataaaaaatatatatttttttataaattcctatttttccgattatttatttattttttttttttattgtttaaaaatatataaaaaaaattcttatatatttaatattataagtccataaatatatataatttatataatttatatatttccatactttttttattatatatcatgagataaaataaatttcatatagaaatcgtatttatttattaatttgatatatattaataaataaatattaataataatatgttatatatatatatatttatttaattattatataggaaactatatatatatatatatattatatttttttttttgaaatataataatatataatattattcctaaaaatatctatattcttatacctgaacccttttttttttttttttttttttttgactttcgattgttcattctgtttattgtataaatatataaatatatatatattgtatattttattacatatattttttttttttgaaagttaagaattttaacttaataaaaaaagtacatatatttttaataaatgtcctccattatataaattgttatataaaagattttatataatttaaatagaattcatttataaacaaatttgtttataaaaatatatttatgtatatataatataaatatatatatttatatatatatatatatatatattactatatatatttttttttttttttccttttttttactttcccaagttgtactgcttctaagcttttttaataaacatatataatttgtacaaatattttagattatatacatgatgtatatttgaatatattttctatatatttgtggttccatttttgtatattatatataatatatttatatatatattgatatgtcaatatttgtataacacatgaagtttttgtttttttttttttttttttttaatggagaatatttaataatatatgaaaaaaattttatataaatatatatatatatatatatatatatatatatatatatatatatatgtatatatatatatttatatatacatgtatgttttttaaaaagttaaataattctatagattattttcattgtcttcacatatatgacataaatattttaaaatcgacattccgatatattatatttttagactataatatccgttaataataaatacacgcagtcatattatttattatacattcatttattattttgttttttttaattt**cttacatataactcgaccccgggatggtacc**ATGTGTAGCGCAGCTCGAGCATTTGATTTTTTTACTGATTTAGCCGACGAACCAACACAATTACAGGATGCAGTACCAGAGACAACCGAAAAATTGGCCGAAGTAGTTTCGGATGCAGCAACAAATGTTACTGATGCAGTAAGTGATACAGCTAGTGGTATTGGAAGTTTAGTTGGAGAAGCAGCTAGTAGTTTAGGAAATTTAGTTGGTGAAGCAGCAAGCGGTATAGGAAATATAGTTGGAGGTGCAGCAAGCGGTATAGGAAATATAGTTGGAGGTGCAGCAAGCGGTATAGGAAGTTTAGTTGGTGATGCAGCAAGCGGTTTAGGAAATTTAGTTGGTGATGCAGCAGAGGCACTTGCAACTACCGAATTAAAAGATGTAATACCAGAAAATACTGAATCCACAACTGATTTGGTACCATCTGAGGTATCACCTCCAGTAGATGATTATCTCGACGATGACGGTTTTTCAAGCTTTAGAGAATTTCTTGAAAGTACTCCTTGTTGGCAACGTAGAATGGCTCAAGAAGCTTTACTTAATGAATACGAAGTAGAATCTCCAGCCGAATCTATGTCCCCTATTCTTAGAGTACAATTTTTTGCAGATTTTGCAAAACAAGCCGTACATGTTGCTAAACAAAATTATCTCTATGTTGTGATATTCCTATTCTTTGTTATTAACATATTATTGTTCATCAACTTTTACAACTTAGGAAAAAGAAAAGGATATTACCTAGCAAAAAAACAAAAAAAAGAACAAATGCTAGAACAAAACCCAGAACAAAACCCAGAACAAAACGCACAACAAAACGCACAACAAAACGCACAACAAAACGCACAACAAAACGCACAACAAAACGCACAACAAAACGCACAACAAAACACACAACAAAACACACAACAAAAAACACAACAGAACCCACAACAAAACGCACAACAAAACACACAACAAAACACACAACAACAATCCACAACCAAATCCACAACAAAAACAGTTGCTAGAGAAACC**cctaggATGAGTAAAGGAGAAGAACTTTTC**ACTGGAGTTGTCCCAATTCTTGTTGAATTAGATGGTGATGTTAATGGGCACAAATTTTCTGTCAGTGGAGAGGGTGAAGGTGATGCAACATACGGAAAACTTACCCTTAAATTTATTTGCACTACTGGAAAACTACCTGTTCCATGGCCAACACTTGTCACTACTTTCGCGTATGGTCTTCAATGCTTTGCGAGATACCCAGATCATATGAAACAGCATGACTTTTTCAAGAGTGCCATGCCCGAAGGTTATGTACAGGAAAGAACTATATTTTTCAAAGATGACGGGAACTACAAGACACGTGCTGAAGTCAAGTTTGAAGGTGATACCCTTGTTAATAGAATCGAGTTAAAAGGTATTGATTTTAAAGAAGATGGAAACATTCTTGGACACAAATTGGAATACAACTATAACTCACACAATGTATACATCATGGCAGACAAACAAAAGAATGGAATCAAAGTTAACTTCAAAATTAGACACAACATTGAAGATGGAAGCGTTCAACTAGCAGACCATTATCAACAAAATACTCCAATTGGCGATGGCCCTGTCCTTTTACCAGACAACCATTACCTGTCCACACAATCTGCCCTTTCGAAAGATCCCAACGAAAAGAGAGACCACATGGTCCTTCTTGAGTTTGTAACAGCTGCTGGGATTACACATGGCATGGATGAGCTCTACAAAtaactcgag

pArl2 with mal7 promoter and skip-peptide mediated expression of 2 markers (episomal)

XXXXXX = MAL7-promotor

XXXXXX = SBP1

XXXXXX = GFP

XXXXXX = T2A

XXXXXX = MAHRP2

XXXXXX = mScarlet

XXXXXX = yDHODH

>pArl2-mal7-SBP1-GFP-T2A-MAHRP2-mScarlet

ggaattgtgagcggataacaatttcacacaggaaacagctatgaccatgattacgccaagctatttaggtgacactatagaatactcgcggccgctaacgtaacagacttaggaggAGATCTttattattattacatgtagaacgtttggaatatattcatagttttaaaatatatttagaaatatttcttaatatgtttcatacaattaaaatggaattatatataatttatttagaatgaaaagaaaatatacttatataaaaccttatatgttaaaaatatattatatgtatggaaatttatatgatttaattattattattttttttttttataaataaatattttgttagataaccagaatatagtatgtataaaaaaaataattaataattaatttatatggaaaaataagtacttttatatataattaataaagtacattaattatttctttttttttatttatttttttccttttttgtccattatatttttaataagaataattatatatttttagattaaaattataactttgaattaggaagtaaaaattattaaaagatttaaatggtacatattatataatatatatatatatattataattgttatagttgtagatcccttttttttttttttttttaagtttataaaaaaaaatgttataaaagtacataatttatctaccattatttatatactctaggatttattcataaataaataaatactactacttattataaaaaaaatgggacagttttattattatatacaaataaaaatattctgagaaaattcgttcgaaattattattttttttttaatttaaatattagtttctataaaattataattaaaatataaatctcatttttcttttgatttattattaaaaaaaaaaaaaaaaaaattgaaattttattattttatatataatattatatattataaaatataatatatgattaataatatatataaaattctaataaacttggagatatgataaaaaataaaatagaataaacgttttttttttatatatatatattatatatgtagattaaacgtacttacaacggaattatcaaaatatgttataattgaactagtgcataaatatattttaattaatattttttatacatgttaaaaaaaaataaatagtacgactatacatatattttaatttattaatattatatttatatatatatatatatttataatatatttatatatttatataatatatataataattataagaatacttttttctattatttaaaaaagagagaactacatgtaaataaacaataaattatttaaaatatagaaaaattcataggaaacgaaaaaaaaaaaaaagaaaaaaaatctgaccatgtatatataatatatatatataatatatttttatatttattttatatataaatattaaaaagaatgataataatatttgttgtaaatatttttatctctatatttatatataaccttttttttaaaataattatatttttgaaaattattaaaaagttatatttattataatttctagttataaatatttatttttatgaacaaattaataactctttagtatttttaaaaaaaaaataataattagaattttatttttcttaatattttaagaatatatatgtattattataatataatagaggtatttatatttttaaacttataaaaataaaatttttaaaaaaatattatttattttagttgagatataaataatattttttttttattttattttatttaattttaattttttttttttttttaattttattaaaattttattaaCTCGAGATGTGTAGCGCAGCACGAGCATTTGATTTTTTTACCGATTTAGCCGACGAACCAACACAATTACAGGATGCAGTACCAGAGACAACCGAAAAATTGGCCGAAGTAGTTTCGGATGCAGCAACAAATGTTACTGATGCAGTAAGTGATACAGCTAGTGGTATTGGAAGTTTAGTTGGAGAAGCAGCTAGTAGTTTAGGAAATTTAGTTGGTGAAGCAGCAAGCGGTATAGGAAATATAGTTGGAGGTGCAGCAAGCGGTATAGGAAATATAGTTGGAGGTGCAGCAAGCGGTATAGGAAGTTTAGTTGGTGATGCAGCAAGCGGTTTAGGAAATTTAGTTGGTGATGCAGCAGAGGCACTTGCAACTACCGAATTAAAAGATGTAATACCAGAAAATACTGAATCCACAACTGATTTGGTACCATCTGAGGTATCACCTCCAGTAGATGATTATCTCGACGATGACGGTTTTTCAAGCTTTAGAGAATTTCTTGAAAGTACTCCTTGTTGGCAACGTAGAATGGCTCAAGAAGCTTTACTTAATGAATACGAAGTAGAATCTCCAGCCGAATCTATGTCCCCTATTCTTAGAGTACAATTTTTTGCAGATTTTGCAAAACAAGCCGTACATGTTGCTAAACAAAATTATCTCTATGTTGTGATATTCCTATTCTTTGTTATTAACATATTATTGTTCATCAACTTTTACAACTTAGGAAAAAGAAAAGGATATTACCTAGCAAAAAAACAAAAAAAAGAACAAATGCTAGAACAAAACCCAGAACAAAACCCAGAACAAAACGCACAACAAAACGCACAACAAAACGCACAACAAAACGCACAACAAAACGCACAACAAAACGCACAACAAAACGCACAACAAAACACACAACAAAACACACAACAAAAAACACAACAGAACCCACAACAAAACGCACAACAAAACACACAACAAAACACACAACAACAATCCACAACCAAATCCA**CAACAAAAACAGTTGCTAGAGAAACCCCTAGGATGAGTAAAGGAGAAGAACTTTTCACTGGAG**TTGTCCCAATTCTTGTTGAATTAGATGGTGATGTTAATGGGCACAAATTTTCTGTCAGTGGAGAGGGTGAAGGTGATGCAACATACGGAAAACTTACCCTTAAATTTATTTGCACTACTGGAAAACTACCTGTTCCATGGCCAACACTTGTCACTACTTTCGCGTATGGTCTTCAATGCTTTGCGAGATACCCAGATCATATGAAACAGCATGACTTTTTCAAGAGTGCCATGCCCGAAGGTTATGTACAGGAAAGAACTATATTTTTCAAAGATGACGGGAACTACAAGACACGTGCTGAAGTCAAGTTTGAAGGTGATACCCTTGTTAATAGAATCGAGTTAAAAGGTATTGATTTTAAAGAAGATGGAAACATTCTTGGACACAAATTGGAATACAACTATAACTCACACAATGTATACATCATGGCAGACAAACAAAAGAATGGAATCAAAGTTAACTTCAAAATTAGACACAACATTGAAGATGGAAGCGTTCAACTAGCAGACCATTATCAACAAAATACTCCAATTGGCGATGGCCCTGTCCTTTTACCAGACAACCATTACCTGTCCACACAATCTGCCCTTTCGAAAGATCCCAACGAAAAGAGAGACCACATGGTCCTTCTTGAGTTTGTAACAGCTGCTGGGATTACACATGGCATGGATGAACTATACAAAGTCGACGGAGAAGGAAGAGGAA**GTTTATTAACATGTGGAGATGTAGAAGAAAATCCAGGACCAATGCAGCCTTGTCCATATGATGTATAC**AATCAAATAAACCATGTAGGAACTCATTGGGCTCAACATTTAGGAGAACACTTACATCATTTAGCACATATGCATCAACATACTCCACATGTACATCATCACATTCCACATGTGCATACGCTAGCTCATGATAGGCCTTGTATGCCAATAACTGCTTTCTTTTGTCGACATCATGAACATTGTAGTTCCCATTTAATGTTAATCTTTTTATTATTGGCTTTCTTCCTAGTAGTAGTTTATAGATTATATAACGAGgtaagttattattaacatatataagttttttattcaatacatattatatatatatatatattagctgattttattgcattatatattatatatgtacctttattccttttttatttttcagGTTGTTAATTCAGCTAAAACTGTACGTATTGTAAATATAACACCAGTAAATGAGGAACATAAGGCTGAAGCTAGTAAGGAACAATCAAAATCAACAAGTGATTCTTCTACTAGTACACAACAAACATTACCTAGGATGGTGAGTAAGGGTGAGGCAGTGATTAAGGAGTTTATGCGTTTTAAGGTGCACATGGAGGGTAGTATGAACGGTCACGAGTTTGAGATTGAGGGTGAGGGTGAGGGTCGTCCATACGAGGGTACACAGACAGCAAAGCTTAAGGTGACAAAGGGTGGTCCACTTCCATTTAGTTGGGATATTCTTAGTCCACAGTTTATGTACGGTAGTCGTGCATTTACAAAGCACCCAGCAGATATTCCAGATTACTACAAGCAGAGTTTTCCAGAGGGTTTTAAGTGGGAGCGTGTGATGAACTTTGAGGATGGTGGTGCAGTGACAGTGACACAGGATACAAGTCTTGAGGATGGTACACTTATTTACAAGGTGAAGCTTCGTGGTACAAACTTTCCACCAGATGGTCCAGTGATGCAGAAGAAGACAATGGGTTGGGAGGCAAGTACAGAGCGTCTTTACCCAGAGGATGGTGTGCTTAAGGGTGATATTAAGATGGCACTTCGTCTTAAGGATGGTGGTCGTTACCTTGCAGATTTTAAGACAACATACAAGGCAAAGAAGCCAGTGCAGATGCCAGGTGCATACAACGTGGATCGTAAGCTTGATATTACAAGTCACAACGAGGATTACACAGTGGTGGAGCAGTACGAGCGTAGTGAGGGTCGTCACA**GTACAGGTGGTATGGATGAGCTTTACAAGTAACCCGGGtcgagggatatggcagcttaatgttc**gtttttcttatttatatatttataccaattgattgtatttataactgtaaaaatgtgtatgttgtgtgcatatttttttttgtgcatgcacatgcatgtaaatagctaaaattatgaacattttattttttgttcagaaaaaaaaaactttacacacataaaatggctagtatgaatagccatattttatataaattaaatcctatgaatttatgaccatattaaaaatttagatatttatggaacataatatgtttgaaacaataagacaaaattattattattattattatttttactgttataattatgtgtctccttcaatgattcataaatagttggacttgatttttaaaatgtttataatatgattagcatagttaaataaaaaaagttgaaaaattaaaaaaaaacatataaacacaaatgatggtttttccttcaatttcgatatcaatttatagaaacaaaatatatacttgtataattttatttttttatataaatcattacatatataattatacaatattttttctaagagataattatatattaatatatataaaaaaaggtgttttttttttttttttttatttttatttttattttatggtaatattttattttccttattttataaattatattagtttatatgtgattaattttatatattatcaatttatatatttttaaatgcttacttaattatctttttttttttttttttttttttttcccctctttttatattaatttatttttgaaaaaattgatatatatatatatatataatatatatatatacatgtagtagtattaaacaatgtataatatatataaataatatatttatatatttcatttcaattttaattttttttggttttttttttttttctttttgtcatatttaaaaaaaattatattcatataagttatgcattttttataaacattattcaatatatgtataatataatatatatatatatattaatgtattattccaatgtgcatgataaaagaaaaaaataatatttataaaaaaaaagaaaaataaaacaaaaaaagaaaaaaaaaaaaaaaaaaaaaaaaatacaaaaataaataatataatttataattatatattcttgtcacaataaaaatatatatatatatatatatatttataatatgtatattttaaactagaaaaggaataactaatattttatttattatcattcaagatttatattttataataataaatacctaatagaaatatatcaggatccATGACAGCCAGTTTAACTACCAAGTTCTTGAACAATACCTATGAAAACCCATTTATGAATGCATCCGGTGTTCATTGCATGACTACACAAGAATTAGATGAATTAGCAAACTCTAAAGCTGGCGCATTCATTACAAAGAGTGCTACAACCTTAGAAAGAGAAGGTAACCCTGAACCACGTTACATTTCTGTCCCTCTAGGCAGTATCAACTCCATGGGTTTACCAAACGAAGGTATCGACTACTATTTGTCCTATGTATTAAACCGTCAAAAGAATTATCCTGATGCACCTGCTATTTTCTTCTCAGTTGCTGGTATGAGCATTGATGAAAATTTAAATTTGTTGAGGAAAATCCAAGATAGCGAATTCAACGGTATTACCGAGTTAAACTTGTCTTGTCCTAATGTGCCTGGGAAACCACAAGTTGCTTATGACTTTGACTTGACAAAGGAAACCTTGGAAAAGGTTTTTGCCTTTTTCAAAAAACCTCTTGGTGTCAAGTTGCCTCCTTATTTTGATTTTGCCCATTTTGATATCATGGCAAAAATATTGAACGAGTTCCCATTAGCTTATGTCAACTCTATCAATAGTATAGGAAATGGTCTTTTCATTGATGTGGAGAAGGAGAGTGTAGTAGTGAAGCCAAAGAATGGTTTCGGGGGTATTGGAGGTGAATATGTTAAGCCAACCGCGCTCGCCAATGTTCGTGCATTTTACACTCGTTTGAGACCTGAAATCAAAGTTATCGGTACAGGTGGAATTAAGTCCGGTAAGGATGCATTTGAACATCTTCTATGTGGTGCCTCTATGCTACAGATTGGTACAGAATTACAAAAAGAGGGCGTCAAGATTTTTGAACGTATCGAAAAAGAATTAAAAGACATAATGGAAGCTAAGGGTTATACATCCATAGATCAGTTCCGTGGGAAGTTGAACAGCATTTAAaagcttatttaataatagattaaaaatattataaaaataaaaacataaacacagaaattacaaaaaaaatacatatgaattttttttttgtaatcttccttataaatatagaataatgaatcatataaaacatatcattattcatttatttacatttaaaattattgtttcagtatctttaatttattatgtatatataaaaataacttacaattttattaataaacaatatatgtttattaattcatgttttgtaatttatgggatagcgattttttttactgtctgtatttttcttttttaattatgttttaattgtattttatttttattattgttctttttatagtattattttaaaacaaaatgtattttctaagaacttataataataataaatataaattttaataaaaattatatttatcttttacaatatgaacataaagtacaacattaatatatagcttttaatatttttattcctaatcatgtaaatcttaaatttttctttttaaacatatgttaaatatttatttctcattatatataagaacatatttattaaatctagaattctatagtgagtcgtattacaattcactggccgtcgttttacaacgtcgtgactgggaaaaccctggcgttacccaacttaatcgccttgcagcacatccccctttcgccagctggcgtaatagcgaagaggcccgcaccgatcgcccttcccaacagttgcgcagcctgaatggcgaatggcgcctgatgcggtattttctccttacgcatctgtgcggtatttcacaccgcatatggtgcactctcagtacaatctgctctgatgccgcatagttaagccagccccgacacccgccaacacccgctgacgcgccctgacgggcttgtctgctcccggcatccgcttacagacaagctgtgaccgtctccgggagctgcatgtgtcagaggttttcaccgtcatcaccgaaacgcgcgagacgaaagggcctcgtgatacgcctatttttataggttaatgtcatgataataatggtttcttagacgtcaggtggcacttttcggggaaatgtgcgcggaacccctatttgtttatttttctaaatacattcaaatatgtatccgctcatgagacaataaccctgataaatgcttcaataatattgaaaaaggaagagtatgagtattcaacatttccgtgtcgcccttattcccttttttgcggcattttgccttcctgtttttgctcacccagaaacgctggtgaaagtaaaagatgctgaagatcagttgggtgcacgagtgggttacatcgaactggatctcaacagcggtaagatccttgagagttttcgccccgaagaacgttttccaatgatgagcacttttaaagttctgctatgtggcgcggtattatcccgtattgacgccgggcaagagcaactcggtcgccgcatacactattctcagaatgacttggttgagtactcaccagtcacagaaaagcatcttacggatggcatgacagtaagagaattatgcagtgctgccataaccatgagtgataacactgcggccaacttacttctgacaacgatcggaggaccgaaggagctaaccgcttttttgcacaacatgggggatcatgtaactcgccttgatcgttgggaaccggagctgaatgaagccataccaaacgacgagcgtgacaccacgatgcctgtagcaatgccaacaacgttgcgcaaactattaactggcgaactacttactctagcttcccggcaacaattaatagactggatggaggcggataaagttgcaggaccacttctgcgctcggcccttccggctggctggtttattgctgataaatctggagccggtgagcgtgggtctcgcggtatcattgcagcactggggccagatggtaagccctcccgtatcgtagttatctacacgacggggagtcaggcaactatggatgaacgaaatagacagatcgctgagataggtgcctcactgattaagcattggtaactgtcagaccaagtttactcatatatactttagattgatttaaaacttcatttttaatttaaaaggatctaggtgaagatcctttttgataatctcatgaccaaaatcccttaacgtgagttttcgttccactgagcgtcagaccccgtagaaaagatcaaaggatcttcttgagatcctttttttctgcgcgtaatctgctgcttgcaaacaaaaaaaccaccgctaccagcggtggtttgtttgccggatcaagagctaccaactctttttccgaaggtaactggcttcagcagagcgcagataccaaatactgtccttctagtgtagccgtagttaggccaccacttcaagaactctgtagcaccgcctacatacctcgctctgctaatcctgttaccagtggctgctgccagtggcgataagtcgtgtcttaccgggttggactcaagacgatagttaccggataaggcgcagcggtcgggctgaacggggggttcgtgcacacagcccagcttggagcgaacgacctacaccgaactgagatacctacagcgtgagctatgagaaagcgccacgcttcccgaagggagaaaggcggacaggtatccggtaagcggcagggtcggaacaggagagcgcacgagggagcttccagggggaaacgcctggtatctttatagtcctgtcgggtttcgccacctctgacttgagcgtcgatttttgtgatgctcgtcaggggggcggagcctatcgaaaaacgccagcaacgcggcctttttacggttcctggccttttgctggccttttgctcacatgttctttcctgcgttatcccctgattctgtggataaccgtattaccgcctttgagtgagctgataccgctcgccgcagccgaacgaccgagcgcagcgagtcagtgagcgaggaagcggaagagcgcccaatacgcaaaccgcctctccccgcgcgttggccgattcattaatgcagctggcacgacaggtttcccgactggaaagcgggcagtgagcgcaacgcaattaatgtgagttagctcactcattaggcaccccaggctttacactttatgcttccggctcgtatgttgtgt

SLI integration constructs (3xHA) for IT4

XXXXX = hDHFR (WR resistance)

XXXXX = PfIT_060021400 C-terminal targeting sequence

XXXXX = 3xHA

XXXXX = Skip peptide

XXXXX = Neomycin

>pSLI-PfIT_060021400-3xHA-T2A-NeoR

ggatatggcagcttaatgttcgtttttcttatttatatatttataccaattgattgtatttataactgtaaaaatgtgtatgttgtgtgcatatttttttttgtgcatgcacatgcatgtaaatagctaaaattatgaacattttattttttgttcagaaaaaaaaaactttacacacataaaatggctagtatgaatagccatattttatataaattaaatcctatgaatttatgaccatattaaaaatttagatatttatggaacataatatgtttgaaacaataagacaaaattattattattattattatttttactgttataattatgtgtctccttcaatgattcataaatagttggacttgatttttaaaatgtttataatatgattagcatagttaaataaaaaaagttgaaaaattaaaaaaaaacatataaacacaaatgatggtttttccttcaatttcgatatcaatttatagaaacaaaatatatacttgtataattttatttttttatataaatcattacatatataattatacaatattttttctaagagataattatatattaatatatataaaaaaaggtgttttttttttttttttttatttttatttttattttatggtaatattttattttccttattttataaattatattagtttatatgtgattaattttatatattatcaatttatatatttttaaatgcttacttaattatctttttttttttttttttttttttttcccctctttttatattaatttatttttgaaaaaattgatatatatatatatatataatatatatatatacatgtagtagtattaaacaatgtataatatatataaataatatatttatatatttcatttcaattttaattttttttggttttttttttttttctttttgtcatatttaaaaaaaattatattcatataagttatgcattttttataaacattattcaatatatgtataatataatatatatatatatattaatgtattattccaatgtgcatgataaaagaaaaaaataatatttataaaaaaaaagaaaaataaaacaaaaaaagaaaaaaaaaaaaaaaaaaaaaaaaatacaaaaataaataatataatttataattatatattcttgtcacaataaaaatatatatatatatatatatatttataatatgtatattttaaactagaaaaggaataactaatattttatttattatcattcaagatttatattttataataataaatacctaatagaaatatatcaggatccatgcatggttcgctaaactgcatcgtcgctgtgtcccagaacatgggcatcggcaagaacggggactacccctggccaccgctcaggaacgaatttagatatttccagagaatgaccacaacctcttcagtagaaggtaaacagaatctggtgattatgggtaagaagacctggttctccattcctgagaagaatcgacctttaaagggtagaattaatttagttctcagcagagaactcaaggaacctccacaaggagctcattttctttccagaagtctagatgatgccttaaaacttactgaacaaccagaattagcaaataaagtagacatggtctggatagttggtggcagttctgtttataaggaagccatgaatcacccaggccatcttaaactatttgtgacaaggatcatgcaagactttgaaagtgacacgttttttccagaaattgatttggagaaatataaacttctgccagaatacccaggtgttctctctgatgtccaggaggagaaaggcattaagtacaaatttgaagtatatgagaagaatgattaagcttatttaataatagattaaaaatattataaaaataaaaacataaacacagaaattacaaaaaaaatacatatgaattttttttttgtaatcttccttataaatatagaataatgaatcatataaaacatatcattattcatttatttacatttaaaattattgtttcagtatctttaatttattatgtatatataaaaataacttacaattttattaataaacaatatatgtttattaattcatgttttgtaatttatgggatagcgattttttttactgtctgtatttttcttttttaattatgttttaattgtattttatttttattattgttctttttatagtattattttaaaacaaaatgtattttctaagaacttataataataataaatataaattttaataaaaattatatttatcttttacaatatgaacataaagtacaacattaatatatagcttttaatatttttattcctaatcatgtaaatcttaaatttttctttttaaacatatgttaaatatttatttctcattatatataagaacatatttattaaatctagaattctatagtgagtcgtattacaattcactggccgtcgttttacaacgtcgtgactgggaaaaccctggcgttacccaacttaatcgccttgcagcacatccccctttcgccagctggcgtaatagcgaagaggcccgcaccgatcgcccttcccaacagttgcgcagcctgaatggcgaatggcgcctgatgcggtattttctccttacgcatctgtgcggtatttcacaccgcatatggtgcactctcagtacaatctgctctgatgccgcatagttaagccagccccgacacccgccaacacccgctgacgcgccctgacgggcttgtctgctcccggcatccgcttacagacaagctgtgaccgtctccgggagctgcatgtgtcagaggttttcaccgtcatcaccgaaacgcgcgagacgaaagggcctcgtgatacgcctatttttataggttaatgtcatgataataatggtttcttagacgtcaggtggcacttttcggggaaatgtgcgcggaacccctatttgtttatttttctaaatacattcaaatatgtatccgctcatgagacaataaccctgataaatgcttcaataatattgaaaaaggaagagtatgagtattcaacatttccgtgtcgcccttattcccttttttgcggcattttgccttcctgtttttgctcacccagaaacgctggtgaaagtaaaagatgctgaagatcagttgggtgcacgagtgggttacatcgaactggatctcaacagcggtaagatccttgagagttttcgccccgaagaacgttttccaatgatgagcacttttaaagttctgctatgtggcgcggtattatcccgtattgacgccgggcaagagcaactcggtcgccgcatacactattctcagaatgacttggttgagtactcaccagtcacagaaaagcatcttacggatggcatgacagtaagagaattatgcagtgctgccataaccatgagtgataacactgcggccaacttacttctgacaacgatcggaggaccgaaggagctaaccgcttttttgcacaacatgggggatcatgtaactcgccttgatcgttgggaaccggagctgaatgaagccataccaaacgacgagcgtgacaccacgatgcctgtagcaatgccaacaacgttgcgcaaactattaactggcgaactacttactctagcttcccggcaacaattaatagactggatggaggcggataaagttgcaggaccacttctgcgctcggcccttccggctggctggtttattgctgataaatctggagccggtgagcgtgggtctcgcggtatcattgcagcactggggccagatggtaagccctcccgtatcgtagttatctacacgacggggagtcaggcaactatggatgaacgaaatagacagatcgctgagataggtgcctcactgattaagcattggtaactgtcagaccaagtttactcatatatactttagattgatttaaaacttcatttttaatttaaaaggatctaggtgaagatcctttttgataatctcatgaccaaaatcccttaacgtgagttttcgttccactgagcgtcagaccccgtagaaaagatcaaaggatcttcttgagatcctttttttctgcgcgtaatctgctgcttgcaaacaaaaaaaccaccgctaccagcggtggtttgtttgccggatcaagagctaccaactctttttccgaaggtaactggcttcagcagagcgcagataccaaatactgtccttctagtgtagccgtagttaggccaccacttcaagaactctgtagcaccgcctacatacctcgctctgctaatcctgttaccagtggctgctgccagtggcgataagtcgtgtcttaccgggttggactcaagacgatagttaccggataaggcgcagcggtcgggctgaacggggggttcgtgcacacagcccagcttggagcgaacgacctacaccgaactgagatacctacagcgtgagctatgagaaagcgccacgcttcccgaagggagaaaggcggacaggtatccggtaagcggcagggtcggaacaggagagcgcacgagggagcttccagggggaaacgcctggtatctttatagtcctgtcgggtttcgccacctctgacttgagcgtcgatttttgtgatgctcgtcaggggggcggagcctatcgaaaaacgccagcaacgcggcctttttacggttcctggccttttgctggccttttgctcacatgttctttcctgcgttatcccctgattctgtggataaccgtattaccgcctttgagtgagctgataccgctcgccgcagccgaacgaccgagcgcagcgagtcagtgagcgaggaagcggaagagcgcccaatacgcaaaccgcctctccccgcgcgttggccgattcattaatgcagctggcacgacaggtttcccgactggaaagcgggcagtgagcgcaacgcaattaatgtgagttagctcactcattaggcaccccaggctttacactttatgcttccggctcgtatgttgtgtggaattgtgagcggataacaatttcacacaggaaacagctatgaccatgattacgccaagctatttaggtgacactatagaatactc**gcggccgcTAATTGGAGACATATCTTCATCTGATATTACTTCATCAG**AAAGTGAGTATGAAGAATTGGATATCAATGATATATACCCATACAAATCACCTAAATATAAAACGTTGATTGAAGTGGTACTAGAACCATCAAAAAGAGATACAATGAACACGCAAAGTGATATACCATTAAATGATAAACTTGATAGTAATAAACTTACAGATGAAGAATGGAATCAACTGAAACAGGATTTTATTTCAAATATTTCACAAAATTCTCAAATGGATTTACCCAAAAATAATATAAGTGGGAATATTCAAATGGATACCCATCCTCATGTTAATATTTTAGACGATAGTATGCAAGAAAAACCTTTTATTACATCTATTCATGATAGAGATTTACATAATGGTGAAGAAGTTACCTATAATATTAATTTGGATGATCACAAAAATATGAATTTTTCAACTAATCATGATAATATACCACCAAAAAATGATCAAAATGATTTATATACTGGTATAGATTTGATTAATGATTCGATAAGTGGTAACCATAATGTTAATATTTATGATGAATTGTTAAAAAGAAAAGAAAACGAATTATTTGGAACAAATCATACAAAACATACAACAACAAATATTGTTGCCAAACAAACACATAATGACCCTATAGTCAATCAAATAAATTTGTTCCATAAATGGTTAGATAGACATAGAAATATGTGCGAACAGTGGGATAAAAATAAAAAGGAGGAATTGTTAGATAAATTGAATGAAGAATGGAATAAAGAAAATAAAAATAATAGTAATGTCACAGACACAAATGGTGAAAATAATATTACAA**GGGTGTTGAATAGTGATGTTTCTATCCAAATAGATATGAATTCTAAACCTATTGGTACC**ACCATGTACCCATACGATGTTCCAGATTACGCTACGATGTACCCGTACGACGTGCCGGACTACGCGACTATGTATCCATATGATGTTCCAGATTATGCTGTCGACGGAGAAGGAAGAGGAAGTTTATTAACATGTGGAGATGTAGAAGAAAATCCAGGACCAATGATTGAACAAGATGGATTGCACGCAGGTTCTCCGGCCGCTTGGGTGGAGAGGCTATTCGGCTATGACTGGGCACAACAGACAATCGGCTGCTCTGATGCCGCCGTGTTCCGGCTGTCAGCGCAGGGGCGCCCGGTTCTTTTTGTCAAGACCGACCTGTCCGGTGCCCTGAATGAACTGCAGGACGAGGCAGCGCGGCTATCGTGGCTGGCCACGACGGGCGTTCCTTGCGCAGCTGTGCTCGACGTTGTCACTGAAGCGGGAAGGGACTGGCTGCTATTGGGCGAAGTGCCGGGGCAGGATCTCCTGTCATCTCACCTTGCTCCTGCCGAGAAAGTATCCATCATGGCTGATGCAATGCGGCGGCTGCATACGCTTGATCCGGCTACCTGCCCATTCGACCACCAAGCGAAACATCGCATCGAGCGAGCACGTACTCGGATGGAAGCCGGTCTTGTCGATCAGGATGATCTGGACGAAGAGCATCAGGGGCTCGCGCCAGCCGAACTGTTCGCCAGGCTCAAGGCGCGCATGCCCGACGGCGAGGATCTCGTCGTGACCCATGGCGATGCCTGCTTGCCGAATATCATGGTGGAAAATGGCCGCTTTTCTGGATTCATCGACTGTGGCCGGCTGGGTGTGGCGGACCGCTATCAGGACATAGCGTTGGCTACCCGTGATATTGCTGAAGAGCTTGGCGGCGAATGGGCTGACCGCTTCCTCGTGCTTTACGGTATCGCCGCTCCCGATTCGCAGCGCATCGCCTTCTATCGCCTTCTTGACGAGTTCTTCTAACTCGAG

SLI2a integration and TGD constructs (mScarlet)

xxxxx = mScarlet

xxxxx = Skip peptide

xxxxx = yDHODH

xxxxx = BSD-Resistance

>pSLI2a-**Insert**-mScarlet-T2A-yDHODH

**gcggccgcINSERTCCTAGGATGGTGAGTAAGGGTGAGGCAGTGATTAAGG**AGTTTATGCGTTTTAAGGTGCACATGGAGGGTAGTATGAACGGTCACGAGTTTGAGATTGAGGGTGAGGGTGAGGGTCGTCCATACGAGGGTACACAGACAGCAAAGCTTAAGGTGACAAAGGGTGGTCCACTTCCATTTAGTTGGGATATTCTTAGTCCACAGTTTATGTACGGTAGTCGTGCATTTACAAAGCACCCAGCAGATATTCCAGATTACTACAAGCAGAGTTTTCCAGAGGGTTTTAAGTGGGAGCGTGTGATGAACTTTGAGGATGGTGGTGCAGTGACAGTGACACAGGATACAAGTCTTGAGGATGGTACACTTATTTACAAGGTGAAGCTTCGTGGTACAAACTTTCCACCAGATGGTCCAGTGATGCAGAAGAAGACAATGGGTTGGGAGGCAAGTACAGAGCGTCTTTACCCAGAGGATGGTGTGCTTAAGGGTGATATTAAGATGGCACTTCGTCTTAAGGATGGTGGTCGTTACCTTGCAGATTTTAAGACAACATACAAGGCAAAGAAGCCAGTGCAGATGCCAGGTGCATACAACGTGGATCGTAAGCTTGATATTACAAGTCACAACGAGGATTACACAGTGGTGGAGCAGTACGAGCGTAGTGAGGGTCGTCACAGTA**CAGGTGGTATGGATGAGCTTTACAAGGTCGACGGAGAAGGAAGAGGAAGTTTATTAAC**ATGTGGAGATGTAGAAGAAAATCCAGGACCAATGACAGCCAGTTTAACTACCAAGTTCTTGAACAATACCTATGAAAACCCATTTATGAATGCATCCGGTGTTCATTGCATGACTACACAAGAATTAGATGAATTAGCAAACTCTAAAGCTGGCGCATTCATTACAAAGAGTGCTACAACCTTAGAAAGAGAAGGTAACCCTGAACCACGTTACATTTCTGTCCCTCTAGGCAGTATCAACTCCATGGGTTTACCAAACGAAGGTATCGACTACTATTTGTCCTATGTATTAAACCGTCAAAAGAATTATCCTGATGCACCTGCTATTTTCTTCTCAGTTGCTGGTATGAGCATTGATGAAAATTTAAATTTGTTGAGGAAAATCCAAGATAGCGAATTCAACGGTATTACCGAGTTAAACTTGTCTTGTCCTAATGTGCCTGGGAAACCACAAGTTGCTTATGACTTTGACTTGACAAAGGAAACCTTGGAAAAGGTTTTTGCCTTTTTCAAAAAACCTCTTGGTGTCAAGTTGCCTCCTTATTTTGATTTTGCCCATTTTGATATCATGGCAAAAATATTGAACGAGTTCCCATTAGCTTATGTCAACTCTATCAATAGTATAGGAAATGGTCTTTTCATTGATGTGGAGAAGGAGAGTGTAGTAGTGAAGCCAAAGAATGGTTTCGGGGGTATTGGAGGTGAATATGTTAAGCCAACCGCGCTCGCCAATGTTCGTGCATTTTACACTCGTTTGAGACCTGAAATCAAAGTTATCGGTACAGGTGGAATTAAGTCCGGTAAGGATGCATTTGAACATCTTCTATGTGGTGCCTCTATGCTACAGATTGGTACAGAATTACAAAAAGAGGGCGTCAAGATTTTTGAACGTATCGAAAAAGAATTAAAAGACATAATGGAAGCTAAGGGTTATACATCCATAGATCAGTTCCGTGGGAAGTTGAACAGCATTTAACCCGGGtcgagggatatggcagcttaatgttcgtttttcttatttatatatttataccaattgattgtatttataactgtaaaaatgtgtatgttgtgtgcatatttttttttgtgcatgcacatgcatgtaaatagctaaaattatgaacattttattttttgttcagaaaaaaaaaactttacacacataaaatggctagtatgaatagccatattttatataaattaaatcctatgaatttatgaccatattaaaaatttagatatttatggaacataatatgtttgaaacaataagacaaaattattattattattattatttttactgttataattatgtgtctccttcaatgattcataaatagttggacttgatttttaaaatgtttataatatgattagcatagttaaataaaaaaagttgaaaaattaaaaaaaaacatataaacacaaatgatggtttttccttcaatttcgatatcaatttatagaaacaaaatatatacttgtataattttatttttttatataaatcattacatatataattatacaatattttttctaagagataattatatattaatatatataaaaaaaggtgttttttttttttttttttatttttatttttattttatggtaatattttattttccttattttataaattatattagtttatatgtgattaattttatatattatcaatttatatatttttaaatgcttacttaattatctttttttttttttttttttttttttcccctctttttatattaatttatttttgaaaaaattgatatatatatatatatataatatatatatatacatgtagtagtattaaacaatgtataatatatataaataatatatttatatatttcatttcaattttaattttttttggttttttttttttttctttttgtcatatttaaaaaaaattatattcatataagttatgcattttttataaacattattcaatatatgtataatataatatatatatatatattaatgtattattccaatgtgcatgataaaagaaaaaaataatatttataaaaaaaaagaaaaataaaacaaaaaaagaaaaaaaaaaaaaaaaaaaaaaaaatacaaaaataaataatataatttataattatatattcttgtcacaataaaaatatatatatatatatatatatttataatatgtatattttaaactagaaaaggaataactaatattttatttattatcattcaagatttatattttataataataaatacctaatagaaatatatcaGGATCCATGGCCAAGCCTTTGTCTCAAGAAGAATCCACCCTCATTGAAAGAGCAACGGCTACAATCAACAGCATCCCCATCTCTGAAGACTACAGCGTCGCCAGCGCAGCTCTCTCTAGCGACGGCCGCATCTTCACTGGTGTCAATGTATATCATTTTACTGGGGGACCTTGTGCAGAACTCGTGGTGCTGGGCACTGCTGCTGCTGCGGCAGCTGGCAACCTGACTTGTATCGTCGCGATCGGAAATGAGAACAGGGGCATCTTGAGCCCCTGCGGACGGTGCCGACAGGTGCTTCTCGATCTGCATCCTGGGATCAAAGCGATAGTGAAGGACAGTGATGGACAGCCGACGGCAGTTGGGATTCGTGAATTGCTGCCCTCTGGTTATGTGTGGGAGGGCAAGCTTatttaataatagattaaaaatattataaaaataaaaacataaacacagaaattacaaaaaaaatacatatgaattttttttttgtaatcttccttataaatatagaataatgaatcatataaaacatatcattattcatttatttacatttaaaattattgtttcagtatctttaatttattatgtatatataaaaataacttacaattttattaataaacaatatatgtttattaattcatgttttgtaatttatgggatagcgattttttttactgtctgtatttttcttttttaattatgttttaattgtattttatttttattattgttctttttatagtattattttaaaacaaaatgtattttctaagaacttataataataataaatataaattttaataaaaattatatttatcttttacaatatgaacataaagtacaacattaatatatagcttttaatatttttattcctaatcatgtaaatcttaaatttttctttttaaacatatgttaaatatttatttctcattatatataagaacatatttattaaatctagaattctatagtgagtcgtattacaattcactggccgtcgttttacaacgtcgtgactgggaaaaccctggcgttacccaacttaatcgccttgcagcacatccccctttcgccagctggcgtaatagcgaagaggcccgcaccgatcgcccttcccaacagttgcgcagcctgaatggcgaatggcgcctgatgcggtattttctccttacgcatctgtgcggtatttcacaccgcatatggtgcactctcagtacaatctgctctgatgccgcatagttaagccagccccgacacccgccaacacccgctgacgcgccctgacgggcttgtctgctcccggcatccgcttacagacaagctgtgaccgtctccgggagctgcatgtgtcagaggttttcaccgtcatcaccgaaacgcgcgagacgaaagggcctcgtgatacgcctatttttataggttaatgtcatgataataatggtttcttagacgtcaggtggcacttttcggggaaatgtgcgcggaacccctatttgtttatttttctaaatacattcaaatatgtatccgctcatgagacaataaccctgataaatgcttcaataatattgaaaaaggaagagtatgagtattcaacatttccgtgtcgcccttattcccttttttgcggcattttgccttcctgtttttgctcacccagaaacgctggtgaaagtaaaagatgctgaagatcagttgggtgcacgagtgggttacatcgaactggatctcaacagcggtaagatccttgagagttttcgccccgaagaacgttttccaatgatgagcacttttaaagttctgctatgtggcgcggtattatcccgtattgacgccgggcaagagcaactcggtcgccgcatacactattctcagaatgacttggttgagtactcaccagtcacagaaaagcatcttacggatggcatgacagtaagagaattatgcagtgctgccataaccatgagtgataacactgcggccaacttacttctgacaacgatcggaggaccgaaggagctaaccgcttttttgcacaacatgggggatcatgtaactcgccttgatcgttgggaaccggagctgaatgaagccataccaaacgacgagcgtgacaccacgatgcctgtagcaatgccaacaacgttgcgcaaactattaactggcgaactacttactctagcttcccggcaacaattaatagactggatggaggcggataaagttgcaggaccacttctgcgctcggcccttccggctggctggtttattgctgataaatctggagccggtgagcgtgggtctcgcggtatcattgcagcactggggccagatggtaagccctcccgtatcgtagttatctacacgacggggagtcaggcaactatggatgaacgaaatagacagatcgctgagataggtgcctcactgattaagcattggtaactgtcagaccaagtttactcatatatactttagattgatttaaaacttcatttttaatttaaaaggatctaggtgaagatcctttttgataatctcatgaccaaaatcccttaacgtgagttttcgttccactgagcgtcagaccccgtagaaaagatcaaaggatcttcttgagatcctttttttctgcgcgtaatctgctgcttgcaaacaaaaaaaccaccgctaccagcggtggtttgtttgccggatcaagagctaccaactctttttccgaaggtaactggcttcagcagagcgcagataccaaatactgtccttctagtgtagccgtagttaggccaccacttcaagaactctgtagcaccgcctacatacctcgctctgctaatcctgttaccagtggctgctgccagtggcgataagtcgtgtcttaccgggttggactcaagacgatagttaccggataaggcgcagcggtcgggctgaacggggggttcgtgcacacagcccagcttggagcgaacgacctacaccgaactgagatacctacagcgtgagctatgagaaagcgccacgcttcccgaagggagaaaggcggacaggtatccggtaagcggcagggtcggaacaggagagcgcacgagggagcttccagggggaaacgcctggtatctttatagtcctgtcgggtttcgccacctctgacttgagcgtcgatttttgtgatgctcgtcaggggggcggagcctatcgaaaaacgccagcaacgcggcctttttacggttcctggccttttgctggccttttgctcacatgttctttcctgcgttatcccctgattctgtggataaccgtattaccgcctttgagtgagctgataccgctcgccgcagccgaacgaccgagcgcagcgagtcagtgagcgaggaagcggaagagcgcccaatacgcaaaccgcctctccccgcgcgttggccgattcattaatgcagctggcacgacaggtttcccgactggaaagcgggcagtgagcgcaacgcaattaatgtgagttagctcactcattaggcaccccaggctttacactttatgcttccggctcgtatgttgtgtggaattgtgagcggataacaatttcacacaggaaacagctatgaccatgattacgccaag**ctatttaggtgacactatagaatactc**

**Inserts used:**

>MAHRP2

**taaCAGCCTTGTCCATATGATGTATACAATC**AAATAAACCATGTAGGAACTCATTGGGCTCAACATTTAGGAGAACACTTACATCATTTAGCACATATGCATCAACATACTCCACATGTACATCATCACATTCCACATGTGCATACGCTAGCTCATGATAGGCCTTGTATGCCAATAACTGCTTTCTTTTGTCGACATCATGAACATTGTAGTTCCCATTTAATGTTAATCTTTTTATTATTGGCTTTCTTCCTAGTAGTAGTTTATAGATTATATAACGAGgtaagttattattaacatatataagttttttattcaatacatattatatatatatatatattagctgattttattgcattatatattatatatgtacctttattccttttttatttttcagGTTGTTAATTCAGCTAAAACTGTACGTATTGTAAATATAACACCAGTAAATGAGGAACATAAGGCTGAAGCTAGTAAGGAACAATCAAAATCAACAAGT**GATTCTTCTACTAGTACACAACAAACATTA**

>MSRP6

**taaGATGATGAAGATGATGAAGATGATGAAGATGATG**TAGATGATGAAGATGATCTAGATGTTGAAGATAATGTAGATGATGAATATGATGATGAGCATAATCATAATTATAATGATGACAAATTGAGTGAAAATCCTGAGAAGTATTCAAATTATAATAAAAATATACACGAAGATAAGAAAAAAGATAATTTGAATGAACCACATTTTAAACAAACTCATTATATATATTCATCAAATCACGATAATAATGAAACATCTAGATTTCCAAAAAAAAATGTACCTAAATATGATGAAAAATTAAATAATGAATTTAAAACATATTTAAGAAGACCTGAAAAGAAAGAAGAAACAAAAGAATATCCTAACAATGGATGTTCTGTAGTTCAAATATCTATAGTAACTAATGAAGATTTTTTAAAAAAAGTTAGAGAAAGAAATAAAAAGAGAAATAATAAAAAAAGAACAAATATTTATGATAGTGATGAAGAATCAGAAAGTTCAGAAGAAACAAGTAAAGATCCCTATTCATCGGGTCCATATACAGTTGATCATAAAAATGAAAATTGTTTTCCTTACAAGCAAAATTCAGATATTCATCCAAATATAAATAATGATAATAATGATAATAATGAAAATAATGATAATAATGAAAATAATGATAATAATATACCTGTTAATAATAAAAAAAATTCTCATACTCCTCATGTACCAAAAGATACCAAAAATAATATAGAAAAAAATCATAGTAACTATAATTACAAACCAATTAATCAAGAAAATTTATCAATGTATTTTAATAATAATAATAAGAAGAAGAATAAAAAGAATAGTCAAAAAAATAATCAAAAGAATAATCAAAAGAATAATCAAAAGCATAACCAAGATCTTTATAATATAAAAAAAAATAATGATCTACCTCAATATGAAGAATCTAATAAAACTTATAAAAAAATTTTTTCAAATGAACATGAACCACGTCCGATGATAACAACATCCAGATATGAAAATAAAATATCAAATGATAATAACAACAATAACTATAATTTGAATGAACAAGAAAGAAAACTTATTAAAAATATGATAGATATATTTATTTATACATACAAGTTAAATTATACAAACAGTAAATCTATATCTAAATTATTTAAAAATAATTTACTTAAAAAAAACTTCCGTACTTATTTCACAAATTATATATATACACTTTTTAATTATGGAAAGACATATAATTTTCTAACACCTTATAATAAAGATAATGATCATATGTATAGACAATTATTTGATGAAGCTGTACAAATGATGGATCTA**CTTATTAATAAAATGGACTTAGCTTTAAATCCCACGAAATTA**

>ΔPf332

**taaTCTAATATAAATAACAAAGACTCTAGTACAG**AATGGAATTGTAAAGAAGATGTGGGATGTGTTCCACCTAGGAGACAGAATTTGAATATGGAAAGGTTGGATAATGAAAATGAAGATTCTGTACCCGATTTCATGAAGAAAACTTTTTATCTTGCTGCTGCTGGAGAAGGAAAGAAGTTACGTGAGAAGCATGATGAGAGTTGTGATGAATTCTGTGACGCATGGAATAGAAGTTTAGCTGATTATAAAGATATATTTCAAGGAAAGGATATGTGGAATGATGGGAAATATGGTGAAGCGAAAAATCATATTAAGAATGCTTTTGGTGATATGAACAATAGAAAAACTATGTTAAATGAAATTGAGAAAGGAATTAAAGATGAAACGTTTAGTCGTGAAAATGGTTTAGACGTTTGTAAATCTCAATGTGAGGAAAGAAGTAGAGATGACACAGAAGATCAATTTTTGAGGTTTTTT**GCAGAATGGGAAGAAGAATTTTGTGATGGG**

>ΔPIESP2 (BsiWI instead of AvrII after insert)

taaTTACTCTTTTTTGCAAAACTTGTCGTATTTACCTTTTTCTTTTGGCTTTTAAAATATGGGAAAACGgtaagacgtacatgatattcagaataatatatatatatatatatatgtatgtttttatagtacaattaatacgccttttataacgcttataaagaataaaaatctttatatatatatatatatatatatatttgtatgtatttataatttacctttaatcattgttatattaattttttattttttattttctttcagAGGTCATATCCCAAATCTGGCCATAAGGGACATACGAAATTAAATCAACCAGTAGTTAGAACATTAGCAGATTTTAATGACATGTTTGCAAACCAAAAAAATACATTTAATTTTCTAAAACATATAAATCATTATAAAAATGAACAAGATACAAATAATACACACACGCCAAATCATGATGAATATTCT

>ΔMSRP6 (BsiWI instead of AvrII after insert)

taaTCTTGCTTATTTTTAATATTTTTAAGTGTAATATTTTGTAGTGAACCAGATACAAATTCATTTGATGAAAATGTAAAGAAGAATGAAGTTTTTAATGCCTTAAATGAACATTTAGAAAGTATAAGTAATATCGTAAAAGTAAATATTATGGATGCCCTTTCAAATAACCCTTCGTTAATAAAAAAGACATACGAAGCTGTTGAAATAAATGATGATGATTATGTTCTGGAATATGTTGGGGATTCGACAGGAAAATATGGAGAGGGCTCTATATTCTATGATGAAAGTAAGAAATATAATTATAGAAAGTATTTAGATATAGAAAATGAATTACGAAATATGAAAGGTGAAGATGATGATGATTTTGATGAAGATGATGATGATTTTGATGAAGATGATGAA

SLI2a-TGD constructs (GFP)

xxxxx = SBP1 N-terminal targeting sequence

xxxxx = GFP

xxxxx = Skip peptide

xxxxx = yDHODH

xxxxx = BSD-Resistance

>pSLI2a-TGD-SBP1-GFP-T2A-yDHODH

**gcggccgctaa**TGTAGCGCAGCTCGAGCATTTGATTTTTTTACTGATTTAGCCGACGAACCAACACAATTACAGGATGCAGTACCAGAGACAACCGAAAAATTGGCCGAAGTAGTTTCGGATGCAGCAACAAATGTTACTGATGCAGTAAGTGATACAGCTAGTGGTATTGGAAGTTTAGTTGGAGAAGCAGCTAGTAGTTTAGGAAATTTAGTTGGTGAAGCAGCAAGCGGTATAGGAAATATAGTTGGAGGTGCAGCAAGCGGTATAGGAAATATAGTTGGAGGTGCAGCAAGCGGTATAGGAAGTTTAGTTGGTGATGCAGCAAGCGGTTTAGGAAATTTAGTTGGTGATGCAGCAGAGGCACTTGCAACTACCGAATTAAAAGATGTAATACCAGAAAATACTGAATCCACAACTGATTTGGTACCATCTGAGGTATCACCTCCAGTAGATGATTATCTCGACGATGACGGTTTTTCAAGCTTTAGAGAATTTCTTGAAAGTACTCCTTGTTGGCAACGTAGAATGGCTCAAGAAGCTTTACTTAATGAATACGAAGTAGAATCTCCAGCCGAATCTATGTCCCCTATTCTTAGAGTACAATTTTTT**CGTACG**ATGAGTAAAGGAGAAGAACTTTTCACTGGAGTTGTCCCAATTCTTGTTGAATTAGATGGTGATGTTAATGGGCACAAATTTTCTGTCAGTGGAGAGGGTGAAGGTGATGCAACATACGGAAAACTTACCCTTAAATTTATTTGCACTACTGGAAAACTACCTGTTCCATGGCCAACACTTGTCACTACTTTCGCGTATGGTCTTCAATGCTTTGCGAGATACCCAGATCATATGAAACAGCATGACTTTTTCAAGAGTGCCATGCCCGAAGGTTATGTACAGGAAAGAACTATATTTTTCAAAGATGACGGGAACTACAAGACACGTGCTGAAGTCAAGTTTGAAGGTGATACCCTTGTTAATAGAATCGAGTTAAAAGGTATTGATTTTAAAGAAGATGGAAACATTCTTGGACACAAATTGGAATACAACTATAACTCACACAATGTATACATCATGGCAGACAAACAAAAGAATGGAATCAAAGTTAACTTCAAAATTAGACACAACATTGAAGATGGAAGCGTTCAACTAGCAGACCATTATCAACAAAATACTCCAATTGGCGATGGCCCTGTCCTTTTACCAGACAACCATTACCTGTCCACACAATCTGCCCTTTCGAAAGATCCCAACGAAAAGAGAGACCACATGGTCCTTCTTGAGTTTGTAACAGCTGCTGGGATTACACATGGCATGGATGAGCTCTACAAAGTCGACGGAGAAGGAAGAGGAAGTTTATTAACATGTGGAGATGTAGAAGAAAATCCAGGACCAATGACAGCCAGTTTAACTACCAAGTTCTTGAACAATACCTATGAAAACCCATTTATGAATGCATCCGGTGTTCATTGCATGACTACACAAGAATTAGATGAATTAGCAAACTCTAAAGCTGGCGCATTCATTACAAAGAGTGCTACAACCTTAGAAAGAGAAGGTAACCCTGAACCACGTTACATTTCTGTCCCTCTAGGCAGTATCAACTCCATGGGTTTACCAAACGAAGGTATCGACTACTATTTGTCCTATGTATTAAACCGTCAAAAGAATTATCCTGATGCACCTGCTATTTTCTTCTCAGTTGCTGGTATGAGCATTGATGAAAATTTAAATTTGTTGAGGAAAATCCAAGATAGCGAATTCAACGGTATTACCGAGTTAAACTTGTCTTGTCCTAATGTGCCTGGGAAACCACAAGTTGCTTATGACTTTGACTTGACAAAGGAAACCTTGGAAAAGGTTTTTGCCTTTTTCAAAAAACCTCTTGGTGTCAAGTTGCCTCCTTATTTTGATTTTGCCCATTTTGATATCATGGCAAAAATATTGAACGAGTTCCCATTAGCTTATGTCAACTCTATCAATAGTATAGGAAATGGTCTTTTCATTGATGTGGAGAAGGAGAGTGTAGTAGTGAAGCCAAAGAATGGTTTCGGGGGTATTGGAGGTGAATATGTTAAGCCAACCGCGCTCGCCAATGTTCGTGCATTTTACACTCGTTTGAGACCTGAAATCAAAGTTATCGGTACAGGTGGAATTAAGTCCGGTAAGGATGCATTTGAACATCTTCTATGTGGTGCCTCTATGCTACAGATTGGTACAGAATTACAAAAAGAGGGCGTCAAGATTTTTGAACGTATCGAAAAAGAATTAAAAGACATAATGGAAGCTAAGGGTTATACATCCATAGATCAGTTCCGTGGGAAGTTGAACAGCATTTAACCCGGGtcgagggatatggcagcttaatgttcgtttttcttatttatatatttataccaattgattgtatttataactgtaaaaatgtgtatgttgtgtgcatatttttttttgtgcatgcacatgcatgtaaatagctaaaattatgaacattttattttttgttcagaaaaaaaaaactttacacacataaaatggctagtatgaatagccatattttatataaattaaatcctatgaatttatgaccatattaaaaatttagatatttatggaacataatatgtttgaaacaataagacaaaattattattattattattatttttactgttataattatgtgtctccttcaatgattcataaatagttggacttgatttttaaaatgtttataatatgattagcatagttaaataaaaaaagttgaaaaattaaaaaaaaacatataaacacaaatgatggtttttccttcaatttcgatatcaatttatagaaacaaaatatatacttgtataattttatttttttatataaatcattacatatataattatacaatattttttctaagagataattatatattaatatatataaaaaaaggtgttttttttttttttttttatttttatttttattttatggtaatattttattttccttattttataaattatattagtttatatgtgattaattttatatattatcaatttatatatttttaaatgcttacttaattatctttttttttttttttttttttttttcccctctttttatattaatttatttttgaaaaaattgatatatatatatatatataatatatatatatacatgtagtagtattaaacaatgtataatatatataaataatatatttatatatttcatttcaattttaattttttttggttttttttttttttctttttgtcatatttaaaaaaaattatattcatataagttatgcattttttataaacattattcaatatatgtataatataatatatatatatatattaatgtattattccaatgtgcatgataaaagaaaaaaataatatttataaaaaaaaagaaaaataaaacaaaaaaagaaaaaaaaaaaaaaaaaaaaaaaaatacaaaaataaataatataatttataattatatattcttgtcacaataaaaatatatatatatatatatatatttataatatgtatattttaaactagaaaaggaataactaatattttatttattatcattcaagatttatattttataataataaatacctaatagaaatatatcaGGATCCATGGCCAAGCCTTTGTCTCAAGAAGAATCCACCCTCATTGAAAGAGCAACGGCTACAATCAACAGCATCCCCATCTCTGAAGACTACAGCGTCGCCAGCGCAGCTCTCTCTAGCGACGGCCGCATCTTCACTGGTGTCAATGTATATCATTTTACTGGGGGACCTTGTGCAGAACTCGTGGTGCTGGGCACTGCTGCTGCTGCGGCAGCTGGCAACCTGACTTGTATCGTCGCGATCGGAAATGAGAACAGGGGCATCTTGAGCCCCTGCGGACGGTGCCGACAGGTGCTTCTCGATCTGCATCCTGGGATCAAAGCGATAGTGAAGGACAGTGATGGACAGCCGACGGCAGTTGGGATTCGTGAATTGCTGCCCTCTGGTTATGTGTGGGAGGGCAAGCTTatttaataatagattaaaaatattataaaaataaaaacataaacacagaaattacaaaaaaaatacatatgaattttttttttgtaatcttccttataaatatagaataatgaatcatataaaacatatcattattcatttatttacatttaaaattattgtttcagtatctttaatttattatgtatatataaaaataacttacaattttattaataaacaatatatgtttattaattcatgttttgtaatttatgggatagcgattttttttactgtctgtatttttcttttttaattatgttttaattgtattttatttttattattgttctttttatagtattattttaaaacaaaatgtattttctaagaacttataataataataaatataaattttaataaaaattatatttatcttttacaatatgaacataaagtacaacattaatatatagcttttaatatttttattcctaatcatgtaaatcttaaatttttctttttaaacatatgttaaatatttatttctcattatatataagaacatatttattaaatctagaattctatagtgagtcgtattacaattcactggccgtcgttttacaacgtcgtgactgggaaaaccctggcgttacccaacttaatcgccttgcagcacatccccctttcgccagctggcgtaatagcgaagaggcccgcaccgatcgcccttcccaacagttgcgcagcctgaatggcgaatggcgcctgatgcggtattttctccttacgcatctgtgcggtatttcacaccgcatatggtgcactctcagtacaatctgctctgatgccgcatagttaagccagccccgacacccgccaacacccgctgacgcgccctgacgggcttgtctgctcccggcatccgcttacagacaagctgtgaccgtctccgggagctgcatgtgtcagaggttttcaccgtcatcaccgaaacgcgcgagacgaaagggcctcgtgatacgcctatttttataggttaatgtcatgataataatggtttcttagacgtcaggtggcacttttcggggaaatgtgcgcggaacccctatttgtttatttttctaaatacattcaaatatgtatccgctcatgagacaataaccctgataaatgcttcaataatattgaaaaaggaagagtatgagtattcaacatttccgtgtcgcccttattcccttttttgcggcattttgccttcctgtttttgctcacccagaaacgctggtgaaagtaaaagatgctgaagatcagttgggtgcacgagtgggttacatcgaactggatctcaacagcggtaagatccttgagagttttcgccccgaagaacgttttccaatgatgagcacttttaaagttctgctatgtggcgcggtattatcccgtattgacgccgggcaagagcaactcggtcgccgcatacactattctcagaatgacttggttgagtactcaccagtcacagaaaagcatcttacggatggcatgacagtaagagaattatgcagtgctgccataaccatgagtgataacactgcggccaacttacttctgacaacgatcggaggaccgaaggagctaaccgcttttttgcacaacatgggggatcatgtaactcgccttgatcgttgggaaccggagctgaatgaagccataccaaacgacgagcgtgacaccacgatgcctgtagcaatgccaacaacgttgcgcaaactattaactggcgaactacttactctagcttcccggcaacaattaatagactggatggaggcggataaagttgcaggaccacttctgcgctcggcccttccggctggctggtttattgctgataaatctggagccggtgagcgtgggtctcgcggtatcattgcagcactggggccagatggtaagccctcccgtatcgtagttatctacacgacggggagtcaggcaactatggatgaacgaaatagacagatcgctgagataggtgcctcactgattaagcattggtaactgtcagaccaagtttactcatatatactttagattgatttaaaacttcatttttaatttaaaaggatctaggtgaagatcctttttgataatctcatgaccaaaatcccttaacgtgagttttcgttccactgagcgtcagaccccgtagaaaagatcaaaggatcttcttgagatcctttttttctgcgcgtaatctgctgcttgcaaacaaaaaaaccaccgctaccagcggtggtttgtttgccggatcaagagctaccaactctttttccgaaggtaactggcttcagcagagcgcagataccaaatactgtccttctagtgtagccgtagttaggccaccacttcaagaactctgtagcaccgcctacatacctcgctctgctaatcctgttaccagtggctgctgccagtggcgataagtcgtgtcttaccgggttggactcaagacgatagttaccggataaggcgcagcggtcgggctgaacggggggttcgtgcacacagcccagcttggagcgaacgacctacaccgaactgagatacctacagcgtgagctatgagaaagcgccacgcttcccgaagggagaaaggcggacaggtatccggtaagcggcagggtcggaacaggagagcgcacgagggagcttccagggggaaacgcctggtatctttatagtcctgtcgggtttcgccacctctgacttgagcgtcgatttttgtgatgctcgtcaggggggcggagcctatcgaaaaacgccagcaacgcggcctttttacggttcctggccttttgctggccttttgctcacatgttctttcctgcgttatcccctgattctgtggataaccgtattaccgcctttgagtgagctgataccgctcgccgcagccgaacgaccgagcgcagcgagtcagtgagcgaggaagcggaagagcgcccaatacgcaaaccgcctctccccgcgcgttggccgattcattaatgcagctggcacgacaggtttcccgactggaaagcgggcagtgagcgcaacgcaattaatgtgagttagctcactcattaggcaccccaggctttacactttatgcttccggctcgtatgttgtgtggaattgtgagcggataacaatttcacacaggaaacagctatgaccatgattacgccaagctatttaggtgacactatagaatactc
